# Integrated structure-function dataset reveals key mechanisms underlying photochromic fluorescent proteins

**DOI:** 10.1101/2020.09.25.313528

**Authors:** Elke De Zitter, Siewert Hugelier, Sam Duwé, Wim Vandenberg, Alison G. Tebo, Luc Van Meervelt, Peter Dedecker

## Abstract

Photochromic fluorescent proteins have become versatile tools in the life sciences, though our understanding of their structure-function relation is limited. Starting from a single scaffold, we have developed a range of 27 photochromic fluorescent proteins that cover a broad range of spectroscopic properties, yet differ only in one or two mutations. We also determined 43 different crystal structures of these mutants. Correlation and principal component analysis of the spectroscopic and structural properties confirmed the complex relationship between structure and spectroscopy, suggesting that the observed variability does not arise from a limited number of mechanisms, but also allowed us to identify consistent trends and to relate these to the spatial organization around the chromophore. We find that particular changes in spectroscopic properties can come about through multiple different underlying mechanisms, of which the polarity of the chromophore environment and hydrogen bonding of the chromophore are key modulators. Furthermore, some spectroscopic parameters, such as the photochromism, appear to be largely determined by a single or a few structural properties, while other parameters, such as the absorption maximum, do not allow a clear identification of a single cause. We also highlight the role of water molecules close to the chromophore in influencing photochromism. We anticipate that our dataset can open opportunities for the development and evaluation of new and existing protein engineering methods.

## Introduction

Photochromic fluorescent proteins (RSFPs) are fluorophores that can reversibly switch between a fluorescent and a non-fluorescent dark state upon irradiation with light of an appropriate wavelength. Many forms of advanced fluorescence imaging, such as sub-diffraction fluorescence imaging or dynamic labelling, rely on the availability of these smart labels, where the achievable performance and the overall image quality is crucially determined by the label’s properties [1].

Over the years, several families of RSFPs have been obtained from a number of different organisms, and have been the subject of several spectroscopic, structural and theoretical studies.[2–10] These studies have revealed a rich photochemistry, involving multiple spectroscopic states and associated interconversions. Their complex photochemistry is a direct consequence of their intricate structure-function relationship, in which the probe properties are determined not just by the structure of the chromophore, but also by the chromophore’s interactions with the surrounding amino acids. Mutating just a single residue in the vicinity of the chromophore can easily lead to profound changes in the spectroscopic properties [11–13].

The many applications of these types of ‘smart labels’ has resulted in extensive efforts to engineer variants with properties tailored to particular applications. This has turned out to be challenging, since the complex structure-function relationship of these proteins makes it difficult to apply rational development strategies. Successful application in e.g. advanced imaging also tends to require that systems perform well across multiple different parameters, which is considerably more complex than optimizing for just a single property [14], especially if the extent of correlations between the different parameters is unknown. Nevertheless, considerable progress has been made in our understanding of the structure-function relation of fluorescent proteins, including their photochromism. For example, it is known that photochromism typically involves a chromophore *cis/trans* isomerization [15] and (de-)protonation [2,16,17], and requires an environment capable of both allowing this isomerization and of stabilizing the resulting states. [12, 14, 18] However, achieving a more quantitative understanding is complicated by the fact that most studies investigate one or only a few mutants, possibly focusing on only a few spectroscopic parameters, and do so for different scaffolds arising from all over the diverse FP family.

The limited availability of structure/function data of RSFPs not only complicates our understanding of their operating mechanisms, but also limits efforts to apply innovative new methodologies, such as machine learning, for RSFP engineering. Likewise, fluorescent proteins represent exciting targets for the development and validation of new approaches in computational modelling [19], on account of their well-defined and rigid structure combined with the high sensitivity of the spectroscopic properties to seemingly minor changes in the FP composition or organization. The availability of a high-quality dataset describing closely related yet diverse FPs offers intriguing prospects for the evaluation and development of such tools.

In this contribution we aimed to broaden our knowledge of FP structure/function relationships by generating a large structural and spectroscopic dataset of photochromic fluorescent proteins based on just a single protein scaffold, rsGreen0.7 [14]. Starting from this ancestor, we selected 27 mutants that displayed a large variety in spectroscopic properties yet differed in only one or two amino acid mutations, aiming to realize a large spectroscopic diversity without focusing on applicability for a particular use case. We characterized these mutants on 19 different spectroscopic parameters, and also obtained 43 crystal structures of the fluorescent and non-fluorescent states (including the previously published structures of rsGreen0.7 [14]; eleven structures could not be determined with sufficiently high resolution). We identified correlations between the different spectroscopic parameters and/or structural properties, and established a structural picture of the changes underlying the variability in the spectroscopic properties. Overall, our results confirm that the spectroscopic properties of fluorescent proteins are highly correlated, and that it is difficult to *a priori* predict a mutational outcome. Nevertheless, we do identify several fundamental relationships that can ease the FP design process. Our extensive structure-function data not only leads to the conclusions presented here, but also provides a rich dataset that can be leveraged for future investigations into this and other classes of proteins.

## Results and discussion

### Generation of a spectroscopically diverse set of mutants

We based our work on the photochromic green fluorescent rsGreen0.7, a mutant of EGFP that has been used in a range of imaging experiments and methodologies [14]. Using the crystal structures of this mutant and our experience with rsGreen-mutagenesis, we identified a number of amino acids in the chromophore pocket (Phe145, His148, Phe165, Asn205 and Glu222), likely to give rise to a considerable variation in spectroscopic properties. We also introduced the K206A mutation, which enhances the tendency for protein dimerization [20], to increase the likelihood of crystallogenesis. Mutants were generated by subjecting these sites to saturation mutagenesis, followed by spectroscopic characterization using a two-level screening assay consisting of transformation in *E. coli*, in-colony characterization [14], and on-microscope screening in a 96-well format [21]. During the first screening round in bacterial colonies, mutants deemed worthy of further investigation were identified based on their relative brightness, photoswitching kinetics and contrast between the fluorescent and dim state. The second screening step was based on the same optical properties, but the higher sensitivity and illumination intensities of the microscope allowed a more detailed assessment. Mutants that clearly differed in spectroscopic properties from rsGreen0.7 and each other were retained for purification and *in vitro* assessment, without regards to usefulness for a particular purpose apart from the presence of a measurable fluorescence.

All purified fluorescent proteins underwent an extensive spectroscopic characterization in which 19 parameters were evaluated (Table 1 and Figure S1). Figure 1 highlights a few of these measurements, including absorption and excitation/emission spectra at pH 7.4 (Figure 1A,B), the evolution of absorption and emission during photoswitching (Figure 1C) and an expansion of the second photoswitching cycle (Figure 1D). The behavior of Dronpa [22], one of the first photoswitchable proteins identified, is also shown for comparison.

**Figure 1:**
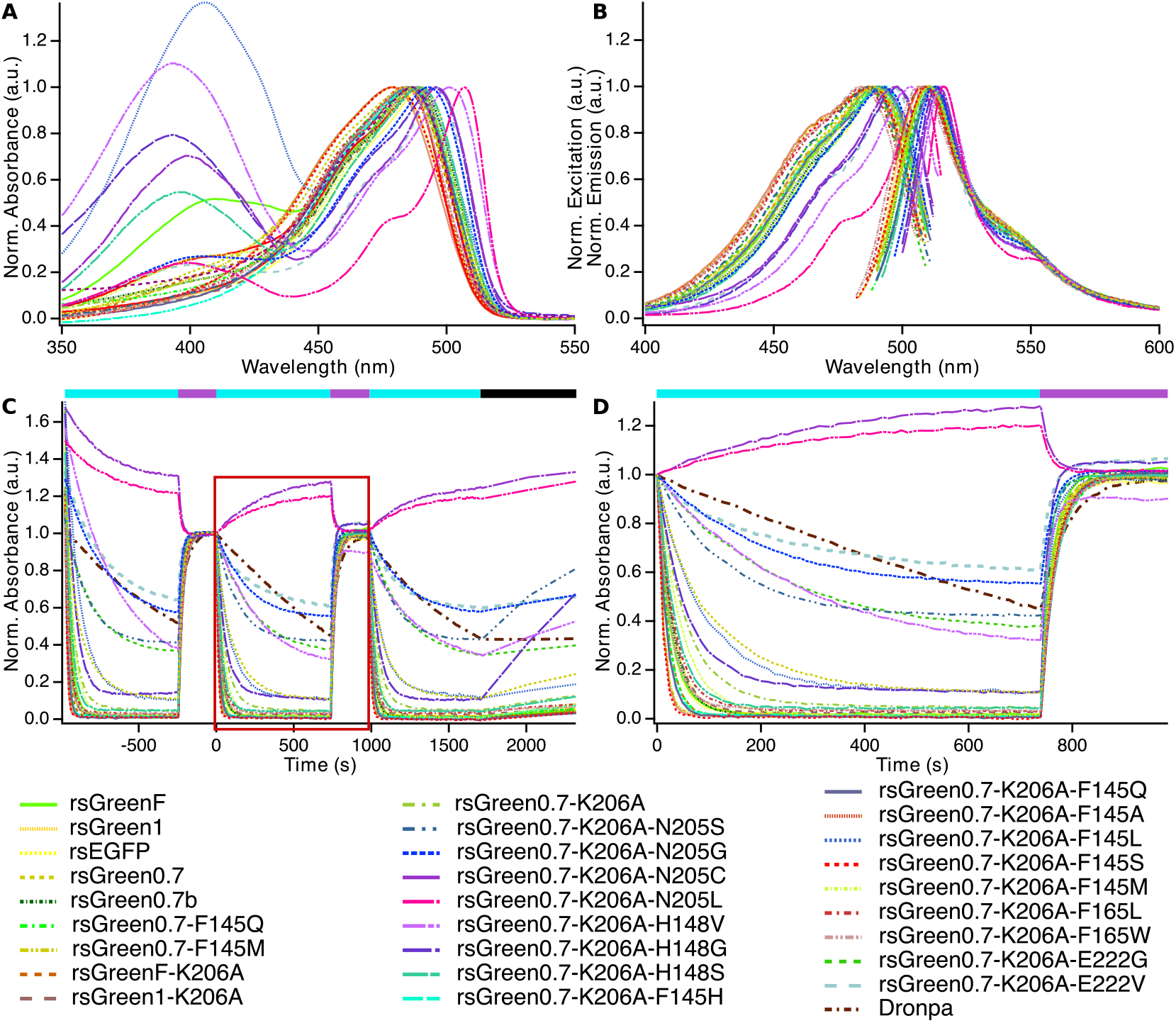
**(A-B)** Normalized absorption **(A)**, excitation and emission **(B)** spectra of rsEGPF and the rsGreen-variants at pH 7.4. **(C)** Normalized time evolution of the anionic absorption band during illumination with 488-nm light (cyan bar), 405-nm light (violet bar) and a thermal recovery period (black bar). Dronpa was added as a reference and is depicted in bold. **(D)** Zoomed-in region from the second cycle that was used to calculate the dynamic macroscopic properties except for A_unrecov_ and k_therm_.

**Table 1:** Spectroscopic properties of rsEGFP, rsGreen1, rsGreenF, rsGreen0.7 and rsGreen0.7-mutants.

| Protein | $\lambda_{\text{abs}}$<br>(nm) | $\lambda_{\text{ex}}$<br>(nm) | $\lambda_{\text{em}}$<br>(nm) | $\lambda_{\text{off}}^{\text{abs}}$<br>(nm) | $\delta_{\text{Stokes}}$<br>(nm) | $\lambda_{\text{ex}} - \lambda_{\text{abs}}$<br>(nm) | $\epsilon_{\text{overall}}$<br>(mM<sup>-1</sup> cm<sup>-1</sup>) | $\epsilon_{\text{anionic}}$<br>(mM<sup>-1</sup> cm<sup>-1</sup>) | $\epsilon_{\text{off}}$<br>(mM<sup>-1</sup> cm<sup>-1</sup>) |
| --- | --- | --- | --- | --- | --- | --- | --- | --- | --- |
| rsGreenF | 486 | 489 [485] | 511 [511] | 391 [390] | 22 | 3 | 35 [41] | 65 [52] | 30 [18] |
| rsGreen1 | 486 | 488 [486] | 510 [509] | 397 [396] | 22 | 2 | 62 [58] | 71 [61] | 26 [20] |
| rsEGFP | 489 | 491 [491] | 510 [510] | 397 [396] | 19 | 2 | 58 [46,47] | 72 [57] | 28 [18] |
| rsGreen0.7 | 484 | 489 [487] | 510 [511] | 397 [396] | 21 | 5 | 46 [48] | 65 [62] | 27 [21] |
| rsGreen0.7b | 489 | 491 [489] | 511 [511] | 396 [396] | 20 | 2 | 55 [44] | 70 [53] | 26 [20] |
| rsGreen0.7-F145Q | 484 | 491 | 512 | 393 | 21 | 7 | 53 | 61 | 21 |
| rsGreen0.7-F145M | 479 | 488 | 510 | 396 | 22 | 9 | 50 | 59 | 25 |
| rsGreenF-K206A | 485 | 490 | 509 | 390 | 19 | 5 | 52 | 70 | 27 |
| rsGreen1-K206A | 487 | 487 | 508 | 396 | 21 | 0 | 75 | 72 | 33 |
| rsGreen0.7-K206A | 488 | 489 | 510 | 399 | 21 | 1 | 56 | 67 | 29 |
| rsGreen0.7-K206A-N205S | 486 | 487 | 513 | 414 | 26 | 1 | 50 | 59 | 19 |
| rsGreen0.7-K206A-N205G | 493 | 493 | 513 | 410 | 20 | 0 | 52 | 74 | 33 |
| rsGreen0.7-K206A-N205C | 496 | 498 | 514 | 496 | 16 | 2 | 21 | 50 | NA |
| rsGreen0.7-K206A-N205L | 507 | 508 | 516 | 507 | 8 | 1 | 64 | 104 | NA |
| rsGreen0.7-K206A-H148V | 501 | 503 | 516 | 394 | 13 | 2 | 25 | 67 | 38 |
| rsGreen0.7-K206A-H148G | 496 | 498 | 514 | 400 | 16 | 2 | 30 | 64 | 40 |
| rsGreen0.7-K206A-H148S | 490 | 491 | 511 | 399 | 20 | 1 | 29 | 62 | 29 |
| rsGreen0.7-K206A-F145H | 485 | 494 | 511 | 395 | 17 | 9 | 59 | 66 | 24 |
| rsGreen0.7-K206A-F145Q | 485 | 492 | 513 | 395 | 21 | 7 | 55 | 61 | 23 |
| rsGreen0.7-K206A-F145A | 478 | 487 | 509 | 397 | 22 | 9 | 53 | 58 | 23 |
| rsGreen0.7-K206A-F145L | 487 | 491 | 513 | 393 | 22 | 4 | 19 | 60 | 35 |
| rsGreen0.7-K206A-F145S | 479 | 488 | 509 | 394 | 21 | 9 | 53 | 58 | 22 |
| rsGreen0.7-K206A-F145M | 488 | 492 | 510 | 393 | 18 | 4 | 46 | 59 | 26 |
| rsGreen0.7-K206A-F165L | 483 | 486 | 509 | 393 | 23 | 3 | 55 | 64 | 23 |
| rsGreen0.7-K206A-F165W | 484 | 484 | 506 | 398 | 22 | 0 | 54 | 64 | 27 |
| rsGreen0.7-K206A-E222G | 487 | 487 | 511 | 412 | 24 | 0 | 68 | 71 | 29 |
| rsGreen0.7-K206A-E222V | 496 | 497 | 511 | 403 | 14 | 1 | 56 | 87 | 34 |

| Protein | pK<sub>a,app</sub> | $\lambda_{\text{abs,pH10}} - \lambda_{\text{abs,pH6.5}}$<br>(nm) | Q<sub>fluo</sub><br>(%) | Molecular<br>brightness | Q<sub>off</sub><br>(10<sup>-1</sup> %) | Q<sub>on</sub><br>(%) | A<sub>res</sub><br>(%) | A<sub>res,∞</sub><br>(%) | k<sub>therm</sub><br>(10<sup>-4</sup> s<sup>-1</sup>) | A<sub>unrecov</sub><br>(%) |
| --- | --- | --- | --- | --- | --- | --- | --- | --- | --- | --- |
| rsGreenF | 7.2 [6.7] | 2 | 39 [39] | 14 | 7.3 [8.7] | 12.7 [1.8] | 1.7 | 1.6 | 0.5 | 40.0 |
| rsGreen1 | 6.2 [5.9] | 0 | 43 [42] | 27 | 3.7 [4.2] | 14.0 [1.4] | 1.2 | 1.0 | 0.4 | 8.6 |
| rsEGFP | 6.6 [6.7,6.5] | 2 | 42 [42, 36] | 24 | 4.5 [5.5] | 14.6 [1.7] | 2.0 | 1.9 | 0.6 | 15.8 |
| rsGreen0.7 | 6.9 [6.7] | 16 | 30 [40] | 14 | 1.3 [2.6] | 14.6 [1.2] | 10.7 | 8.2 | 2.6 | 16.5 |
| rsGreen0.7b | 6.8 [6.7] | 6 | 38 [43] | 21 | 4.1 [5.3] | 13.9 [1.6] | 1.3 | 1.2 | 0.4 | 18.7 |
| rsGreen0.7-F145Q | 6.3 | 0 | 32 | 17 | 6.7 | 13.8 | 0.9 | 0.7 | 0.8 | 8.0 |
| rsGreen0.7-F145M | 6.5 | 2 | 23 | 12 | 12.8 | 14.0 | 1.5 | 1.4 | 0.8 | 16.8 |
| rsGreenF-K206A | 6.8 | 0 | 43 | 22 | 6.9 | 14.5 | 1.0 | 1.0 | 0.4 | 25.1 |
| rsGreen1-K206A | 6.3 | 0 | 47 | 35 | 3.0 | 11.6 | 3.0 | 2.9 | 0.5 | 9.5 |
| rsGreen0.7-K206A | 6.9 | 2 | 39 | 22 | 1.9 | 12.2 | 4.8 | 4.0 | 1.4 | 18.4 |
| rsGreen0.7-K206A-N205S | 6.6 | 0 | 49 | 25 | 1.7 | 17.9 | 42.4 | 25.5 | 17.8 | 7.7 |
| rsGreen0.7-K206A-N205G | 7.1 | 2 | 43 | 22 | 0.4 | 10.4 | 55.3 | 51.9 | 3.7 | 22.7 |
| rsGreen0.7-K206A-N205C | 7.7 | 0 | 22 | 5 | NA | NA | NA | NA | NA | NA |
| rsGreen0.7-K206A-N205L | 7.4 | 0 | 30 | 19 | NA | NA | NA | NA | NA | NA |
| rsGreen0.7-K206A-H148V | 7.7 | 2 | 33 | 8 | 1.1 | 13.4 | 32.3 | 18.1 | 6.7 | 36.3 |
| rsGreen0.7-K206A-H148G | 7.8 | 4 | 40 | 12 | 3.9 | 14.2 | 11.2 | 2.6 | 15.3 | 39.2 |
| rsGreen0.7-K206A-H148S | 7.5 | 2 | 51 | 15 | 6.1 | 16.9 | 4.4 | 3.9 | 1.4 | 30.5 |
| rsGreen0.7-K206A-F145H | 6.1 | -2 | 72 | 42 | 6.3 | 13.4 | 1.0 | 0.8 | 1.1 | 8.1 |
| rsGreen0.7-K206A-F145Q | 6.2 | 0 | 39 | 21 | 6.9 | 13.8 | 0.6 | 0.5 | 0.5 | 10.8 |
| rsGreen0.7-K206A-F145A | 6.2 | 0 | 35 | 19 | 13.2 | 13.2 | 1.1 | 1.1 | 0.5 | 13.3 |
| rsGreen0.7-K206A-F145L | 7.8 | 2 | 50 | 9 | 3.6 | 13.4 | 11.1 | 9.9 | 1.5 | 41.5 |
| rsGreen0.7-K206A-F145S | 6.5 | 2 | 27 | 14 | 13.5 | 12.0 | 0.4 | 0.3 | 0.7 | 17.7 |
| rsGreen0.7-K206A-F145M | 7.1 | 0 | 56 | 26 | 3.3 | 14.0 | 1.7 | 1.1 | 1.2 | 31.5 |
| rsGreen0.7-K206A-F165L | 6.6 | 2 | 28 | 16 | 4.0 | 13.3 | 2.9 | 2.6 | 0.8 | 11.9 |
| rsGreen0.7-K206A-F165W | 6.5 | 2 | 40 | 22 | 5.0 | 10.3 | 3.3 | 3.2 | 0.4 | 9.9 |
| rsGreen0.7-K206A-E222G | 6.4 | 0 | 52 | 36 | 0.4 | 15.5 | 37.9 | 35.9 | 1.4 | 5.9 |
| rsGreen0.7-K206A-E222V | 7.5 | 2 | 38 | 21 | 0.3 | 12.5 | 61.2 | 59.2 | 2.4 | 24.5 |

### Exploratory analysis of the spectroscopic data

Correlations analysis can identify those parameters that vary in concert due to similar structural dependencies. We therefore calculated Pearson correlation coefficients over our data (Table 2), which allowed us to identify several correlated spectroscopic parameters that hint to shared underlying mechanisms, while providing information that can be used to accelerate future FP engineering efforts. Whereas strong correlations between *λ*_abs_, *λ*_ex_ and *λ*_em_ or between Q_fluo_ and molecular brightness are expected, we also found several relationships that were less anticipated. For example, the difference between the maximal absorption and excitation wavelengths (*λ*_ex_ – *λ*_abs_) positively correlates with the off-switching quantum yield Q_off_. This wavelength difference is usually seen as an indicator of ground state heterogeneity within an FP-ensemble, and is consistent with the presence of at least two populations with different photoswitching propensity and brightness in the ground state[21, 24]. We will discuss particular insights enabled by these correlations further along in this contribution.

**Table 2:**
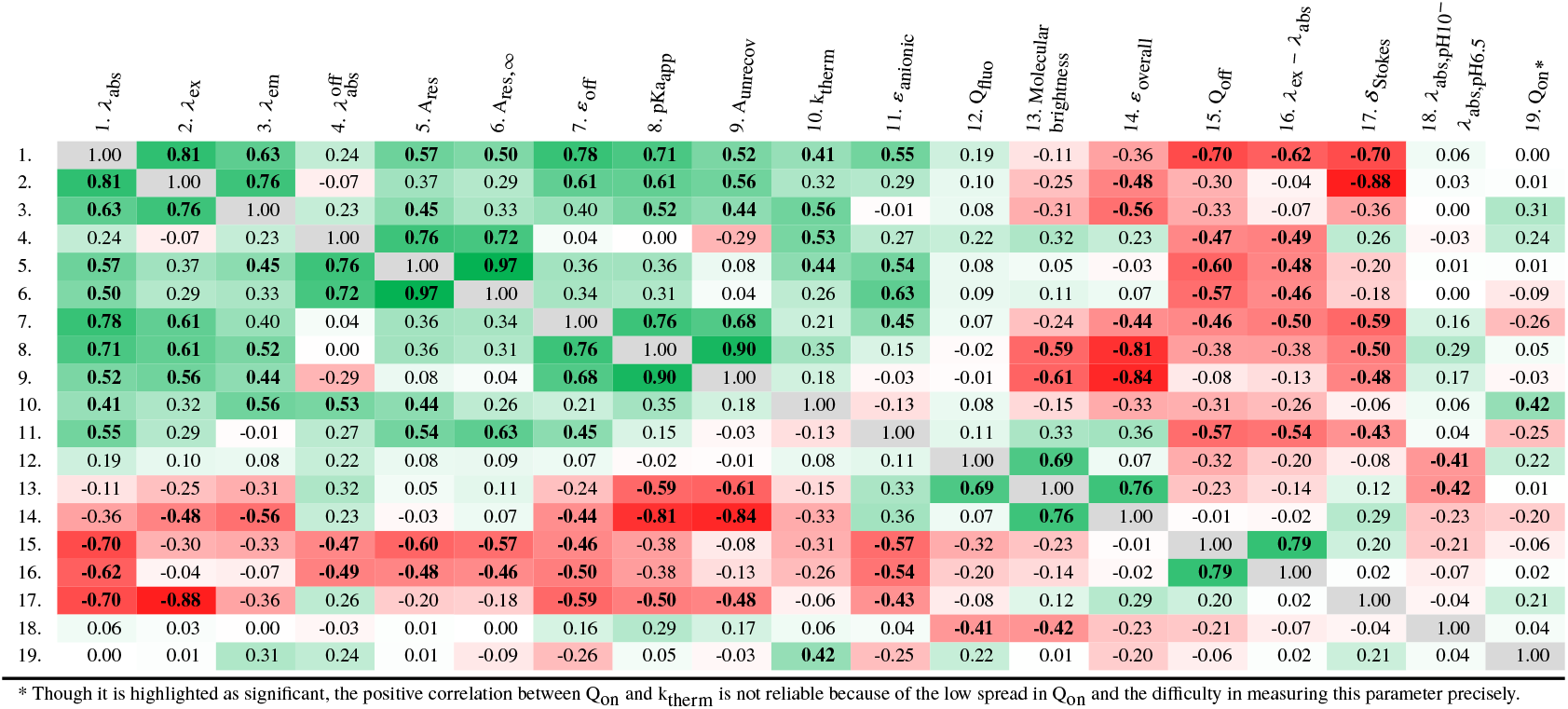
Pearson correlation matrix of the spectroscopic properties. Color code from green (r=1) over white (r = 0) to red (r = −1). Values in bold display statistically significant (*α* = 0.05) correlations. The positive switchers (rsGreen0.7-K206A-N205C and rsGreen0.7-K206A-N205L) were left out of the calculation.

One limitation of this correlation analysis is that it cannot reveal intricate dependencies between multiple parameters, nor whether our mutants can be separated into different groups depending on their properties and structures. Given this substantial dataset, we were interested in exploring this question using principal component analysis (PCA). PCA identifies linear combinations of properties (principal components, PCs) such that the variance of the projection of the protein properties along these PCs is maximized. The obtained principal components are ranked in terms of the variance of the data that they capture, thereby allowing one to make conclusions about linear relationships between the properties. For example, if projecting the protein properties onto the first principal component captures all of the variance, then the sample variability is presumably due to a single mechanism in which the parameters change in the same coordinated manner. We refer to the tutorial review of Bro and Smilde [25] for more information on this analysis methodology.

We applied PCA to the spectroscopic dataset, excluding the two positive switchers (rsGreen0.7-K206A-N205C and rsGreen0.7-K206A-N205L) because their properties cannot be consistently grouped (Q_off_, for example, must be defined differently for positive switchers). We found that 10 principal components were required to capture 98.2% of the observed variance, suggesting that the FP variability does not arise through just a single or a few different mechanisms. The results were subjected to a visual analysis detailed in Supporting Discussion 1.1. To connect the spectroscopic parameters to the structural modifications, we augmented this analysis by labeling each mutant according to where the corresponding mutations are situated with respect to the chromophore. We divided the chromophore environment into three different zones, consisting of zone 1 (amino acid 205 and 222), zone 2 (amino acid 145) and zone 3 (amino acid 148 and 165) (Figure 2, Table S1).

**Figure 2:**
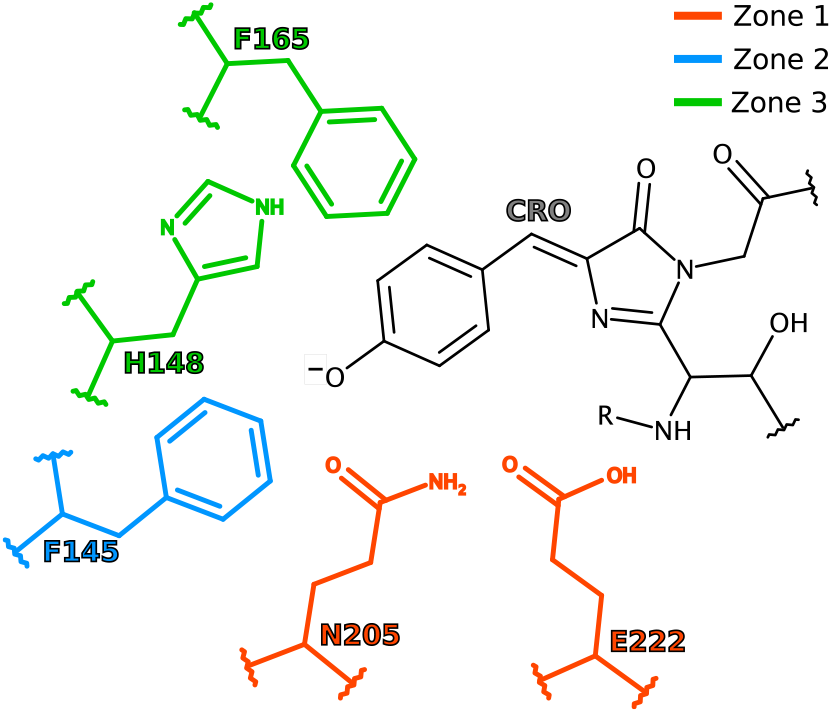
Classification of the FPs according to the position of their mutated amino acid in relation to rsGreen0.7 or rsGreen0.7-K206A.

Ultimately, our methodology allowed us to construct Table 3, which summarizes the changes in spectral parameters as a function of the chromophore zone. This table shows how mutants modified in each chromophore zone differ from a theoretical FP that has properties equal to the averages of the properties of our mutants, and suggests the parameters that will be modified by introducing a mutation at that particular position. For example, introducing a mutation in zone 1 is likely to result in a change in *λ*_ex_ – *λ*_abs_ and Q_off_. This table also presents a modest guide for further engineering: for example, a faster switcher can be created by adding a mutation in zone 2, though this is likely to also introduce additional spectroscopic heterogeneity as expressed by the *λ*_ex_ – *λ*_abs_ parameter. However, it includes only a subset of all spectroscopic properties, and no properties at all for zone 3 mutants, reflecting the difficulty in extracting clear conclusions from the overall complexity associated with FPs.

**Table 3:** Principal component analysis on spectroscopic properties indicating the effect of mutation compared to the average.

| Zone1<br>E222X and N205X | Zone 2<br>F145X | Zone3<br>H148X and F165X |
| --- | --- | --- |
| $\lambda_{\text{ex}} - \lambda_{\text{abs}} \downarrow$ | $\lambda_{\text{ex}} - \lambda_{\text{abs}} \uparrow *$ | No consistent changes identified |
| $\lambda_{\text{abs,off}} \uparrow$ | $\lambda_{\text{abs}} \downarrow *$ | |
| $A_{\text{res}} \uparrow$ | $\epsilon_{\text{overall}} \sim \uparrow$ | |
| $A_{\text{res},\infty} \uparrow$ | $\text{pka}^{\text{app}} \downarrow *$ | |
| $Q_{\text{off}} \downarrow$ | $k_{\text{therm}} \downarrow$ | |
| | $\lambda_{\text{abs,off}} \downarrow$ | |
| | $Q_{\text{off}} \uparrow *$ | |

↑ : higher than average, ↓ : lower than average, ∼ : equal to average, $*$ : exceptions present.

### Inclusion of structural information

We then sought to augment our dataset by including direct structural information. We solved the structures of our mutants in both the on- and off-state using X-ray crystallography, retaining only those samples that diffracted to a resolution of 2.5 Å or better (43 structures in total, including also the structures of rsGreen0.7 published previously [14]; Table S2, Supplementary discussion 1.2). In several structures, a mixture of both *cis* and *trans* states are present, likely due to incomplete photoswitching or partial off-switching during crystal handling. However, reliable structural models for the on- and/or off-states could still be extracted if these displayed high enough occupancies, or when the crystal structure of the protein in the other state was available.

All the characterized proteins manifested as 11-stranded β-barrels with a central helix bearing the chromophore (Figure S2). The conformations of the chromophores are also highly similar, except for the three mutants at position His148, where the hydroxybenzylidene ring is almost coplanar with the imidazolinone in the off-state (Figure S3). Where it is present, His148 moves to take up the space created during the *cis*-to-*trans* isomerization of the chromophore and causes the chromophore to adopt a twisted conformation, indicating that it has an impact on the photo-isomerization efficiency.

Recently, the crystal packing and solvent content were shown to influence the *in crystallo* photoswitching in an rsEGFP2 variant possessing a chlorinated chromophore, giving rise to two possible off-state structures. [26] While the FPs in our study crystallize in different space groups, the off-state chromophore conformations do superpose on each other and resemble those observed in the “expanded crystal” state defined in Ref. [26], except for the structures of the H148 mutants (as noted above). This suggests that the crystals did not dehydrate during the experimental manipulations, nor that off-switching was constrained to follow different pathways through crystallization.

Using this structural data, we expanded our PCA-based analysis methodology by adding a total of 95 different structural descriptors, detailed in the Materials and Methods section and Table S3. These properties can be divided into five groups, relating to (i) properties of the chromophore itself, (ii) distances between the chromophore and nearby atoms, (iii) distances between different amino acids surrounding the chromophore, (iv) angles between the planes of nearby aromatic amino acids and the chromophore hydroxybenzylidene ring, and (v) general descriptors of the chromophore environment.

We redid the correlation analysis and PCA using the spectroscopic and structural descriptors associated with the on-state crystal structures. We find that 13 PCs are required to obtain 90.67% explained variance (Figure S4 and S5, Table S4, S5 and S6). We were similarly able to identify specific changes associated with mutations at a particular zone of the chromophore, analogous to the information shown in Table 3. This is shown in Table S7 rather than in the main text due to the large size of this table.

Performing the same analysis on the off-states was complicated by the lower number of off-state structures available. Furthermore, we were forced to exclude the H148 mutants from consideration due to their different off-state chromophore conformation. The remaining off-state structures mainly belong to zone 2 (Table S1), rendering the previously used zone classification unusable for the off-state analysis. Instead we classified the FPs according to their off-switching quantum yield (Q_off_) resulting in ‘fast’, ‘medium’ and ‘slow’ switchers (Table S1; Figure S6, Table S8 and S9). This lead to the insights summarized in Table S10.

### Combined spectroscopic-structural analysis provides insights into FP functioning

Taken together, our dataset and analysis provide several insights into FP functioning.

#### Hydrogen bonding to water molecules is a key determinant of the photochromism efficiency

The correlation and principal component analysis reveal that fast switchers show more hydrogen bonds with the chromophore (Table S7), most of which involve water molecules (Figure 3A and B). These water molecules are present when mutations of residues next to the chromophore hydroxybenzylidene moiety result in side chains that are smaller in size and more polar. In particular, the presence of F145 leads to slow off-switching that is accelerated by its substitution with smaller amino acids. The opening in the *β*-barrel next to F145 and the hydroxybenzylidene oxygen allows water molecules to be in relatively fast exchange with surrounding water, as compared to other water molecules in the chromophore pocket [27, 28]. Water mobility has been linked to protein interaction and enzymatic kinetics for a wide range of proteins, where internal and external water molecules play an active role in the protein dynamics. [29–31] The dynamic nature of the water molecules thus presumably accelerates the *cis/trans* isomerization by being sufficiently mobile to stabilize a broad range of different conformations involving the chromophore and/or the surrounding amino acids. Indeed, the increased dynamics around the hydroxybenzylidene oxygen are confirmed by the higher crystallographic B-factors for the hydrogen bond interaction partners in the fast switchers as compared to the slow switchers. Intriguingly, a previous computational study [24] reported that photo-isomerization is precluded in the presence of multiple hydrogen bonds with the chromophore hydroxybenzilidene oxygen, since such bonding interferes with the charge transfer required for photo-isomerization. Our observations suggest that this restriction may be less absolute, at least for this FP scaffold, though it does align with the notion that the fast H-bonding rearrangements enabled by the mobile water molecules favor photochromism by facilitating the transient breaking of the hydrogen bonds to the chromophore.

**Figure 3:**
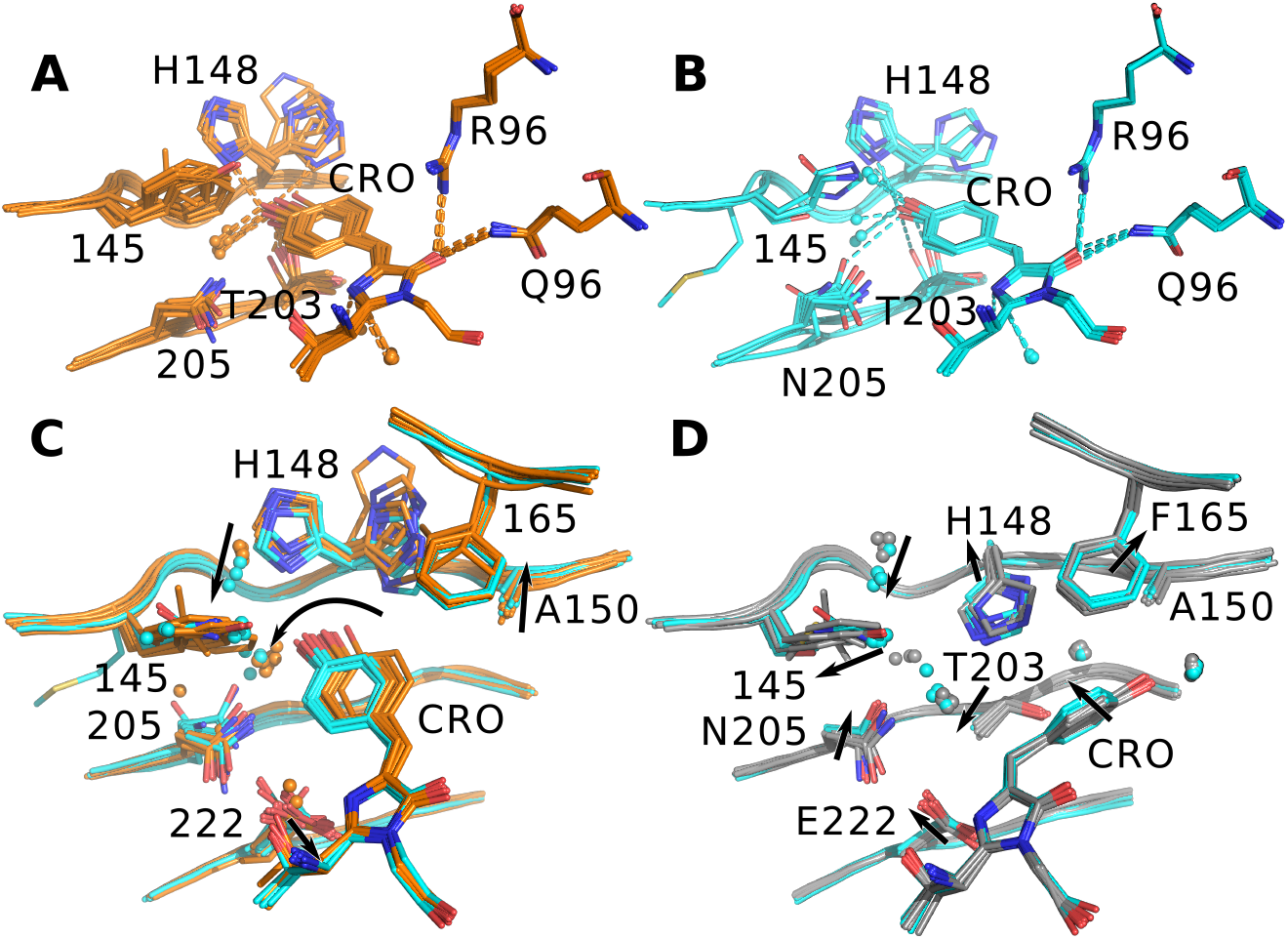
**(A and B)** Hydrogen bonds made by the on-state chromophore of the FPs belonging to the slow **(A)** and the fast switchers **(B)** (list of mutants provided in Table S1), indicating the correlations between the number of hydrogen bonds made by the hydroxybenzylidene oxygen and Q_off_ (r = 0.58, p < 0.01) and λ_ex_ – λ_abs_ (r = 0.72, p < 0.001). Hydrogen bonds are depicted by dashes (criterium: length ≤ 3.2 Å). If due to alternative conformations multiple hydrogen bond possibilities exist, then only the shortest interaction is shown. If the chromophore has alternative conformations, then only the hydrogen bonds made by a single conformation are drawn. **(C and D)** Structural superposition of the fast switchers with slow switchers in the on-state (**C**; cyan and orange, respectively) and off-state (**D**; cyan and grey, respectively). Arrows point to the main structural shifts.

This observation also offers an explanation for the high *λ*_ex_ – *λ*_abs_ values observed for fast switchers, since the dynamic water molecules may accommodate a variety of ground state conformations with varying fluorescence brightnesses. The fast switchers also show a larger tilt angle and rotation of the chromophore around the imidazolinone nitrogen atom. This probably also is a consequence of the established hydrogen bonds (Figure 3 C), since these are largely oriented in a direction that favors this chromophore geometry. [32]

We also observe that the F145M mutant is a fast switcher, which is surprising because methionine is large and rather apolar. However, a detailed inspection of the crystal structure reveals that Met145 is present in two conformations, one of which points away from the chromophore and makes room for the inclusion of water molecules (Figure 3B,C and Supplementary Discussion 1.3), and which is presumably responsible for the faster switching. Intriguingly, the F145M/K206A double mutant displays considerably slower switching compared to the F145M mutant. Residue 145 is located close to residue 206, suggesting that the increased protein dimerization induced by the K206A mutation interferes with the outward conformation of Met145.

However, the simple presence of water molecules near the chromophore is not sufficient to confer fast switching: the FPs mutated at the zone 1 region switch more slowly than the other FPs. The N205G and E222G/V mutations both lead to the introduction of water molecules next to the chromophore imidazolinone, although these changes lead to even slower switching. This shows that the positioning of the water molecules is important, and that it is the water molecules situated around the hydroxybenzilidene oxygen of the chromophore that are important for enhancing the switching quantum yield.

Steric hindrance has also been considered to be a key determinant for photochromism [12, 26, 33]. We attempted to capture this effect by quantifying the chromophore pocket volume. We find that the on-state volume does not correlate with the off-switching quantum yield, indicating that steric hindrance may be less important, or that it may not be captured well by this descriptor. However, this conclusion needs to be mitigated somewhat when considering that the pocket volume descriptor is sensitive to the overall chromophore environment, including also the changes at positions 205 and 222, which should not cause steric hinderance of the hydroxybenzylidene during isomerisation. This suggests either that steric hindrance may be less important, or that it may not be captured well by the total on-state volume. On the other hand, we do find a strong correlation between the off-switching quantum yield and the chromophore volume in the off-state. In the off-state, the positions of the mutated residues show much less variability (Figure 3 D), and therefore the changes in the chromophore pocket volume are dominated by the variability at positions 145, 148, and 150, indicating that steric hindrance by these residues may play a role. While it was difficult to observe a clear effect of steric hindrance through analysis of the pocket volume, the steric hindrance at these residues does appear to be implicated in the off-switching quantum yield.

Overall, the off-switching efficiency appears to be largely determined by the hydrogen bonding to the hydroxybenzylidene oxygen, in line with previous work that highlighted the importance of electrostatic effects [34], and whether these hydrogen bonds arise from dynamic bonding partners. In this light, we suggest that the reduced steric hindrance previously suggested as cause for altered photochromism can also finds its origin in the increased probability for favorable water interactions and stabilization of multiple chromophore conformations.

#### The interplay of multiple structural parameters determines the absorption maximum

Our spectroscopic data shows pronounced differences in the peak absorption wavelength *λ*_abs_. The H148 mutants (forming zone 3 together with the F165 mutants) and some zone 1 mutants have a red-shifted absorption peak compared to the other FPs (Table 1, Figure S1). The PCA cannot clearly isolate these red-shifted FPs from other FPs (Figure S7), while the correlation analysis identifies only a few significant, but weak, correlations between *λ*_abs_ and the structural descriptors (Table S6). This indicates that the absorption wavelength is determined by multiple structural aspects that are difficult to untangle. However, our analysis does allow us to identify a number of prominent contributors.

The correlation analysis reveals that the red shift correlates with a reduction in the number of hydrogen bonds that are made to the hydroxybenzylidene oxygen atom (Figure S8A). This also explains why fast switchers display a blue-shift in their absorption maximum, as indicated by the strong anticorrelation between *λ*_abs_ and Q_off_ (Table 2). Several studies, many involving QM/MM, have revealed that (de-)stabilization of the chromophore charge distribution, both in the ground and excited state, determines the absorption maximum.[35–39] The excited state of the FP chromophore displays charge transfer away from the hydroxybenzylidene moiety [34, 40–42]. Increased hydrogen bonding to the hydroxybenzylidene oxygen stabilizes the ground state and destabilizes the excited state, resulting in a blue-shift.

The blue-shifted absorption of rsGreen0.7-K206A-F165W seems to be contradictory because of the low number of hydrogen bonds. The W165 indole can also engage in electron-donating π-π interactions with the hydroxybenzylidene ring (Figure S8B), which are assumed to stabilize the excited state charge transfer and lead to a red-shift. [34, 39, 43–45] Instead, the observed blue-shift likely arises because the stacking perturbs the conjugation of the chromophore (tilt, twist and methylene bridge bond angles of −156°, −21° and 129° as compared to the average of −164°, −9° and 127°, respectively; see Figure S9 for angle definitions). This indicates that geometrical constraints on the chromophore can cancel the effects of hydrogen bonding and other chromophore interactions. [37, 44]

However, the variation within the F145 mutants (zone 2 mutants) is only marginally captured by these two parameters. Whereas direct interactions, such as hydrogen bonding, with the chromophore can be expected to have the highest influence on the absorption maximum, indirect interactions can also play a role, such as a blue-shifting through the presence of polar residues [39, 45]. However, the net effect of these polar residues appears to be minor: proteins with Y145 (rsEGFP and rsGreen07.b) are actually red-shifted compared to their non-polar residue-containing counterparts (rsGreen0.7 and rsGreenF) while the introduction of H/Q145 does not introduce the blue-shift that might be expected from these polar residues. Other effects may also be too small to be captured in the X-ray structures, such as a possible dependence of *λ*_abs_ on bond-length variations. [35, 46].

Taken together, our results show that *λ*_abs_ depends on multiple structural aspects and is difficult to explain using simple considerations, though hydrogen bonding and chromophore planarity play a key role.

#### Hydrogen bonding networks and steric hindrance appear as main contributors to thermal recovery

Three FPs, two of them belonging to the H148 mutants, show rates of thermal recovery (k_therm_) that are much higher than those observed on the other mutants (Table 1, Figure S1). The H148 mutants have an altered chromophore off-state conformation, making it difficult to directly compare the recovery process with the other mutants. However, k_therm_ shows considerable variation among the three H148 mutants, suggesting that the comparison of these three FPs can hint to key structural contributions to k_therm_.

The fastest recovery (15.3×10^−4^ s^−1^ M^−1^) is observed for the H148G mutant. This mutation leads to a chromophore pocket that is directly open to the surrounding bulk solvent, characterized by fewer structured water molecules and heteroatoms close to the hydroxybenzylidene. Unfortunately, no crystal structure of its off-state could be determined, though the unconstrained on-state chromophore pocket suggests a very low steric hindrance [47] and the absence of large conformational changes during spontaneous chromophore *trans-to-cis* isomerization. In contrast, the H148S mutant shows a slow thermal recovery (1.4×10^−4^ s^−1^ M^−1^). We find that S148 forms part of an extended hydrogen bond network both in the on- and off-state (Figure 4), which has to be broken and rebuilt during *trans-to-cis* isomerization, requiring multiple conformational changes and thus bringing down the thermal recovery rate. Finally, the H148V mutant shows an intermediate thermal recovery (6.7×10^−4^ s^−1^ M^−1^). Valine is slightly larger than serine but apolar and does not support hydrogen bonding, suggesting that its main effect is the result of steric hindrance. We therefore conclude that thermal recovery is affected by both steric hindrance and the hydrogen-bonding interaction network around the chromophore.

**Figure 4:**
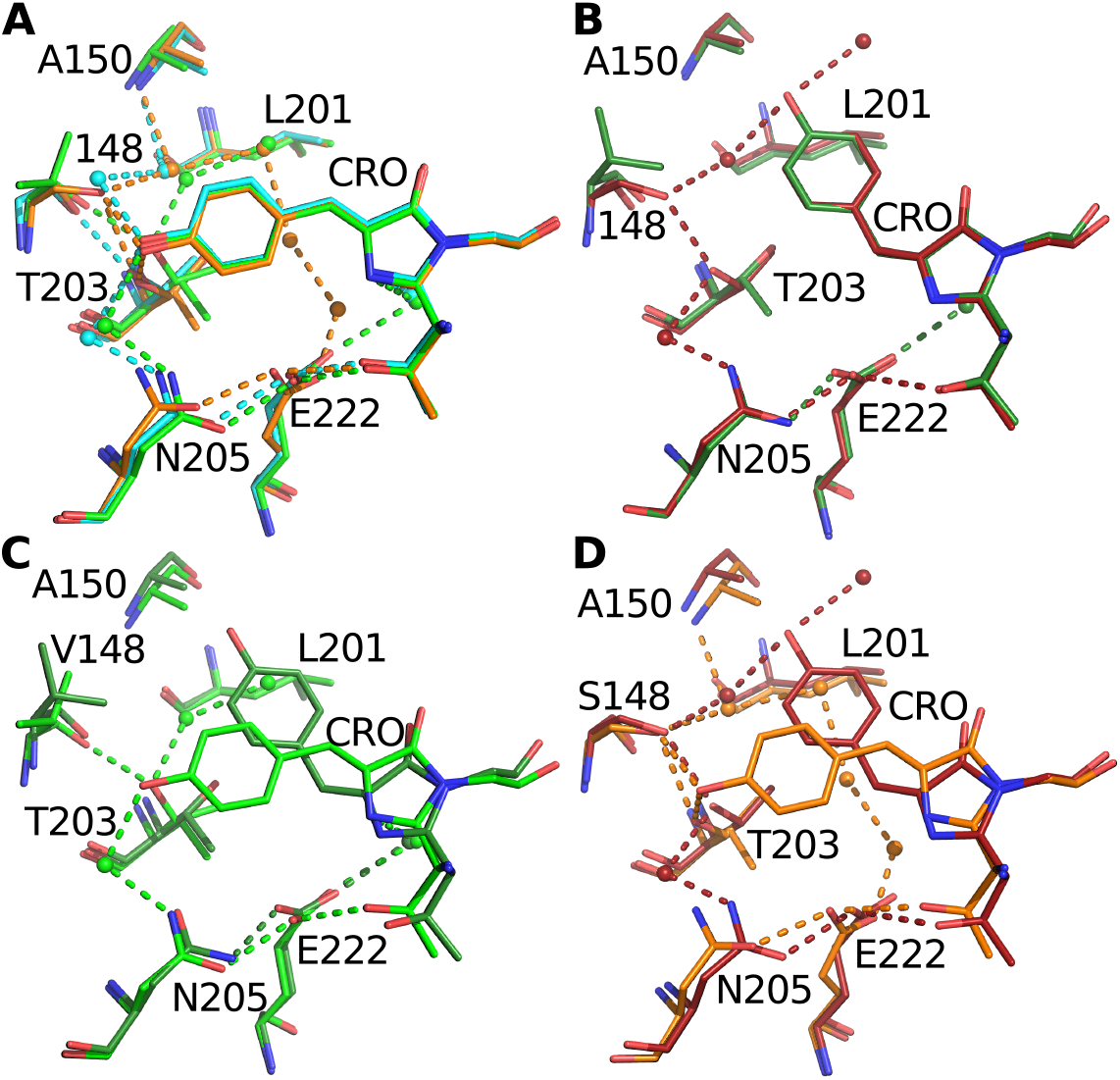
Superposition of the on-state **(A)** and off-state **(B)** chromophore and environment of rsGreen0.7-K206A-H148S (orange, dark red), rsGreen0.7-K206A-H148V (green, dark green) and rsGreen0.7-K206A-H148G (cyan), showing the interaction network around the chromophore. Interactions (hydrogen bonds or electrostatic) are depicted by dashes (criterium: length ≤ 3.2 Å). **(C and D)** Comparison of the on-and off-state interactions made by rsGreen0.7-K206A-H148V and rsGreen0.7-K206A-H148S.

#### Correlation analysis reveals different responses to the polarity of the chromophore environment

An intriguing finding from our data is the presence of the large green and red ‘blocks’ in the correlation table (Table 2, green: spectroscopic parameter 1-11; red: spectroscopic parameter 13-17). This suggests that the spectroscopic parameters divide into two groups that show opposite responses to mutations, though some of the correlations are comparatively weak. We propose that these two groups arise on account of their opposite sensitivity to the introduction of (a-)polar groups in the chromophore environment. Such polar groups can exert influence by engaging in direct hydrogen bonding and/or by creating a generally more polar environment. The correlation of a particular spectroscopic parameter with the number of hydrogen bonds to the chromophore can offer insight into the origin of a particular change. For example, blue shifting of the absorption maximum appears to be mainly determined by the establishment of hydrogen bonds, while the pKa seems to be rather influenced by a global polar environment (correlation between λ_abs_, pKa and number of H-bonds made by hydroxybenzylidene oxygen −0.44 and −0.29, being significant and non-significant (α = 0.05), respectively).

The spectroscopic responses are further modulated by the positioning of these polar/apolar groups relative to the chromophore. This explains why there may be no significant correlation between individual parameters, even though they are part of the same ‘block’ in the correlation table. Indeed, the Stokes shift appears to be less susceptible to mutating residues 145 and 165, but is highly affected by mutations on position 205 and 222 which are closer to the imidazolinone ring compared to the other mutations done in this study. The impact of these mutation sites is reversed for Q_off_, which is why no correlation is observed between the Stokes shift and Q_off_ even though both take part in the red block in the correlation table.

Our structural analysis supervises only the chromophore pocket, while mutations can have effects on much longer distances and the whole β-barrel can influence the spectroscopic behavior of FPs. For example, the altered properties of rsGreen1 compared to rsGreen0.7 must arise from structural differences further away from the chromophore, since they have the same amino acid composition in the chromophore pocket, which adds an extra layer of complexity to structure/spectroscopic function relationships. This was also observed during the PCA in which rsEGFP was classified as a zone 2 FP based on its mutation in the chromophore pocket, but mostly behaved as an outlier. Finally, the spectroscopic properties can be affected by dynamic or electronic effects that are not observable in the crystal structure.

## Conclusions

In conclusion, we have generated a collection of 27 photochromic fluorescent proteins that show a broad spectroscopic diversity, yet differ in only one or two mutations. We characterized their spectroscopic properties using 19 different parameters and also obtained 43 crystal structures. Our work distinguishes itself from previous efforts in that we aimed to recruit a broad diversity of labels, without emphasizing fitness for a particular purpose, while restricting the number of mutations present.

Our study reinforces the notion that comparatively small biochemical changes can trigger large changes in the spectroscopic properties of a fluorescent protein. Nevertheless, our correlation analysis was able to identify multiple significant correlations across our dataset, which can accelerate FP engineering by identifying parameters that are likely to be linked. We then expanded our analysis by including a methodology based on principal component analysis, allowing us to identify correlated changes associated with mutations at particular residues. However, accounting for the dataset variability requires the inclusion of a large number of principal components, suggesting that this variability cannot be considered to arise from a limited number of underlying mechanisms. These findings emphasize the difficulty of rational FP design. Nevertheless, we were able to infer parameters that are likely to be changed by mutating at a particular position relative to the chromophore.

The inclusion of structural data allowed us to identify a number of broad parameters that are (some of) the main levers by which FP properties are determined. We find that hydrogen bonding to the hydroxybenzilidene oxygen by water molecules is a key determinant of the photochromism efficiency. Thermal recovery, in contrast, is controlled by restrictions to conformational changes due to hydrogen bonding but also contains clear contributions from steric hindrance. Other effects, such as shifts in the absorption wavelength, are less easy to untangle and appear to arise from multiple mechanisms. Furthermore, our correlation analysis suggests that the polarity of the chromophore pocket is a key determinant for many of the spectroscopic properties, though their response depends on whether they are mainly affected by hydrogen bonding or by the overall polarity of the pocket.

In addition to the insights developed here, our work presents a high-quality and consistent dataset of closely related FP mutants including both spectroscopic and structural information, all of which is freely available via this publication and the Protein Data Bank. We anticipate that our dataset can open opportunities for the development and evaluation of new and existing protein engineering methods, such as computational modelling and machine learning.

## Acknowledgements

We are grateful to Gerrit Groenhof (University of Jyväskylä) and Jeremy Harvey (KU Leuven) for critical insights and discussion. E.D.Z. and L.V.M. thank the beamline staff from X06DA at the Swiss Light Source (Villigen, Switzerland), Proxima1 and Proxima2A at synchrotron Soleil (Gif-sur-Yvette, France), XRD1 at Elettra (Trieste, Italy) and I02 at Diamond Light Source (Oxfordshire, UK) for assistance during X-ray diffraction data collection. E.D.Z., S.H., and S.D. thank the Research Foundation Flanders (FWO) for a doctoral fellowship and postdoctoral fellowships. This work was supported through funding from the Research Foundation Flanders through grants 1514319N, G090819N, G0B8817N, and the European Research Council through grant 714688 NanoCellActivity.

## Supporting information

Experimental procedures, individual absorption and emission spectra, pH titrations, light titrations, details on the analysis procedures, crystallographic information.

