## Supplementary material for "Integrated structure-function dataset reveals key mechanisms underlying photochromic fluorescent proteins"

September 25, 2020

### 1 Supporting discussion

#### 1.1 Principal component analysis and interpretation

We find that the first three principal components (PCs) explain roughly 70% of the sample variance (35.21%, 21.45% and 11.87%; Table S4A). Limiting our analysis only to PC1, PC2 and PC3 would be equivalent to assigning an excessive 30% of the sample variance to measurement uncertainty. Our data therefore indicates that more PCs must be included to capture the fluorescent protein variability. Overall, this highlights the complex relationships between the spectroscopic parameters, suggesting that no single trend or mechanism accounts for the observed changes in spectroscopic properties.

The classification of the mutants according to their structural ‘zones’ (main text Figure 2 was shown to be meaningful by partial-least squares discriminant analysis (PLS-DA) for at least the mutants classified as zone 1 and zone 2 (Supplementary discussion 1.4, Figure S10 and Tables S11A and B).

An example analysis is shown in Figure SD1A, showing the first and fifth PC. The upper-left quadrant in the score plot is occupied by mutants in zone 2, suggesting zone 2 mutants share a correlated set of spectroscopic changes along the directions of the PCs. Furthermore, since these zone 2 mutants score negatively on PC 1 and positively on PC 5, the parameters that distinguish these FPs from the others are those with loadings that contribute to both PCs with opposite signs (Figure SD1B). We applied this strategy to the first 10 PCs (covering 98.2% of the observed variance), providing the insights that lead to Table 3 in the main text.

#### 1.2 The resolution of the crystal structures

The crystal structures discussed within this work were determined with a high-to-medium resolution between 0.97 Å and 2.5 Å. Not all crystals appeared to exist exclusively in one of the two states. Indeed, efficient photoswitchers might have a noticeable fraction of the off-state populating the on-state structures, while less-efficient photoswitchers can have a substantial presence of on-state in their off-state structures. Because of this and the difference in resolution, different levels of details can be found in the structures. In the high-resolution structures, the atom displacement parameters were anisotropically refined whereas in the medium-resolution structures isotropic descriptions were

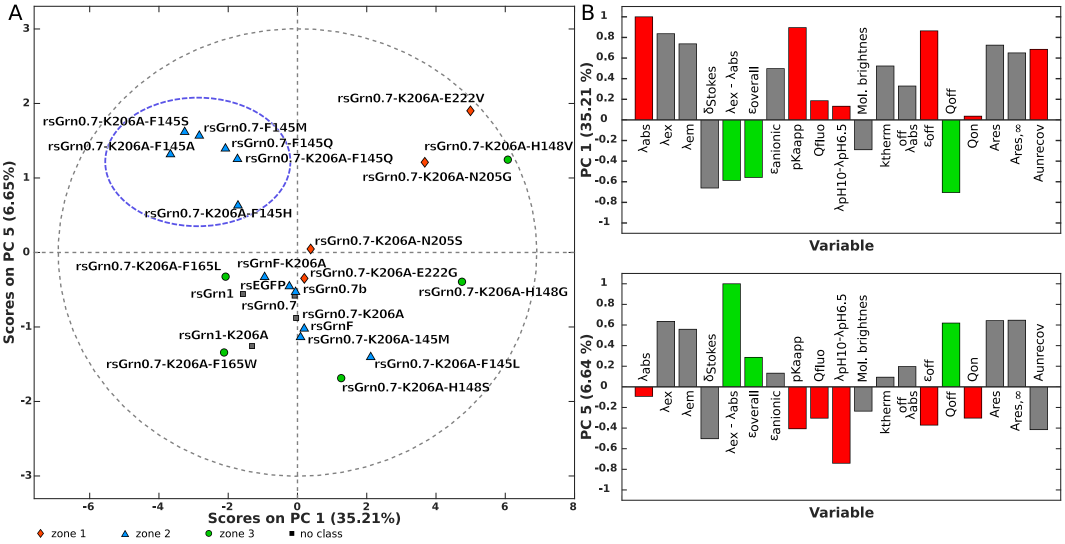


**Figure SD1:** Principal component analysis. (A) Plot showing the scores for each FP on PC1 and PC5, a combination of PCs to separate several mutants on position 145 from the other FPs (first quadrant: negative for PC1 while positive for PC5; blue dashed circle). Gray dashed circle represents the 95 % confidence level. (B) Loadings of the spectroscopic parameters on the same principal components as in A. The FPs in the first quadrant (blue dashed circle) in A are characterized by spectroscopic properties with opposite loadings in PC1 and PC5, marked by green and red bars for properties in which this group of FPs has a higher or lower value than average, respectively. Properties with the same sign in both PCs are colored grey and are not characteristic higher/lower for this group of FPs.

used. Moreover, in the on-state of rsGreen0.7 and rsGreen0.7-K206A-E222G, a double conformation of the chromophore was found (or a single one with adapted anisotropic B-factor ellipsoids can also be modeled [1]). This double conformation is not necessarily exclusive to these two FPs but might be unnoticed in the medium-resolution structures of other FPs. It is however not a general feature as it is not observed in other high-resolution structures, such as in rsGreen0.7-F145Q. The possibility of missing double conformations in the medium resolution structures is, moreover, not restricted for the chromophore but also for all other amino acids and water molecules or ligands. This is furthermore complicated by the appearance of both on- and off-state features in a single crystal. In the off-state of the RSFPs, the chromophore is stabilized through interaction with a water molecule. This water is not observed in the structures of rsGreen0.7-K206A-F145H, rsGreen0.7-K206A-F165W and rsGreen0.7-K206A-H148V. The absence of electron density for this water can be due to partial off-switching of the crystal and medium resolution X-ray diffraction data.

#### 1.3 The double conformation of Met145 of rsGreen0.7-F145M

In rsGreen0.7-F145M, Met145 has two conformations which do not superpose with the M145 side chain of rsGreen0.7-K206A-F145M. One of the conformations is in the chromophore pocket (alternative conformation A, 48% occupancy) whereas the other is outward (alternative conformation B, 52% occupancy). The electron density of the outwards conformation at the CD atom is however weak and thus also partial X-ray induced cleavage might have happened. An outwards conformation of residue 145 was already observed in the blue fluorescent protein, ECPF. [2] As we believe it is the outwards conformation that is responsible for the fast photoswitching kinetics, we only show the outwards conformation in the figures of this work.

##### 1.4 PLS-DA analysis of the FP classification according to their mutation zone

We used Partial Least Squares Discriminant Analysis (PLS-DA) for the analysis of the spectroscopic properties data set in which the FPs are classified according to their mutation zone (zone1, zone2 and zone3; Table S1). Non-classified FPs (rsGreen0.7, rsGreen0.7-K206A and rsGreen1) were excluded from this analysis.

Table S11A shows the results obtained for this PLS-DA. Ten different latent variables (LVs) were calculated, but seven LVs were recommended by looking at the average classification error of the cross-validated data. This table shows that roughly 91 % of the variance in the data points (X-Block cumulative LV) is informative to explain ~86 % of the variance in the spectroscopic properties between the different classes (Y-Block Cumulative LV). However, despite the zone 1 FPs being perfectly classified (Figure S10A left panel; everything above the red dashed line can be classified with a reasonable certainty (95 % confidence interval) to belong to that class), the prediction for the other two classes is not as good: zone 3 FPs are predicted to be part of zone 2 (Figure S10A middle panel), not all zone 2 FPs are classified as zone 2, and a bad overall classification is obtained for zone 3 FPs (Figure S10A right panel).

The fact that zone 3 FPs cannot be clearly distinguished from the others is also reflected in the summary of the PCA analysis as shown in Table 3, as no characteristic properties could be identified. However, an important aspect of this analysis is that the FPs of belonging to zone 1 could be perfectly discriminated from the other FPs. To showcase this further, the FPs of zone 3 were excluded from the analysis and PLS-DA of only zone 1 and zone 2 FP shows that one single LV (representing only ~33 % of the variance in the data points) explains ~85 % of the of the variance in the spectroscopic properties between zone 1 and zone 2 (Table S11B) and has perfect classification for the FPs, as confirmed in Figure S10B.

From this analysis, we can conclude that the used classification using the mutation zones is reasonable for at least zone 1 and 2.

### 2 Methods

#### 2.1 Mutagenesis and transformation

Site directed and site saturation mutagenesis was performed on rsGreen0.7 in a PRSETb plasmid using an modified QuikChange protocol. [3] The primers were designed using a QuikChange Primer Design tool (Agilent Technologies) and ordered from Integrated DNA Technologies (Table S12). In degenerate primers, the target codon was replaced with NNK or a triplet coding for a selective group of amino acids, found by CASTER2.0. [4] The plasmids were transformed into chemocompetent or electrocompetent JM109 (DE3) cells using sonoporation (Branson 2210 ultrasonic cleaner) or electroporation (Biorad MicroPulser Electroporator), plated on lysogeny broth (LB) agar plates supplemented with ampicillin and incubated overnight at 37 °C.

#### 2.2 *In-colony* screening

The first of two screening steps was performed on colonies on LB agar plates. A high power Xenon lamp (Max-302, Asahi Spectra) with filters of 480 nm (40 nm bandwidth) and 400 nm (30 nm bandwidth) were used to illuminate the colonies with actinic (cyan and violet respectively) and excitation light (cyan). The emitted light was collected through a 530 nm filter (40 nm bandwidth) and captured on an EMCCD camera (Cascade 512B, Photometrics). During the experiments, the LB agar plate was kept at 4 °C. The illumination and detection scheme was controlled by and the results analyzed by custom-written software in IgorPro (Wavemetrics). Using this system, colonies with deviating behavior concerning brightness and photoswitching properties were selected and were grown overnight in 1.5 mL LB-cultures in a 96 deep-well block at 37 °C.

The second screening step, was performed on an inverted microscope (Olympus IX71) coupled to a Sola Light Engine (Lumencor) and equipped with a 10x objective (UplanSApo, Olympus). Samples of the bacterial cultures were transferred into a 96 well plate and subjected to cyan and violet actinic light intermitted with cyan excitation light. The emitted light, passing through a zt488rdc dichroic mirror (Chroma), was captured on a EMCCD camera (iXon, Andor). Again the experimental control and analysis were performed using custom-written software in IgorPro.

##### 2.3 Expression and purification

Each protein was expressed from the PRSETb plasmids in JM109(DE3) cells in 1 L LB medium supplemented with ampicillin and grown at 21 °C for 4 days. Cells were harvested by centrifugation (5000 x g, 20 min), resuspended in 10xTN buffer (TN buffer: 10 mM Tris, 30 mM NaCl, pH 7.4) and lysed using a French pressure cell operating at 1150 psi. After spinning down the cell dedris (9300 x g, 20 min), the protein was purified using immobilized metal affinity chromatography (IMAC), using HisTrap HP columns coupled to an Akta Prime Plus device (both GE Healthcare Life Sciences) or manually using Ni-NTA agarose suspension (Qiagen or Macherey-Nagel) and disposable polyethylene columns (Thermo Scientific). In both cases, elution was performed using 10xTN buffer containing 500 mM imidazole. Size-exclusion chromatography was executed on a HiLoad Superdex 200-pg 16/600 column coupled to a Akta Purifier 10 system (GE Healthcare), eluting with TN or HN buffer (HN buffer: 50 mM HEPES, 30 mM NaCl, pH 7.4). Finally, the samples were concentrated using a Vivaspin turbo 10k MWCO (Sartorius).

##### 2.4 *in vitro* characterization

**$\lambda_{abs}$, $\lambda_{ex}$, $\lambda_{em}$, $\varepsilon_{overall}$, $\varepsilon_{anionic}$ and pKa.** Absorbance spectra were measured in TN buffer on a Shimadzu UV-1650PC spectrophotometer with 2.0 nm slit width. Excitation and emission spectra were recorded on a PTI Quanta-Master fluorimeter with slits set to 1.5 nm - 2.0 nm, again in TN buffer (Figure S11). Extinction coefficients were calculated using Ward's method [5] by comparing the absorbance of denatured FP in 0.1 M NaOH with the absorbance at physiological pH 7.4 ($\varepsilon_{overall}$) and at pH 9 or pH 10 where the chromophore is completely deprotonated ($\varepsilon_{anionic}$; Figure S12). The pKa titration was performed in a broad pH-range buffer (50 mM $KH_2PO_4$, 50 mM sodium citrate and 50 mM glycine), with pH ranging from 3 to 11 and the absorbance spectra were measured on a Tecan Safire 2 plate reader within 3 minutes after mixing the FP with the pH-buffer (Figure S13). The apparent pKa was determined as the average inflection point of the peak maxima of the neutral and anionic absorption band (Figure S14).

**$Q_{off}$, $Q_{on}$, $k_{therm}$, $A_{res}$, $A_{res,\infty}$ and $A_{unrecov}$** The photoswitching performance was measured on a custom-build cuvette setup, as described in Moeyaert *et al.* [6] The FP was diluted in 2 mL TN buffer to an OD below 0.15 at the protein absorption maximum around 500 nm. 488-nm (Oxius Simply Light, 16 mW) or 405-nm (Coherent CUBE, 1.8 mW) actinic light was directed from above into a 1×1 cm cuvette (Hellma). The illumination scheme contained two off- and on-switching cycli (60 × 10 s 488-nm, 20 × 10 s 405-nm) followed by a period where the bright state could thermally recover (2 × 5 min dark). The excitation and white light (Ocean Optics DT-MINI-2-GS), were coupled to the cuvette from sidewards using glass fibers. A third glass fiber coupled the cuvette to an Ocean Optics USB-4000 spectrophotometer to record the emission and transmitted light between every illumination step (Figure S15). During the measurements, the sample was kept at a temperature between 12 and 14 °C and was continuously stirred. The illumination and detection scheme was controlled by custom-written software in IgorPro.

The photoswitching quantum yield ($Q_{off}$ and $Q_{on}$) was calculated by plotting the time evolution of the anionic peak during illumination, according to a modified formula described in Moeyaert *et al.* [6] (see Moeyaert *et al.* [6] for a full description of the used symbols):

$$\mathrm{OD} = \frac{l}{l_l}\left(1 - \left(1 - e^{\frac{l}{l_l}\mathrm{OD}_0}\right) e^{-\frac{\ln(10)\varepsilon l_l}{N_A V}\frac{I_0 S\lambda}{hc}(1-\mathrm{OD_{res}})\mathrm{Qt}}\right) + \mathrm{OD_{res}} \quad (1)$$

The rate constant of thermal recovery at a certain temperature, $k_{therm}$ can be calculated using

$$\mathrm{k_{therm}} = \ln\left(-\frac{\mathrm{OD_{max}} - \mathrm{OD_{res}}}{(\mathrm{OD_{therm}} - \mathrm{OD_{res}}) - (\mathrm{OD_{max}} - \mathrm{OD_{res}})}\right)/\mathrm{t_{therm}} \quad (2)$$

with $OD_{max}$, $OD_{res}$ and $OD_{therm}$ being the optical density (OD) of the on-state, the residual OD after off-switching and the OD after the period of thermal recovery $t_{therm}$, respectively.

The residual absorbance at infinite laser power ($A_{res,\infty}$), represents those experimental settings in which a much higher laser power is applied than in the *in vitro* measurement, e.g. during microscopy. In such experiments thermal recovery can be ignored:

$$A_{res,\infty} = \frac{k_{on,488}}{k_{tot,inf\ power}} = \frac{k_{on,488}}{(k_{on,488} + k_{off,488})} \quad (3)$$

in contrast to the residual signal during weak laser power that was used during the *in vitro* measurements ($A_{res}$):

$$A_{res} = \frac{k_{on,488}}{k_{tot}} = \frac{k_{on,488} + k_{therm}}{k_{on,488} + k_{off,488} + k_{therm}} \quad (4)$$

with taking $k_{therm}$ from equation 2. The total rate constant of off-switching ($k_{tot}$) is the negative reciprocal argument of the mono exponential decay of the OD during off-switching, or

$$k_{tot} = -\frac{\ln(10)\varepsilon l_l}{N_A V}\frac{I_0 S \lambda}{hc}(1 - OD_{res})$$

In practice, $A_{res,\infty}$ can be calculated as

$$A_{res,\infty} = \frac{OD_{res} \times k_{tot} - k_{therm}}{k_{tot} - k_{therm}} \quad (5)$$

The parameters $Q_{off}$, $Q_{on}$, $A_{res}$ and $A_{res,\infty}$ are calculated from the second photoswitching cycle. The first photoswitching cycle often behaves different as a decent amount of signal does not recover to the bright state. The unrecoverable signal ($A_{unrecov}$) is a measure for this loss:

$$A_{unrecov} = \frac{OD_{0,first} - OD_{0,second}}{OD_{0,first}} \quad (6)$$

with $OD_{0,first}$ and $OD_{0,second}$ respectively being the OD before and after the first photoswitching cycle.

**$Q_{fluo}$ and molecular brightness** Also on the custom-build cuvette setup, the quantum yield of fluorescence ($Q_{fluo}$) could be determined by quasi-simultaneously measuring the absorbance and emission spectrum, relative to rsEPGF ($Q_{fluo}$=0.42 [7]).

The molecular brightness was calculated as the product of the overall extinction coefficient at physiological pH (see above) and the quantum yield of fluorescence.

**$\varepsilon_{off}$** The extinction coefficient of the off-state ($\varepsilon_{off}$) was calculated based on the known extinction coefficient at $\lambda_{abs}$ of the on-state ($\varepsilon_{overall}$) and the absorption at $\lambda_{abs}^{off}$ after off-switching.

#### 2.5 Crystallization

Crystals were grown at 16 or 20 °C in sitting and/or hanging drops, setup manually or automatically (Evo Freedom (Tecan), Mosquito (ttplabtech) and Crystal Gryphon (Art Robins Instruments)). The crystals for the structure determination of the bright state were soaked in cryoprotectant and frozen in liquid nitrogen. A part of the crystals were illuminated with 473- or 491-nm light to bring them in the off-state. Afterwards, they were also soaked in the cryoprotectant solution and subsequently frozen in liquid nitrogen. All crystals were maintained in liquid nitrogen until data collection. The used crystallization cocktail and cryoprotectant solution for each of the proteins can be found in Table S13. The crystal structures of rsGreen0.7 in the green-on and green-off state were determined and published by us in Duwé *et al.* [1].

#### 2.6 Structure determination

X-ray diffraction data were acquired at PILATUS or EIGER detectors at various beamlines (Table S2). The data were processed using XDS, [8] and scaled and merged using Aimless. [9] Only if the data had a resolution of 2.5 Å or better, it was considered as suitable for this study. The structures were solved via molecular replacement with Phaser [10] or by rigid body refinement in Phenix.refine [11] in case of isomorfism with the phasing model. Phasing models were rsGreen0.7 on- or off-state (resp. PDB ID: 4XOW and 4XOV),[1] rsGeen0.7-K206A on- or off-state, or the corresponding on-state structure to phase an off-state structure. The subsequent likelihood-based refinement was carried out using Phenix.refine and manual model manipulations were performed in COOT. [12]

After molecular replacement, the chromophore was modelled in the mFo-DFc difference map and mutations compared with the phasing model were carried out. Water molecules and ligands (compounds present in the crystallization cocktail) were included in the model in case they were within hydrogen bond distance to chemical relevant groups, appeared in mFo-DFc difference map at 3 r.m.s.d., maintain in the 2mFo-DFc electron density of 1 r.m.s.d. and had reasonable B-factors. The chromophore and ligand dictionary files were created using eLBOW [13] and were manually corrected if needed. For structures in which both on-and off-state features were observed (both *cis* and *trans*-configuration of the chromophore and/or the out- and inwards conformation of His148), alternative conformation C was annotated to non-investigated state (e.g. *cis*-chromophore in an off-state structure was annotated as alternative conformation C). In case of high resolution, riding hydrogens were added and B-factors were refined anisotropically. Except for the chromophore, which we kept anionic in the *cis* state and neutral in the *trans* state, the protonation states of the other residues were automatically annonated and might not represent the actual protonation state.

Data collection and refinement details and statistics can be found in Table S2.

#### 2.7 Quantitative structural information

The crystal structures are quantitated on five levels (Table S3):
(i) Internal descriptors of the chromophore: methylene bridge bond angle, and tilt and twist torsion angles[14] (Figure S9A).
(ii) Distances between residues or water molecules and chromophore anchor points (hydroxybenzylidene oxygen, hydroxybenzylidene plane center, imidazolinone oxygen and imidazolinone nitrogen, Figure S9B) if they were smaller than 3.5 Å in rsGreen0.7. Only those distances of which the total range over the different FPs was at least 0.5 Å were retained and therefore the descriptors concerning the on- and off-state structures might be different. See Table S3 for the specific atoms of the residues used to calculate these distances.
(iii) Interresidue distances of the residues surrounding the chromophore within a sphere of 3.5 Å. Only those distances of which the total range over the different FPs was at least 0.5 Å were retained and therefore the descriptors concerning the on- and off-state structures might be different. See Table S3 for the specific atoms of the residues used to calculate these distances.
(iv) The angle between the Phe145 or Phe165 ring plane and the hydroxybenzylidene ring plane.
(v) Other properties of the chromophore pocket: first, the chromophore pocket volume was calculated using the CASTp server [15] using the crystal structures without chromophore. In the off-states of rsGreen0.7-F145M and rsGreen0.7-K206A, F145 and H148, respectively, are not completely modelled. The chromophore pocket calculation of these FPs can therefore be slightly overestimated although the absence of clear electron density indicates that these residues are very flexible. In rsGreen0.7-K206A-H148G, the $\beta$-barrel is open and no closed pocket could be defined by CASTp. Second, the amount of atoms or heteroatoms within a radius of 3.5 Å around the chromophore or a chromophore anchoir point. Finally, the number of hydrogen bonds established by the chromophore heteroatoms with hydrogen atom or free electron pair, as described in De Zitter *et al.* [16]

Unless stated differently above, all calculations were performed through Pymol v. 2.0. [17] In case of multiple conformations, only the conformation named alternative conformation A, was taken into account. In case of multiple monomers in the asymmetric unit, chain A was used, except for rsGreen0.7-K206A-H148S where chain C was used.

#### 2.8 Principal component and correlation analysis

Principal Component Analysis (PCA) was performed using the Eigenvector Research, Inc. PLS_Toolbox. The data were ordered in a matrix where rows represent observations (i.e. FPs) and columns the spectroscopic/structural properties. Before performing the PCA, all data were autoscaled, because the spectroscopic properties and structural descriptors cover different numerical ranges. As data with a large range has a large variance and data with a small range has a smaller variance, the PCA (which is a maximum variance projection method) will more likely express the variables with large variance in the model compared to the low-variance variables. By autoscaling, all variables are unit-variance scaled and the bias towards large values is removed. Mean-centering of the data was also performed to improve interpretability of the model. Missing data points were included by the 'best guess' method, which uses the PCA model to replace the missing data points and repeats this until convergence of the projected values for the missing data points. For more information about PCA in general, we refer to Bro and Smilde. [18]

Partial Least Squares-Discriminant Analysis (PLS-DA) was used to check whether the zone1/zone2/zone3 sample grouping was reasonable and for the analysis of fast/medium/slow switchers in the data set containing spectroscopic and structural descriptors associated with the off-state structures. It can be seen as an advanced version of PCA with rotated PCs (i.e. latent variables) to maximize the differences between the predefined classes, in order to understand whether the data carries information that allows separating the different sample classes. All PLS-DA analyses were performed using a Venetian blinds cross-validation method for a maximum number of 10 latent variables (LVs). The two positive switchers, rsGreen0.7-K206A-N205C/L, were retained from the PLS-DA with the spectroscopic properties. The two rsGreen0.7-K206A-H148 mutants were, due to their altered chromophore off-state conformation, left out of the PLS-DA with the spectroscopic and structural properties. The PLS-DA analyses were performed using the the Eigenvector Research, Inc. PLS_Toolbox.

For visual interpretation we colored the scores according to sample classes (i.e. zone 1, zone 2 and zone 3). When including the structural descriptors, the number of variables and thus the number of loadings becomes large. To aid the loadings interpretation we also colored the loadings according to variable groups. We classified the structural descriptors as *internal* (group (i) properties), *pocket* (group (v) properties), *hydroxybenzylidene* (group (ii) and (iv) properties), *imidazolinone* (group (ii) properties), or as the associated residue (group (ii), (iii) and (iv) properties; See above, Supplementary Table S3). Whenever two partners are involved in a descriptor, the descriptor will belong to two groups, giving raise to two possible coloring schemes, e.g. the distance $A150_{CB}$ – $hydroxybenzylidene_{OH}$ will belong to group *A150* and *hydroxybenzylidene* whereas 'chromophore pocket volume' will only belong to the *pocket* classifier and have the same color in the two color groupings. This is why in Figures S4,S5and S6 the loadings are depicted twice, once according to each variable group.

### 3 Supporting tables and figures

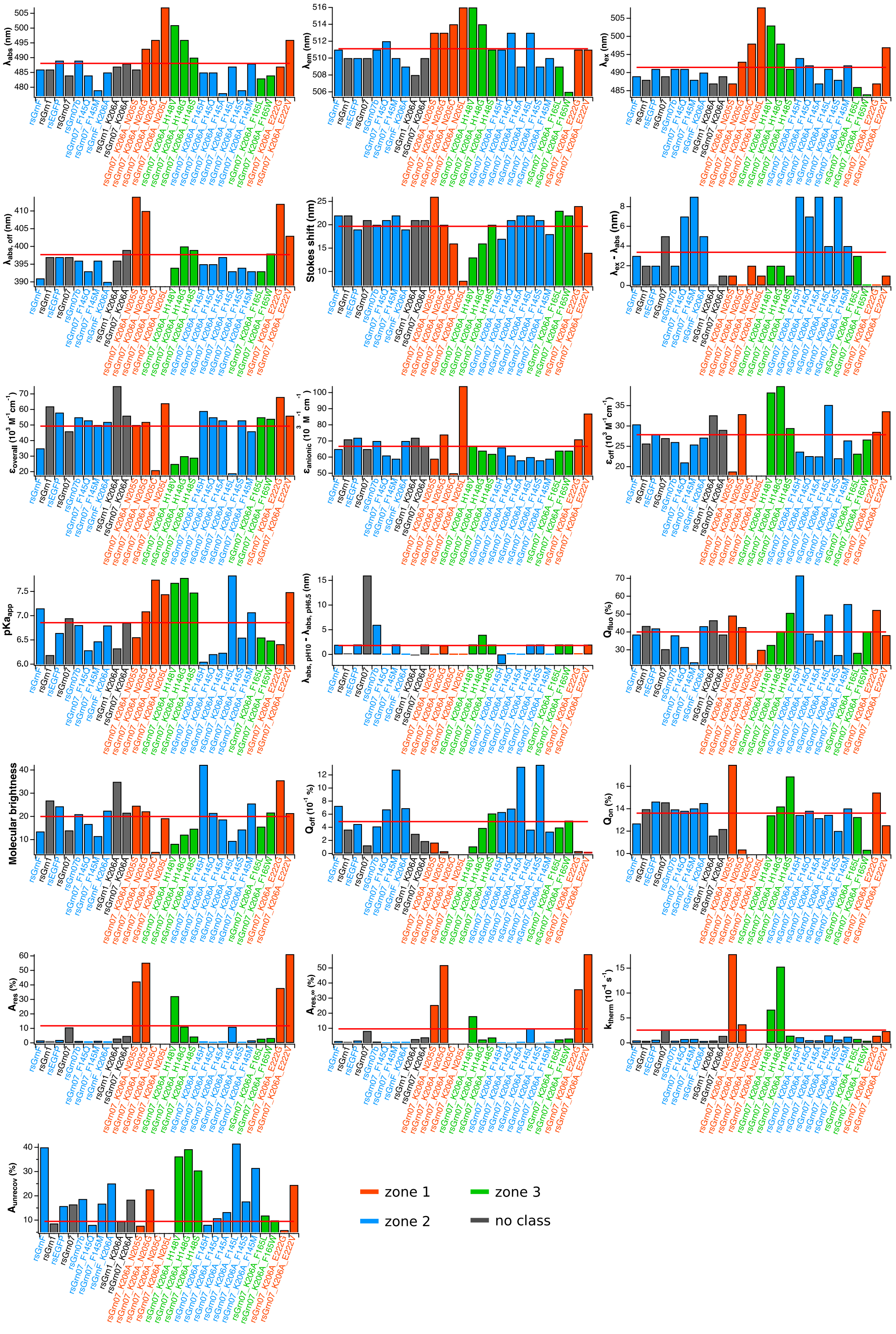


**Figure S1:** Bar chart with distribution of all spectroscopic parameters, colored according to classification of the FPs. Horizontal red line indicates the average value for each property.

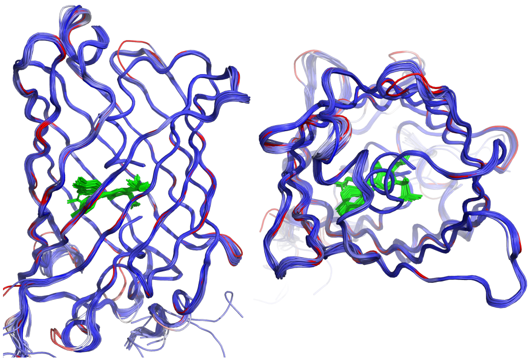

**Figure S2:** Front and top view of superposition of rsEGPF and the rsGreen-variants, colored according to relative distance from rsGreen0.7 on-state (blue to red: 0 – 5 Å and up). The chromophore is depicted in green sticks.

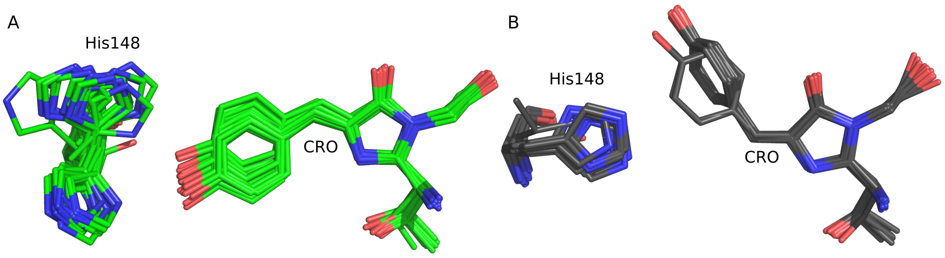


**Figure S3:** Superposition of the chromophore and H148 of rsEGPF and the rsGreen-variants (A) in the on-state and (B) in the off-state, showing a different chromophore off-state conformation for the H148 mutants.

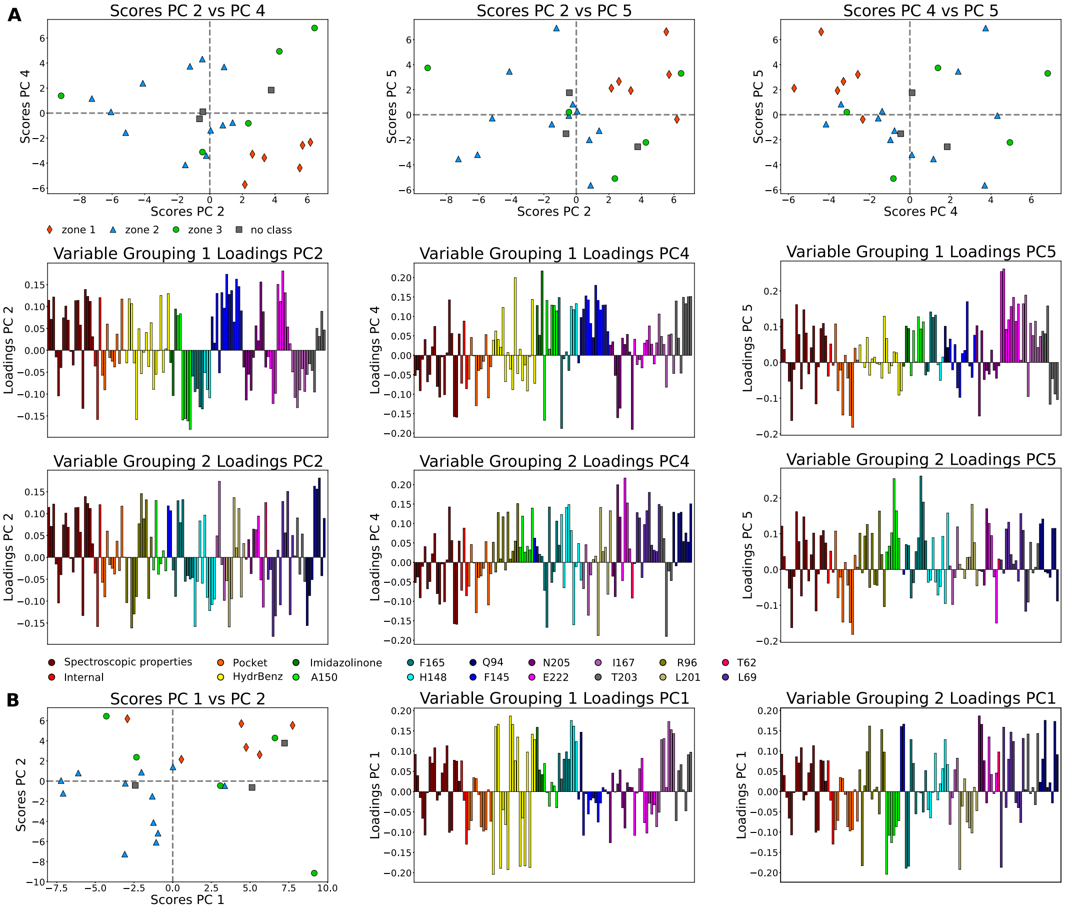


**Figure S4:** PCA of the spectroscopic and on-state structural properties showing how to isolate zone 1 and zone 2 mutants. (A) Score plots with FPs colored according to their mutation zone (top), and loading plots colored according to their groups (middle and bottom; see Materials and Methods section for variable groups) for PC2, PC4 and PC5 in which the zone 1 mutants can be separated from the other FPs. Properties with positive values in PC2 and PC5, and negative in PC4 will be higher for the zone 1 mutants compared to average, whereas the zone 1 mutants will have lower values for properties with negative values in PC2 and PC5, and positive in PC4 (Table S7). (B) PCA scores (left) and loadings (middle and right; loadings for PC2 can be found in (A)) colored as in (A) for PC1 and PC2 that can separate the zone 2 mutants from the other FPs. The zone 2 mutants are negative in PC1 and PC2, consequently variables with positive loadings associated with PC1 and PC2 will be lower for the zone 2 mutants whereas negative loadings in both FPs indicate properties for which the zone2 mutants have higher values.

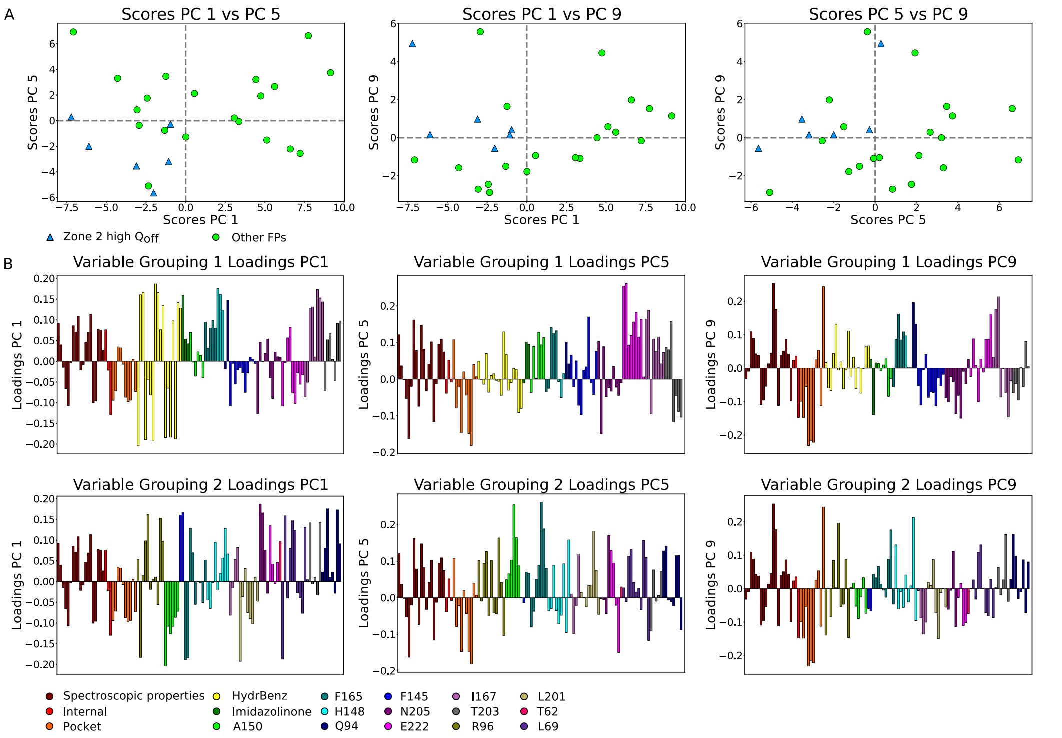


**Figure S5:** PCA of the spectroscopic and on-state structural properties showing how the fast switching FPs can be isolated from the other FPs using principal component 1, 5 and 9. (A) Score plots and (B) loading plots colored and clustered according to their group (see Table S1 and Materials and Methods section for FP and variable groups. respectively). The loadings with negative values in PC1, PC5 and positive in PC9 can be attributed to characteristic properties of the fast switchers.

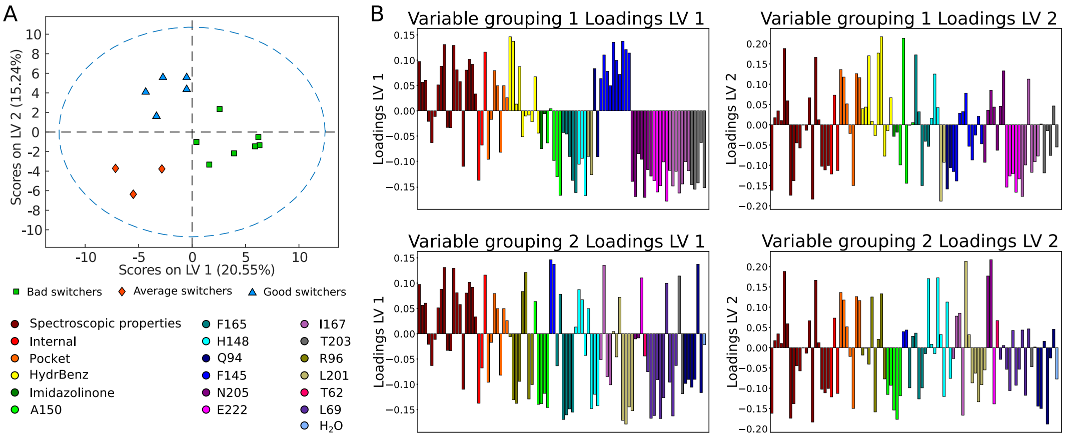


**Figure S6:** PLS-DA of the spectroscopic and off-state structural descriptors showing how fast and slow switching FPs can be separated from each other using LV1 and LV2. (A) Score plots and (B) loading plots colored and clustered according to their groups (see Table S1 and Materials and Methods section for FP and variable groups. respectively). The loadings with negative values in LV1and positive in LV2 can be attributed to properties for which the fast (slow) switchers will have higher (lower) values than average.

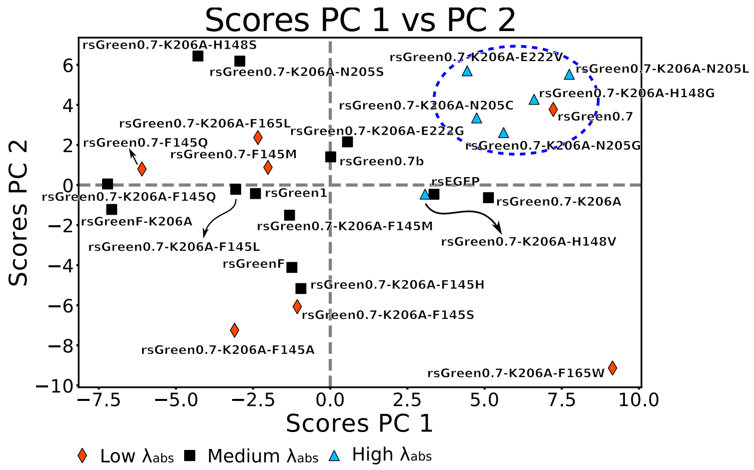


**Figure S7:** PCA showing a weak isolation of FPs with red-shifted absorption maximum by PC1 and PC2 (blue dashed circle).

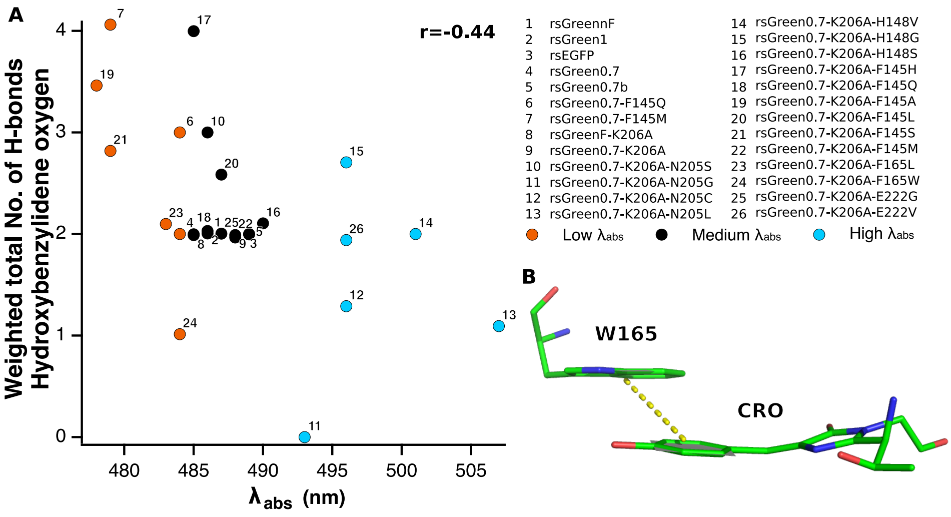


**Figure S8:** (A) Negative correlation between $\lambda_{\text{abs}}$ and number of potential hydrogen bonds made by the chromophore benzylidiene oxygen. (B) Interaction between W165 and the hydroxybenzylidene ring disturbs chromophore planarity in rsGreen0.7-K206A-F165W.

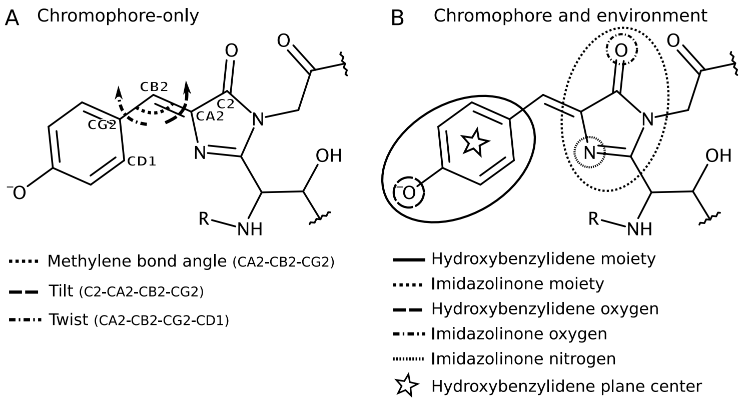


**Figure S9:** Illustration of the internal chromophore descriptors (A) and chromophore anchor points (B). Note that for simplicity we refer to the benzylidene plane center as "hydroxybenzylidene plane center".

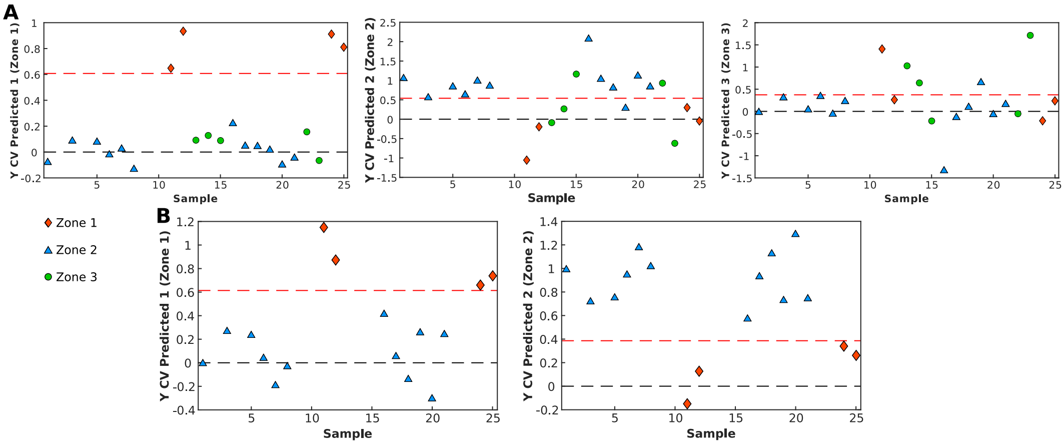


**Figure S10:** PLS-DA of the spectroscopic properties with classification of the FPs according to the mutation zone (Table S1). (A) FPs classified according to zone1, zone2 and zone3. (B) Only FPs classified as zone1 and zone2. Black and red dashed line: 0 cross-validation (CV) and 95% confidence region.

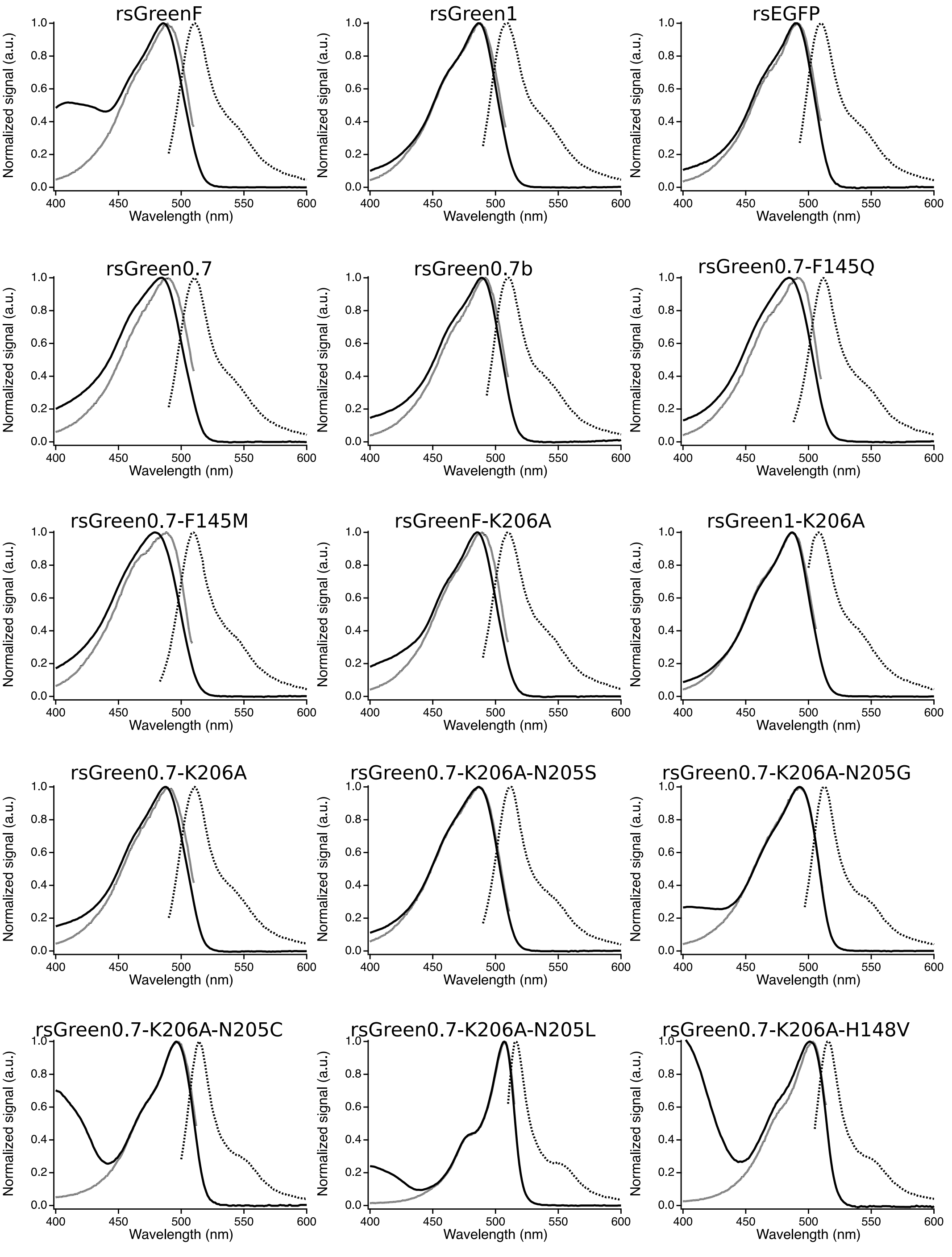


**Figure S11:** Absorption spectra (full line), excitation spectra (dotted line) and emission spectra (dashed line) for rsEGFP and the rsGreen-variants.

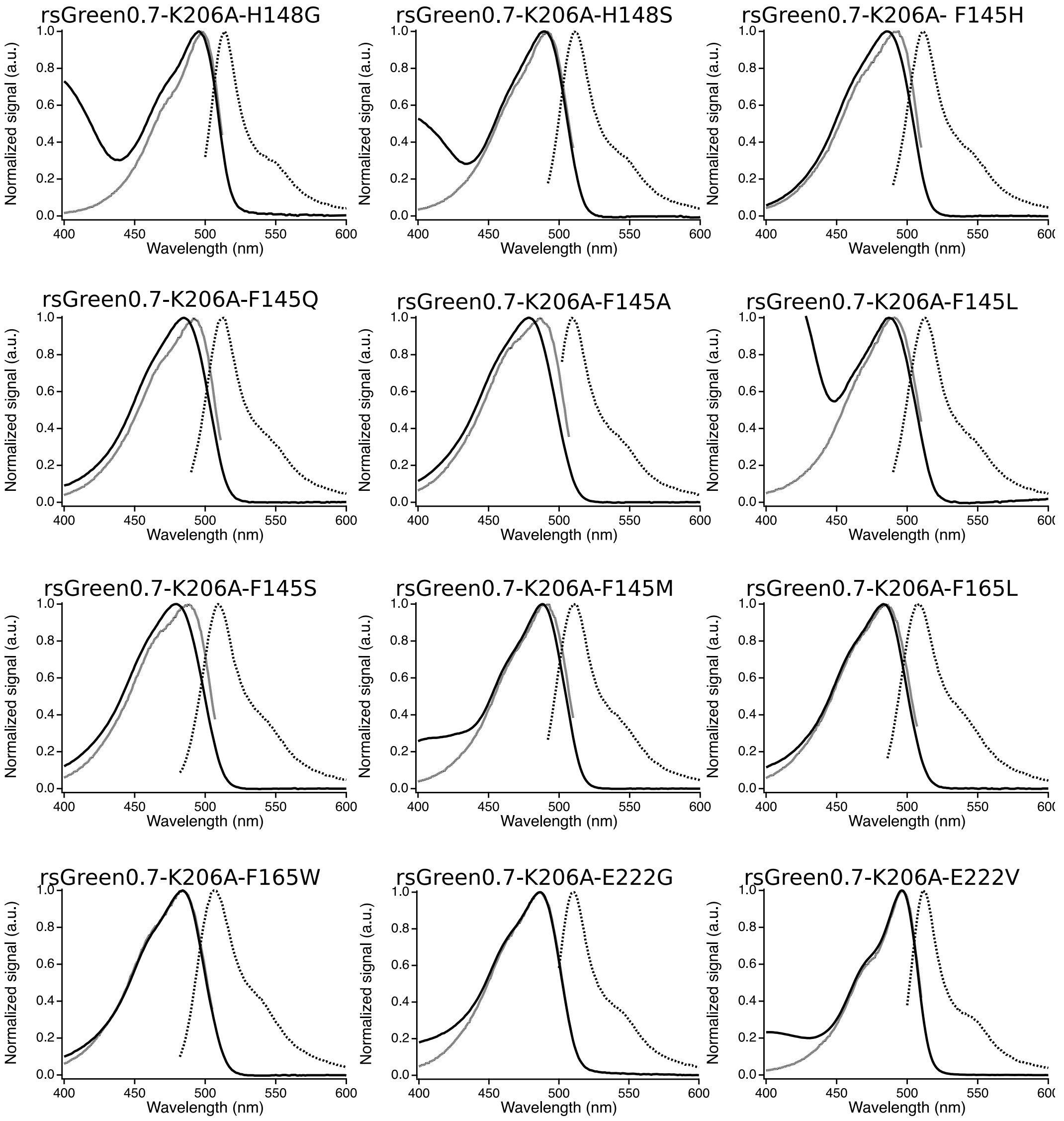


*(Continued)* Absorption spectra (full line), excitation spectra (dotted line) and emission spectra (dashed line) for rsEGFP and the rsGreen-variants.

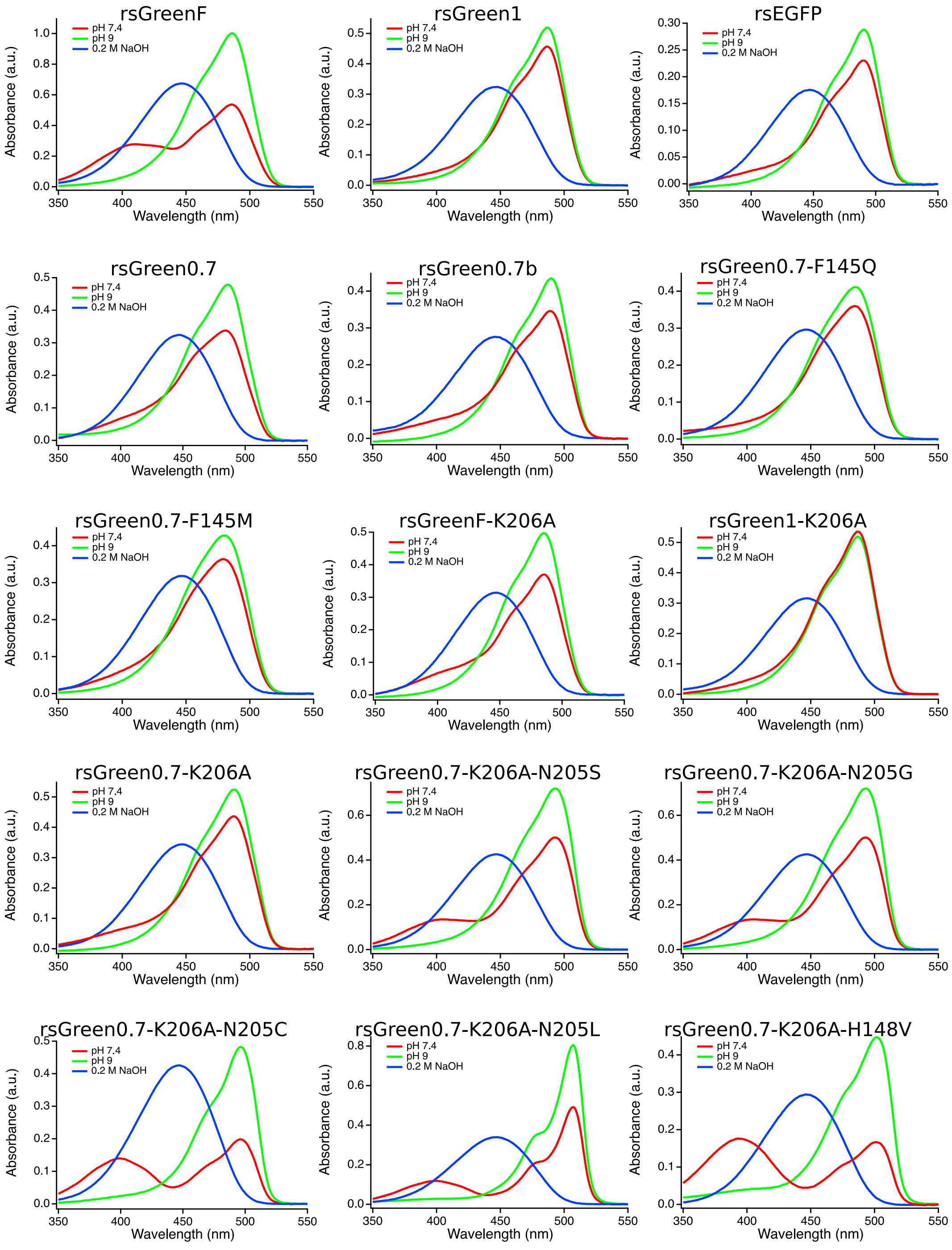


**Figure S12:** Absorption spectra at physiological pH (red), high pH (green) and of the NaOH-induced denatured protein (blue) for rsEGFP and the rsGreen-variants.

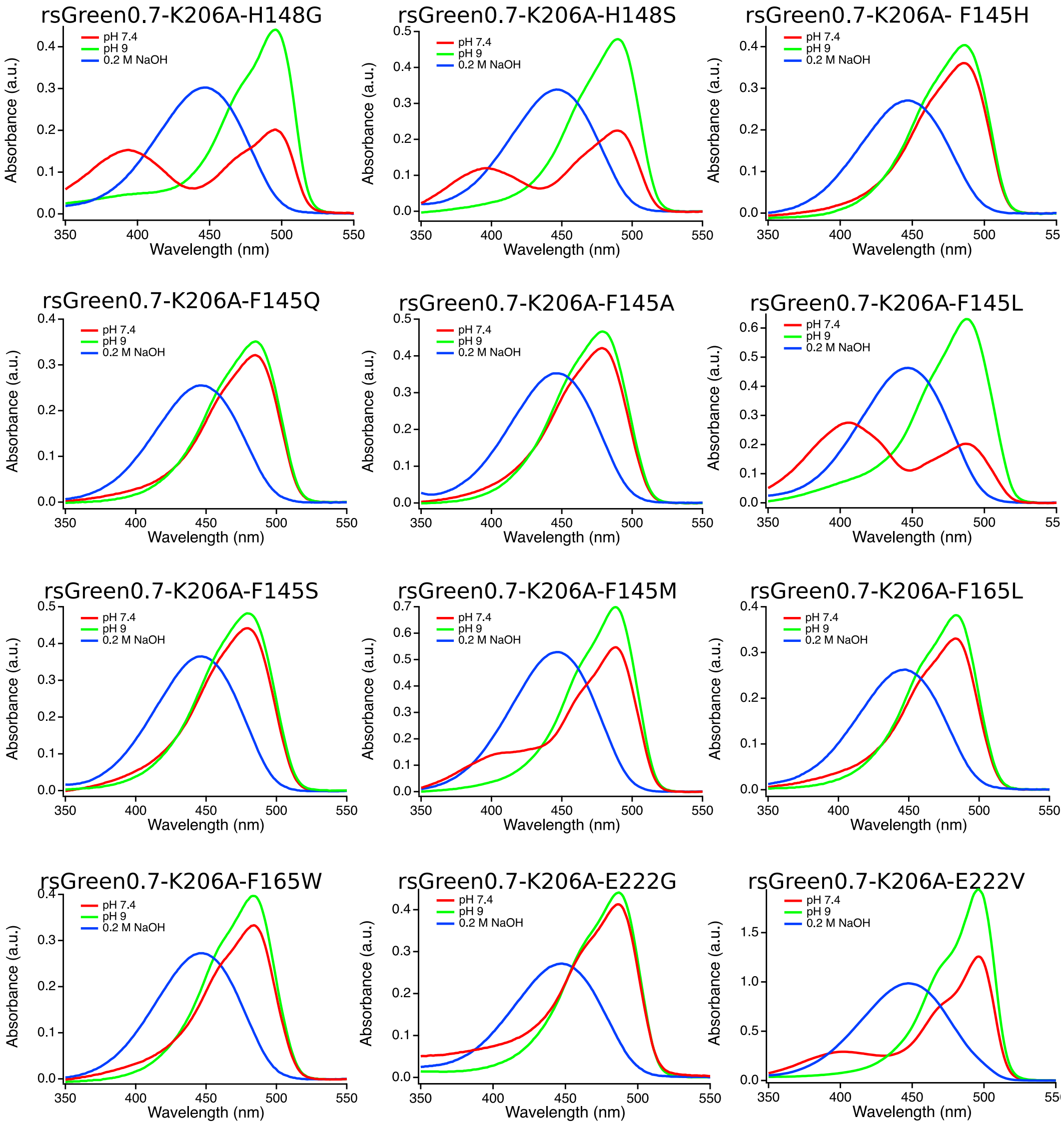


*(Continued)* Absorption spectra at physiological pH (red), high pH (green) and of the NaOH-induced denatured protein (blue) for rsEGFP and the rsGreen-variants.

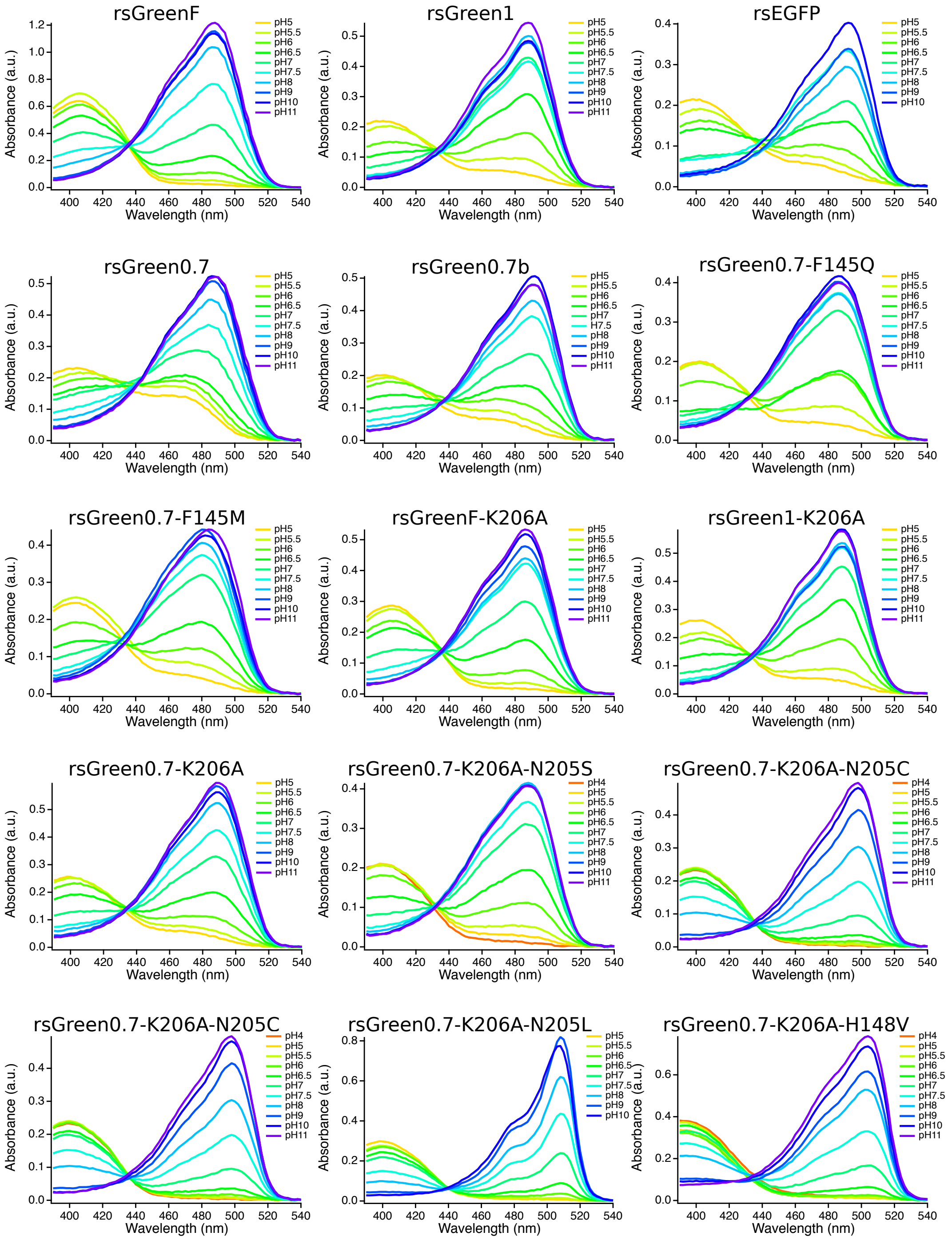


**Figure S13:** Absorption spectra during the pH titration for rsEGFP and the rsGreen-variants.

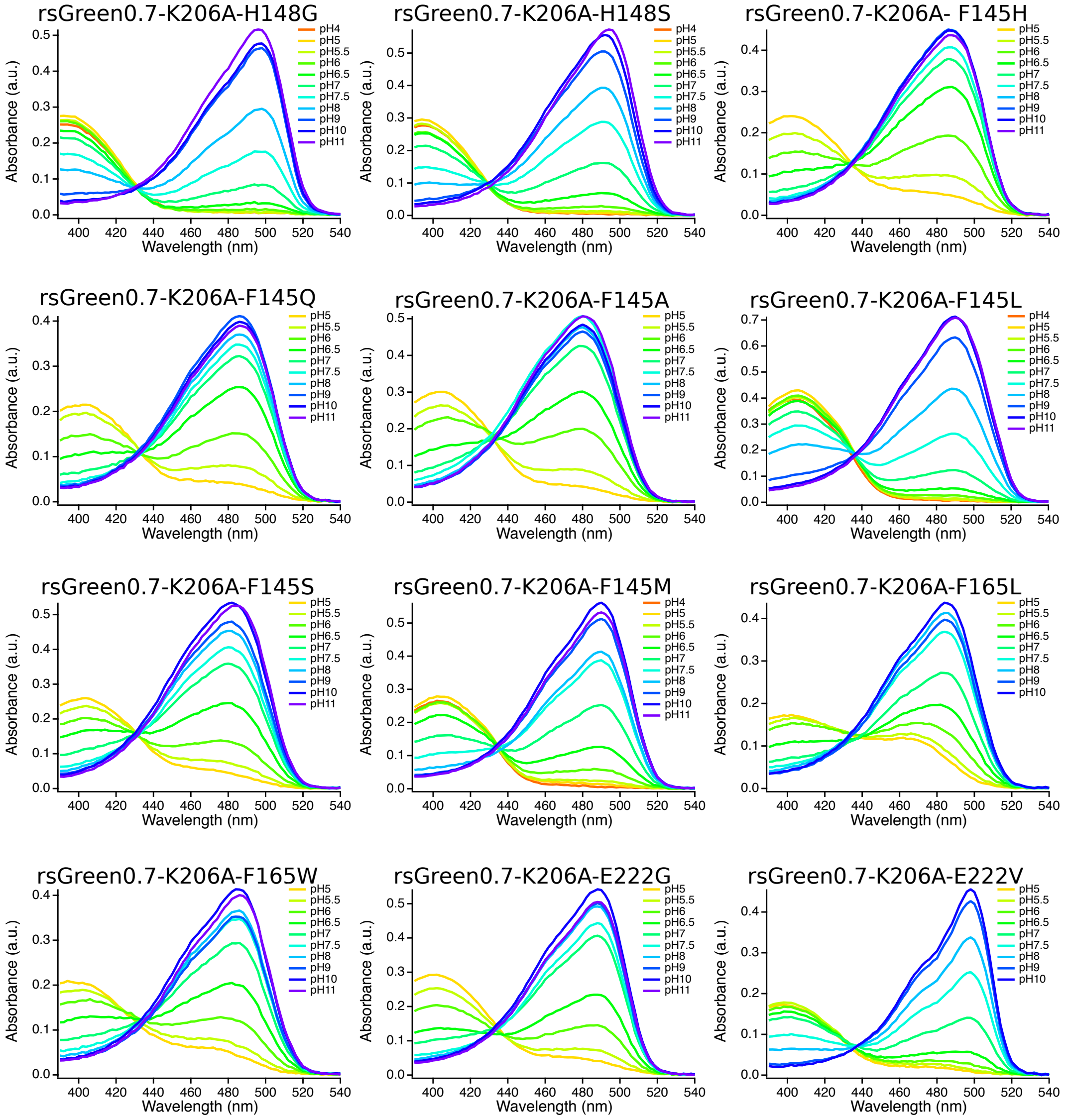


*(Continued)* Absorption spectra during the pH titration for rsEGFP and the rsGreen-variants.

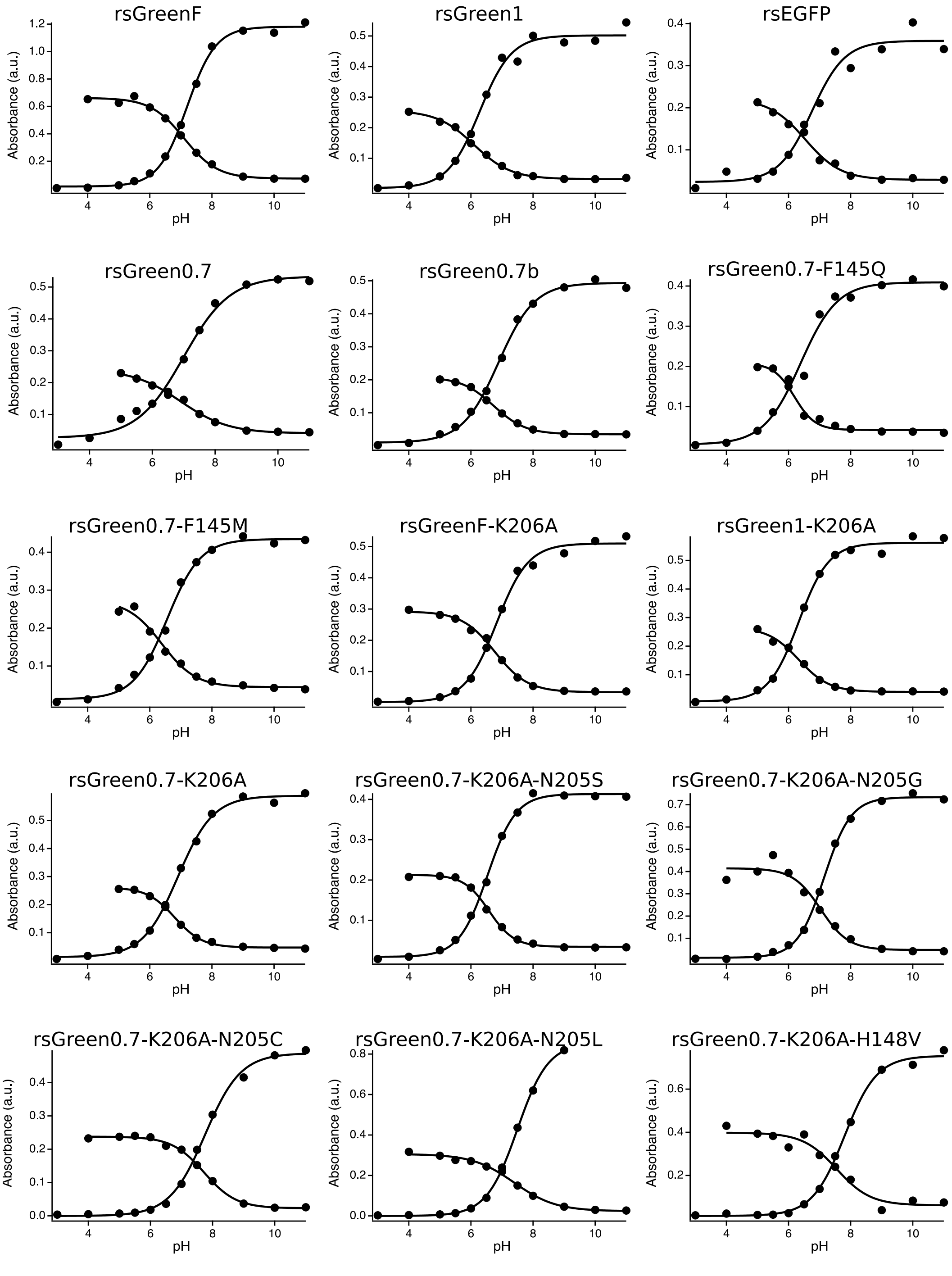


**Figure S14:** Absorption maxima of the anionic and neutral peak in function of the pH for rsEGFP and the rsGreen-variants.

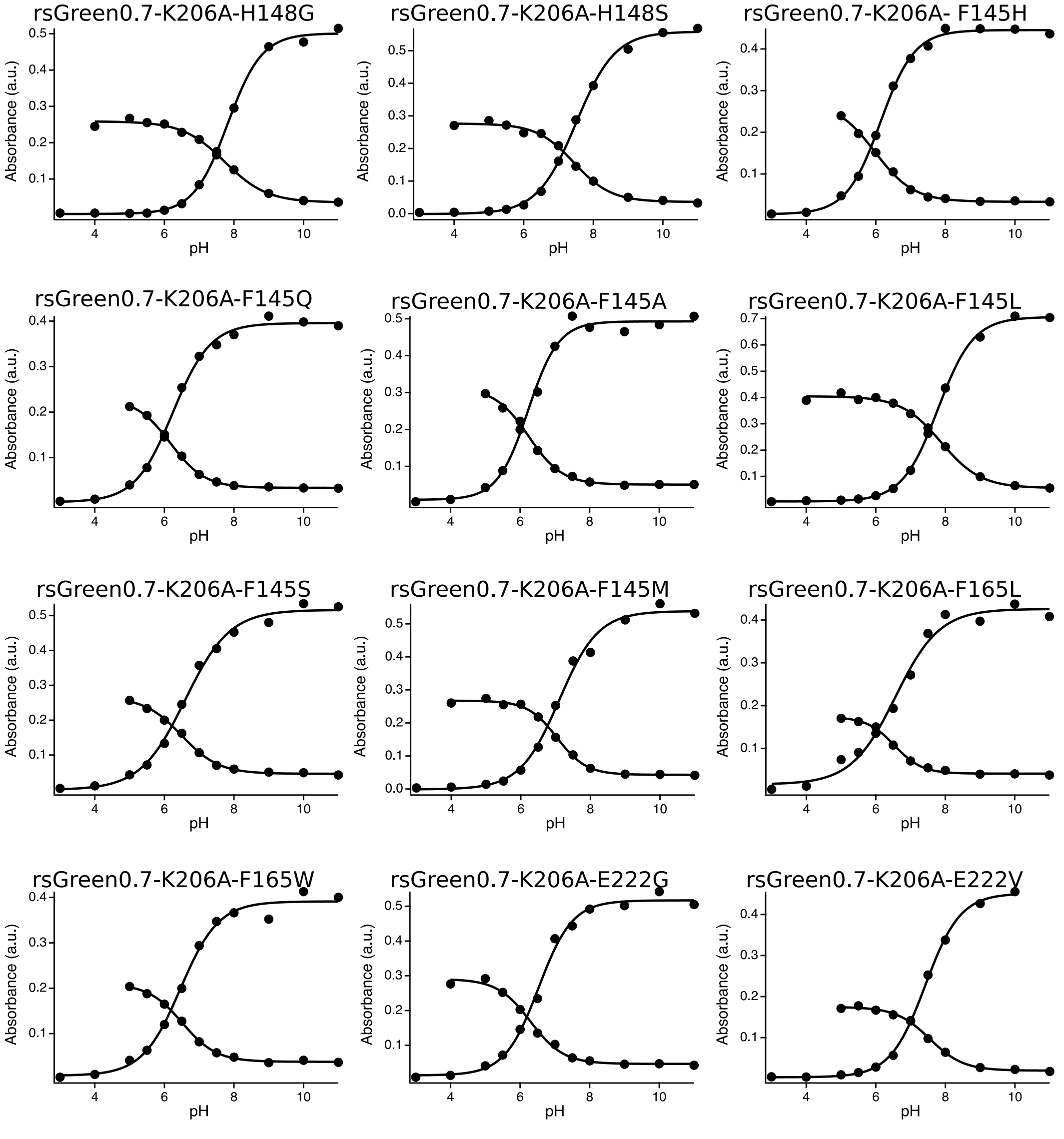


*(Continued)* Absorption maxima of the anionic and neutral peak in function of the pH for rsEGFP and the rsGreen-variants.

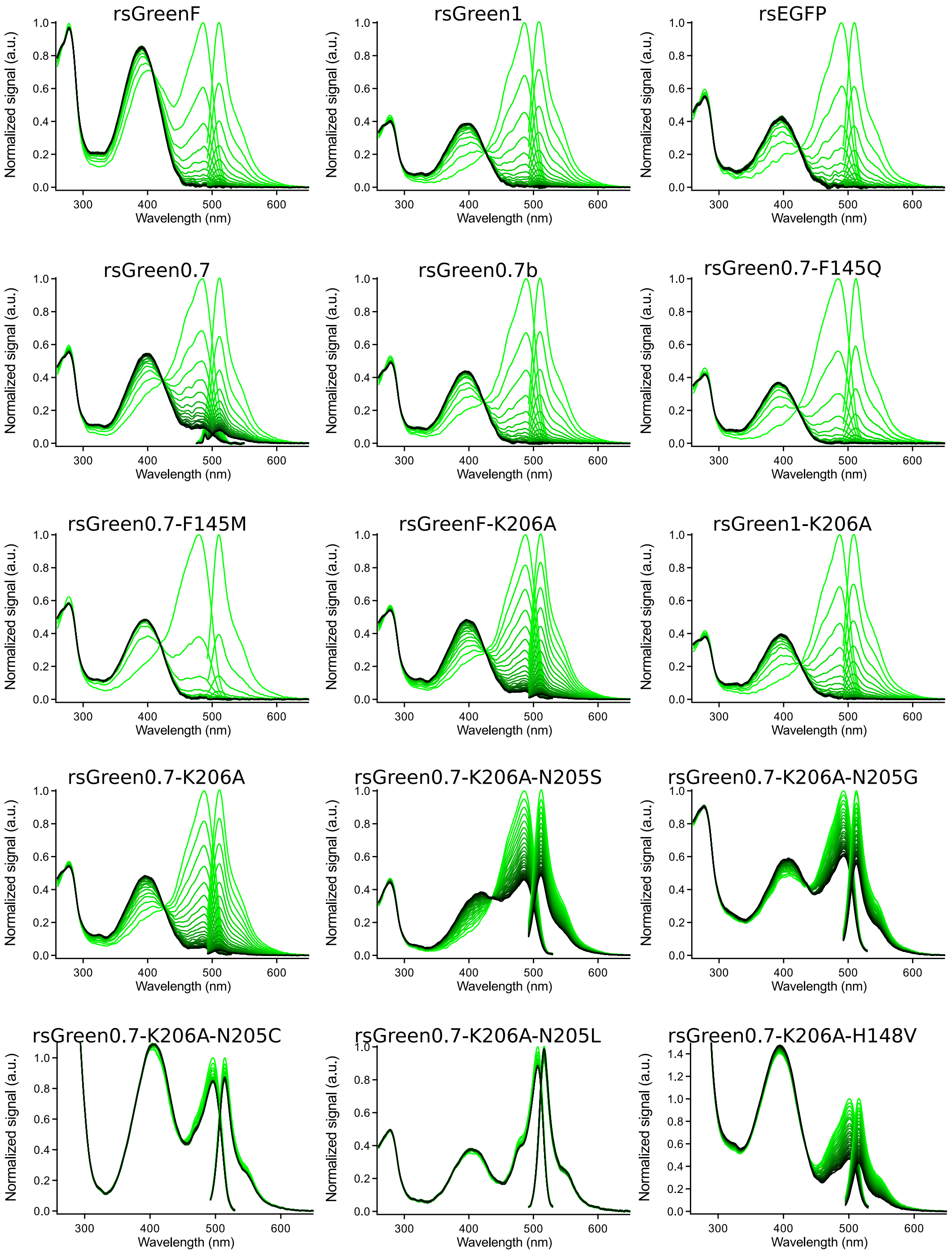


**Figure S15:** Absorption and emission spectra during illumination with 488-nm light for rsEGFP and the rsGreen-variants (from green to black).

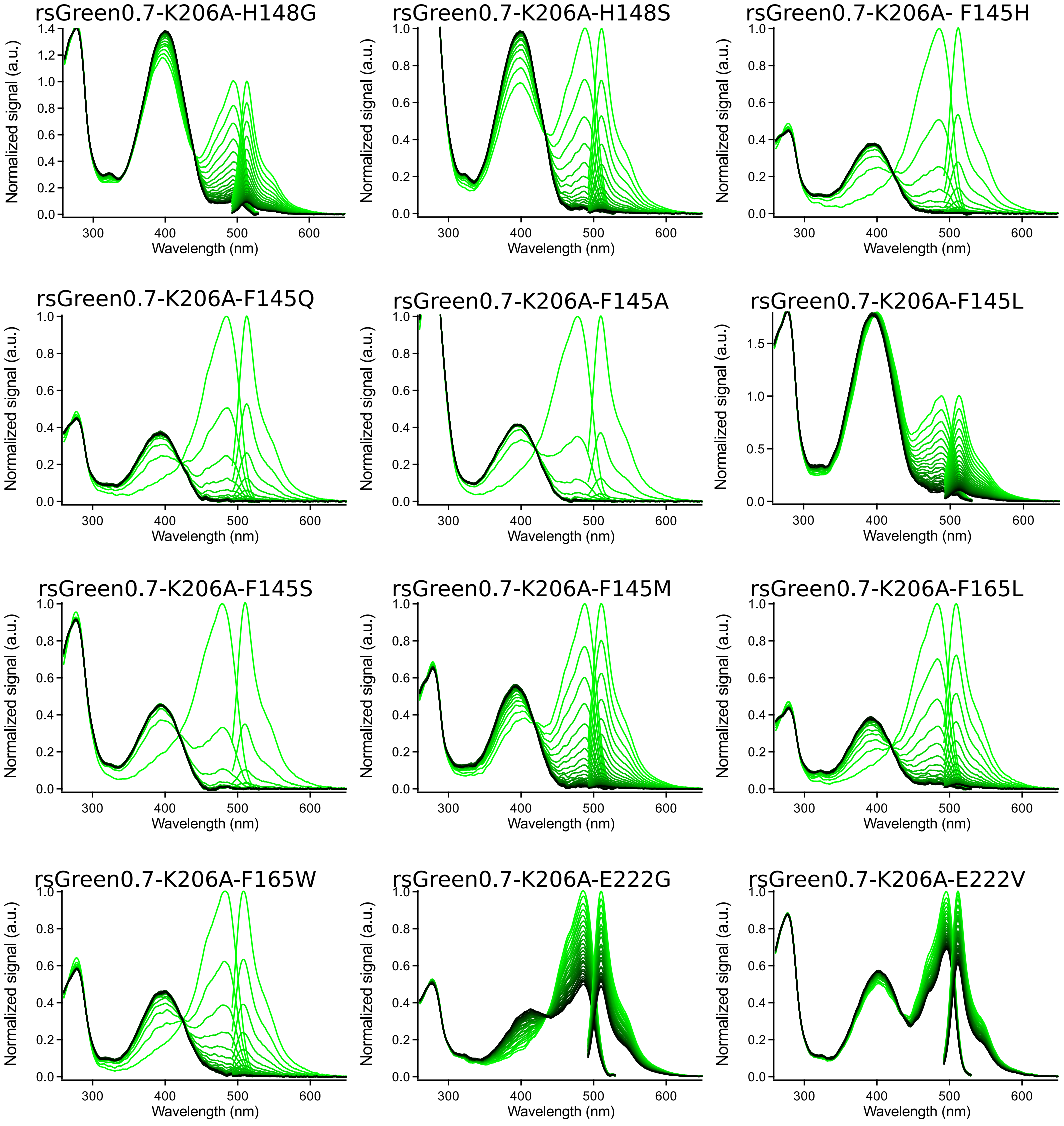


*(Continued)* Absorption and emission spectra during illumination with 488-nm light for rsEGFP and the rsGreen-variants (from green to black).

**Table S1:** FP classification for PCA and state in which the structure was determined.

| Protein | Zone | Switching$^{\$}$ | State of crystal structure(s) |
|---|---|---|---|
| rsGreenF | 2* | medium | on and off |
| rsGreen1 | § | slow | on |
| rsEGFP | 2* | slow | on and off |
| rsGreen0.7 | § | slow | on and off† |
| rsGreen0.7b | 2 | slow | on and off |
| rsGreen0.7-F145Q | 2 | fast | on |
| rsGreen0.7-F145M | 2 | fast | on and off |
| rsGreenF-K206A | 2 | medium | on and off |
| rsGreen1-K206A | § | slow | off |
| rsGreen0.7-K206A | § | slow | on and off |
| rsGreen0.7-K206A-N205S | 1 | slow | on |
| rsGreen0.7-K206A-N205G | 1 | slow | on |
| rsGreen0.7-K206A-N205C# | 1 | § | on |
| rsGreen0.7-K206A-N205L# | 1 | § | on |
| rsGreen0.7-K206A-H148V | 3 | slow | on and off |
| rsGreen0.7-K206A-H148G | 3 | slow | on |
| rsGreen0.7-K206A-H148S | 3 | medium | on and off |
| rsGreen0.7-K206A-F145H | 2 | fast | on and off |
| rsGreen0.7-K206A-F145Q | 2 | fast | on and off |
| rsGreen0.7-K206A-F145A | 2 | fast | on and off |
| rsGreen0.7-K206A-F145L | 2 | slow | on and off |
| rsGreen0.7-K206A-F145S | 2 | fast | on and off |
| rsGreen0.7-K206A-F145M | 2 | slow | on and off |
| rsGreen0.7-K206A-F165L | 3 | slow | on |
| rsGreen0.7-K206A-F165W | 3 | medium | on and off |
| rsGreen0.7-K206A-E222G | 1 | slow | on |
| rsGreen0.7-K206A-E222V | 1 | slow | on |

$ fast: occupying upper left quadrant of Figure SD1, medium: around or above average but not included in 'fast'; slow: below average. (Figure S1)
* Residue 145 was used for classification.
§ No classification possible.
† Structures determined in Duwe *et al.* [1]
#rsGreen0.7-K206A-N205C and rsGreen0.7-K206A-N205L are possitive switchers and were therefore not used in the PCA or PLS-DA.

**Table S2:** Data collection, processing and refinement statistics.

| | rsGreenF Green-on | rsGreenF Green-off | rsEGFP Green-on | rsEGFP Green-off | rsGreen0.7b Green-on | rsGreen0.7b Green-off |
|---|---|---|---|---|---|---|
| PDB ID | 7A7S | 7A7T | 7A7K | 7A7L | 7A7M | 7A7N |
| Beamline | Soleil, Proxima2A | Soleil, Proxima2A | Soleil, Proxima1 | Soleil, Proxima1 | Soleil, Proxima1 | Elettra, XRD1 |
| Wavelength (Å) | 0.98 | 0.98 | 0.89 | 0.98 | 0.89 | 1.00 |
| Space group | $P2_1$ | $P2_1$ | $P2_1$ | $P2_12_12_1$ | $P2_12_12_1$ | $P2_12_12_1$ |
| Unit cell parameters | | | | | | |
| a, b, c (Å) | 37.33 61.97 46.78 | 37.62 62.26 48.46 | 47.48 51.4 47.66 | 51.11 62.00 71.21 | 32.02 61.19 105.67 | 31.87 61.57 105.12 |
| $\alpha$, $\beta$, $\gamma$ (°) | 90.00 91.71 90.00 | 90.00 90.36 90.00 | 90.00 97.76 90 | 90.00 90.00 90.00 | 90.00 90.00 90.00 | 90.00 90.00 90.00 |
| Resolution range (Å) | 46.76 – 1.73 | 48.46 – 1.58 | 47.19 –1.55 | 46.76 – 1.30 | 39.99 – 1.60 | 39.97 – 1.20 |
| | (1.76 – 1.73)* | (1.61 – 1.58) | (1.58 – 1.55)* | (1.32 – 1.30) | (1.63 – 1.60)* | (1.22 – 1.20) |
| $R_{merge}$ (%) | 5.4 (58.9) | 5.1 (58.1) | 5.9 (64.9) | 10.9 (64.9) | 4.7 (55.9) | 4.0 (3.6) |
| $R_{meas}$ (%) | 6.4 (70.0) | 6.1 (70.4) | 8.0 (88.6) | 13.9 (84.3) | 6.0 (72.6) | 5.4 (49.5) |
| $CC_{1/2}$ (%) | 99.8 (84.5) | 99.9 (84.0) | 99.7 (55.8) | 98.5 (80.6) | 99.8 (74.8) | 99.9 (84.4) |
| $\langle I/\sigma(I) \rangle$ | 16.3 (2.6) | 17.2 (2.6) | 10.4 (1.7) | 6.8 (1.7) | 15.4 (2.3) | 14.5 (2.9) |
| No. of reflections | 147910 (8024) | 203376 (8825) | 122879 (6090) | 250702 (11497) | 119169 (5597) | 228162 (10764) |
| No. of unique reflections | 22182 (1195) | 30705 (1492) | 33023 (1610) | 56215 (2726) | 27495 (1321) | 63701 (2967) |
| Multiplicity | 6.7 (6.7) | 6.6 (5.9) | 3.7 (3.8) | 4.5 (4.2) | 4.3 (4.2) | 3.6 (3.6) |
| Completeness (%) | 99.4 (99.1) | 100.0 (99.9) | 99.7 (99.9) | 99.8 (98.8) | 97.7 (99.0) | 97.4 (92.7) |
| $R_{work}/R_{free}$§ (%) | 20.02 / 23.55 | 16.21 / 19.67 | 16.86 / 21.08 | 14.50 / 17.91 | 16.31 / 19.80 | 13.75 / 17.04 |
| On/Off -state chrom. occ. (%) | 100 / 0 | 42 / 58 | 100 / 0 | 0 / 100 | 100 / 0 | 0 / 100 |
| RMSD from ideal | | | | | | |
| Bond lengths (Å) | 0.004 | 0.007 | 0.006 | 0.007 | 0.006 | 0.01 |
| Bond angles (°) | 0.766 | 0.904 | 0.883 | 0.974 | 0.916 | 1.02 |
| No. of protein atoms | 1872 | 1991 | 1958 | 3937 | 1935 | 4156 |
| No. of water molecules | 136 | 295 | 271 | 322 | 219 | 360 |
| Average isotropic B-factors ($Å^2$) | | | | | | |
| Main chain | 36.00 | 19.86 | 20.05 | 18.72 | 19.22 | 12.11 |
| Side chain | 39.19 | 25.03 | 23.98 | 22.22 | 23.44 | 17.14 |
| Water molecules | 42.36 | 36.56 | 33.92 | 30.81 | 32.60 | 27.57 |
| Ligands | NA | NA | 36.31 | 62.02 | NA | NA |
| Ramachandranplot# (%) | | | | | | |
| Residues in favored regions | 97.30 | 98.65 | 98.69 | 99.12 | 98.65 | 98.65 |
| Outliers | 0.00 | 0.00 | 0.00 | 0.00 | 0.00 | 0.00 |
| Rotamer outlier# (%) | 1.00 | 0.94 | 0.94 | 1.35 | 0.48 | 1.31 |

* Values in parentheses are for the highest resolution shell.
§ $R_{free}$ is calculated using a random 5% of data excluded from the refinement.
#Ramachandran and rotamer analysis was carried out using Molprobity [19].

*(Continued)* Data collection, processing and refinement statistics.

| | rsGreen1 Green-on | rsGreen1-K206A Green-off | rsGreenF-K206A Green-on | rsGreenF-K206A Green-off | rsGreen0.7-K206A Green-on | rsGreen0.7-K206A Green-off |
|---|---|---|---|---|---|---|
| PDB ID | 7A7U | 7A7V | 7A7Q | 7A7R | 7A7O | 7A7P |
| Beamline | Soleil, Proxima2A | Soleil, Proxima2A | Soleil, Proxima2A | Soleil, Proxima2A | Elettra, XRD1 | Soleil, Proxima2A |
| Wavelength (Å) | 0.98 | 0.98 | 0.98 | 0.98 | 1.00 | 0.98 |
| Space group | $P4_32_12$ | $P4_32_12$ | $P4_12_12$ | $P4_12_12$ | $H3_2$ | $P22_12_1$ |
| Unit cell parameters | | | | | | |
| a, b, c (Å) | 73.24 73.24 114.73 | 73.55 73.55 114.38 | 106.31 106.31 35.73 | 106.18 106.18 35.91 | 140.21 140.21 72.89 | 50.81 61.35 65.98 |
| $\alpha$, $\beta$, $\gamma$ (°) | 90.00 90.00 90.00 | 90.00 90.00 90.00 | 90.00 90.00 90.00 | 90.00 90.00 90.00 | 90.00 90.00 120.00 | 90.00 90.00 90.00 |
| Resolution range (Å) | 47.20 – 2.15 | 47.34 – 2.00 | 47.54 – 2.00 | 47.49 – 2.35 | 46.65 - 1.80 | 44.93 – 1.48 |
| | (2.22 – 2.15)* | (2.05 – 2.00) | (2.05 – 2.00)* | (2.43 – 2.35) | (1.84 – 1.80)* | (1.51 – 1.48) |
| $R_{merge}$ (%) | 5.8 (57.3) | 5.2(60.6) | 8.2 (75.8) | 14.2 (65.3) | 4.8 (68.5) | 5.7 (63.8) |
| $R_{meas}$ (%) | 6.0 (59.5) | 5.4 (62.9) | 8.5 (78.6) | 15.4 (70.9) | 5.4 (78.6) | 6.7 (75.0) |
| $R_{p.i.m.}$ (%) | 1.6 (15.9) | 1.4 (16.6) | 2.3 (2.1) | 5.9 (27.4) | 2.6 (38.1) | 3.5 (38.7) |
| $CC_{1/2}$ (%) | 1.00 (98.4) | 100.0 (98.7) | 99.7 (94.2) | 99.7 (91.5) | 100.0 (88.8) | 99.8 (87.6) |
| $<I/\sigma(I)>$ | 36.6 (6.5) | 38.5 (6.1) | 26.6 (5.0) | 16.3 (4.7) | 24.6 (3.3) | 15.1 (2.2) |
| No. of reflections | 443705 (39597) | 565641 (43151) | 362669 (27584) | 112431 (11045) | 212503 (12345) | 240614 (12240) |
| No. of unique reflections | 39597 (1499) | 21948 (1585) | 14432 (1040) | 9057 (870) | 25487 (1509) | 35063 (1716) |
| Multiplicity | 25.1 (26.4) | 25.8 (27.2) | 25.1 (26.5) | 12.4 (12.4) | 8.3 (8.2) | 6.9 (7.1) |
| Completeness (%) | 100.0 (99.9) | 100.0 (100.0) | 100.0 (100.0) | 100.0 (100.0) | 100.0 (100.0) | 99.8 (100.0) |
| $R_{work}/R_{free}$§ (%) | 15.95 / 20.12 | 15.92 / 19.36 | 16.14 / 21.80 | 17.39 / 24.32 | 19.52 / 24.12 | 16.96 / 19.73 |
| On/Off -state chrom. occ. (%) | 100 / 0 | 33 / 67 | 100 / 0 | 0 / 100 | 100 / 0 | 67 / 33 |
| RMSD from ideal | | | | | | |
| Bond lengths (Å) | 0.007 | 0.007 | 0.006 | 0.001 | 0.008 | 0.006 |
| Bond angles (°) | 0.925 | 0.876 | 0.841 | 0.459 | 0.988 | 0.885 |
| No. of protein atoms | 1851 | 1856 | 1844 | 1825 | 1887 | 1880 |
| No. of water molecules | 174 | 237 | 143 | 136 | 227 | 282 |
| Average isotropic B-factors ($Å^2$) | | | | | | |
| Main chain | 42.82 | 37.78 | 31.65 | 23.20 | 28.14 | 20.13 |
| Side chain | 45.42 | 41.37 | 35.66 | 26.43 | 30.47 | 23.66 |
| Water molecules | 52.61 | 50.54 | 41.24 | 29.39 | 36.11 | 33.02 |
| Ligands | NA | 60.214 | 46.85 | 44.29 | NA | 28.39 |
| Ramachandranplot# (%) | | | | | | |
| Residues in favored regions | 99.09 | 99.1 | 99.08 | 98.62 | 97.81 | 99.1 |
| Outliers | 0.00 | 0.00 | 0.00 | 0.00 | 0.00 | 0.00 |
| Rotamer outlier# (%) | 0.00 | 0.51 | 1.01 | 1.03 | 0 | 0.00 |

* Values in parentheses are for the highest resolution shell.

§ $R_{free}$ is calculated using a random 5% of data excluded from the refinement.

#Ramachandran and rotamer analysis was carried out using Molprobity [19].

*(Continued)* Data collection, processing and refinement statistics.

| | rsGreen0.7-K206A-N205S Green-on | rsGreen0.7-K206A-N205G Green-on | rsGreen0.7-K206A-N205C Green-on | rsGreen0.7-K206A-N205L Green-on | rsGreen0.7-K206A-E222G Green-on | rsGreen0.7-K206A-E222V Green-on |
|---|---|---|---|---|---|---|
| PDB ID | 7A8O | 7A8M | 7A8L | 7A8N | 7A7Z | 7A80 |
| Beamline | Elettra, XRD1 | Elettra, XRD1 | SLS, X06DA | Soleil, Proxima2A | Elettra, XRD1 | Soleil, Proxima1 |
| Wavelength (Å) | 1.00 | 1.00 | 1.00 | 0.98 | 1.00 | 0.98 |
| Space group | $P4_32_12$ | $H3_2$ | $H3_2$ | $H3_2$ | $I2$ | $P2_1$ |
| Unit cell parameters | | | | | | |
| a, b, c (Å) | 102.64 102.64 50.30 | 140.87 140.87 73.22 | 140.20 140.20 73.12 | 140.73 140.73 73.25 | 48.24 61.56 93.10 | 69.82 61.80 112.12 |
| $\alpha, \beta, \gamma$ (°) | 90.00 90.00 90.00 | 90.00 90.00 120.00 | 90.00 90.00 120.00 | 90.00 90.00 120.00 | 90.00 97.41 90.00 | 90.00 91.01 90.00 |
| Resolution range (Å) | 45.90 – 1.60 (1.63 – 1.60)* | 46.87 – 1.60 (1.63 – 1.60) | 46.71 – 1.75 (1.78 – 1.75)* | 46.84 – 2.10 (2.16 – 2.10) | 44.91 – 1.10 (1.12 – 1.10)* | 46.27 – 2.05 (2.10 – 2.05) |
| $R_{merge}$ (%) | 8.7 (63.9) | 4.2 (64.9) | 4.1 (74.3) | 5.6 (69.7) | 2.4 (56.5) | 9.1 (63.8) |
| $R_{meas}$ (%) | 9.9 (72.9) | 4.7 (72.2) | 4.5 (81.4) | 5.9 (73.4) | 3.2 (75.1) | 10.9 (75.8) |
| $R_{p.i.m.}$ (%) | 4.7 (34.6) | 2.0 (31.6) | 1.9 (33.0) | 1.8 (22.8) | 2.1 (49.1) | 5.8 (40.5) |
| $CC_{1/2}$ (%) | 99.8 (87.4) | 100.0 (91.9) | 100.0 (89.3) | 100.0 (96.4) | 100.0 (87.9) | 99.8 (86.1) |
| $\langle I/\sigma(I) \rangle$ | 12.8 (2.6) | 26.5 (3.5) | 33.1 (3.6) | 33.1 (4.6) | 16.7 (1.8) | 13.2 (2.9) |
| No. of reflections | 281204 (14240) | 367014 (18615) | 312941 (17909) | 324900 (26566) | 390381 (18148) | 405037 (30415) |
| No. of unique reflections | 36042 (1764) | 36684 (1820) | 27802 (1533) | 16005 (1307) | 108890 (5350) | 60229 (4424) |
| Multiplicity | 7.8 (8.1) | 10.0 (10.2) | 11.3 (11.7) | 20.3 (20.3) | 3.6 (3.4) | 6.7 (6.9) |
| Completeness (%) | 100.0 (100.0) | 100.0 (100.0) | 100.0 (100.0) | 98.3 (97.5) | 99.6 (99.7) | 99.9 (100.0) |
| $R_{work}/R_{free}$ § (%) | 16.05 / 20.12 | 20.20 / 24.46 | 18.81 / 22.50 | 20.84 / 25.75 | 12.27 / 14.24 | 18.29 / 23.43 |
| On/Off -state chrom. occ. (%) | 100 / 0 | 100 / 0 | 100 / 0 | 100 / 0 | 100 / 0 | 100 / 0 100 / 0† 100 / 0 100 / 0 |
| RMSD from ideal | | | | | | |
| Bond lengths (Å) | 0.006 | 0.006 | 0.007 | 0.002 | 0.007 | 0.002 |
| Bond angles (°) | 0.862 | 0.86 | 0.847 | 0. 524 | 1.034 | 0.557 |
| No. of protein atoms | 1984 | 1943 | 1956 | 1874 | 4404 | 7465 |
| No. of water molecules | 382 | 288 | 239 | 190 | 506 | 582 |
| Average isotropic B-factors ($Å^2$) | | | | | | |
| Main chain | 18.63 | 28.08 | 30.174 | 43.631 | 14.73 | 24.86 / 27.48/† 28.79 / 26.26 |
| Side chain | 21.86 | 30.80 | 33.08 | 45.886 | 19.78 | 29.47 / 32.04/ 33.33 / 30.63 |
| Water molecules | 35.45 | 39.24 | 39.13 | 46.711 | 33.60 | 35.69 |
| Ligands | 42.93 | NA | NA | NA | NA | 37.64 |
| Ramachandranplot# (%) | | | | | | |
| Residues in favored regions | 98.25 | 98.25 | 97.82 | 99.12 | 99.09 | 98.31 |
| Outliers | 0.00 | 0.00 | 0.44 | 0.00 | 0.00 | 0.00 |
| Rotamer outlier# (%) | 0.00 | 0.00 | 0.00 | 1.01 | 0.42 | 0.00 |

* Values in parentheses are for the highest resolution shell.

§ $R_{free}$ is calculated using a random 5% of data excluded from the refinement.

\# Ramachandran and rotamer analysis was carried out using Molprobity [19].

† Values reported per protein chain.

*(Continued)* Data collection, processing and refinement statistics.

| | rsGreen0.7-K206A-F145H Green-on | rsGreen0.7-K206A-F145H Green-off | rsGreen0.7-K206A-F145Q Green-on | rsGreen0.7-K206A-F145Q Green-off | rsGreen0.7-K206A-F145L Green-on | rsGreen0.7-K206A-F145L Green-off |
|---|---|---|---|---|---|---|
| PDB ID | 7A83 | 7A84 | 7A89 | 7A8A | 7A85 | 7A86 |
| Beamline | Soleil, Proxima2A | Soleil, Proxima2A | Soleil, Proxima2A | Soleil, Proxima2A | Elettra, XRD1 | Soleil, Proxima2A |
| Wavelength (Å) | 0.98 | 0.98 | 0.98 | 0.98 | 1.00 | 0.98 |
| Space group | $I2$ | $H3_2$ | $H3_2$ | $H3_2$ | $I2$ | $P1$ |
| Unit cell parameters | | | | | | |
| a, b, c (Å) | 45.75 61.13 91.83 | 139.97 139.97 73.24 | 135.41 135.41 144.54 | 140.76 140.76 73.26 | 46.30 60.91 92.69 | 48.10 58.29 58.47 |
| $\alpha, \beta, \gamma$ (°) | 90.00 97.16 90.00 | 90.00 90.00 120.00 | 90.00 90.00 120.00 | 90.00 90.00 120.00 | 90.00 96.85 90.00 | 64.06 71.06 72.61 |
| Resolution range (Å) | 42.82 – 1.73 (1.76 – 1.73)* | 46.69 – 2.10 (2.16 – 2.10) | 48.18 – 2.50 (2.60 – 2.50)* | 46.86 – 1.97 (2.02 – 1.97) | 43.24 – 1.52 (1.55 – 1.52)* | 44.72 – 1.90 (1.94 – 1.90) |
| $R_{merge}$ (%) | 4.2 (64.2) | 4.5 (75.2) | 6.0 (69.2) | 5.0 (43.4) | 4.3 (52.9) | 3.2 (45.5) |
| $R_{meas}$ (%) | 4.9 (75.4) | 4.7 (78.9) | 6.4 (72.9) | 5.3 (45.8) | 5.8 (710) | 4.5 (64.3) |
| $R_{p.i.m.}$ (%) | 2.6 (39.3) | 1.4 (23.6) | 2.0 (22.9) | 1. 6 (14.5) | 38 (46.9) | 3.2 (45.5) |
| $CC_{1/2}$ (%) | 99.9 (84.8) | 100.0 (97.6) | 99.9 (97.5) | 100.0 (98.6) | 99.9 (80.3) | 99.9 (90.3) |
| $<I/\sigma(I)>$ | 21.0 (2.8) | 36.8 (5.0) | 31.2 (5.1) | 35.9 (7.7) | 12.4 (1.7) | 14.8 (2.5) |
| No. of reflections | 178548 (10045) | 323440 (28112) | 355351 (39370) | 397345 (28041) | 146055 (7044) | 140799 (9580) |
| No. of unique reflections | 25749 (1406) | 15971 (1294) | 17818 (1989) | 19753 (1410) | 39087 (1909) | 40412 (2624) |
| Multiplicity | 6.9 (7.1) | 20.3 (21.7) | 19.9 (19.8) | 20.1 (19.9) | 3.7 (3.7) | 3.5 (3.7) |
| Completeness (%) | 98.1 (97.0) | 99.1 (98.3)) | 100.0 (99.9) | 100.0 (99.9) | 99.2 (99.4) | 96.6 (96.2) |
| $R_{work}/R_{free}$ § (%) | 16.57 / 19.24 | 22.10 / 26.88 | 17.80 / 23.43 | 20.37 / 25.38 | 21.16 / 23.68 | 18.10 / 21.05 |
| On/Off -state chrom. occ. (%) | 100 / 0 | 39 / 61 | 100 / 0 100 / 0 | 30 / 70 | 100 / 0 | 59 / 41 46 / 54† |
| RMSD from ideal | | | | | | |
| Bond lengths (Å) | 0.007 | 0.008 | 0.007 | 0.007 | 0.006 | 0.007 |
| Bond angles (°) | 0.937 | 0.976 | 0.909 | 0.909 | 0.879 | 0.913 |
| No. of protein atoms | 1920 | 1846 | 3636 | 1955 | 1886 | 3826 |
| No. of water molecules | 192 | 82 | 65 | 179 | 268 | 256 |
| Average isotropic B-factors ($Å^2$) | | | | | | |
| Main chain | 29.58 | 53.76 | 63.33 / 68.67 | 36.03 | 23.13 | 45.43 / 42.00† |
| Side chain | 34.78 | 55.64 | 66.32 / 70.96 | 38.17 | 26.39 | 47.36 / 45.73 |
| Water molecules | 42.57 | 52.78 | 59.87 | 40.90 | 35.21 | 49.16 |
| Ligands | 51.92 | NA | NA | NA | 40.28 | 62.36 |
| Ramachandranplot# (%) | | | | | | |
| Residues in favored regions | 98.20 | 97.73 | 97.51 | 96.48 | 98.17 | 99.09 |
| Outliers | 0.00 | 0.00 | 0.23 | 0.00 | 0.00 | 0.00 |
| Rotamer outlier# (%) | 0.97 | 0.00 | 0.00 | 0.96 | 0.00 | 0.99 |

* Values in parentheses are for the highest resolution shell.

§ $R_{free}$ is calculated using a random 5% of data excluded from the refinement.

#Ramachandran and rotamer analysis was carried out using Molprobity [19].

† Values reported per protein chain.

*(Continued)* Data collection, processing and refinement statistics.

| | rsGreen0.7-K206A-F145M Green-on | rsGreen0.7-K206A-F145M Green-off | rsGreen0.7-K206A-F145A Green-on | rsGreen0.7-K206A-F145A Green-off | rsGreen0.7-K206A-F145S Green-on | rsGreen0.7-K206A-F145S Green-off |
|---|---|---|---|---|---|---|
| PDB ID | 7A87 | 7A88 | 7A81 | 7A82 | 7A8B | 7A8C |
| Beamline | SLS, X06DA | Soleil, Proxima1 | SLS, X06DA | Soleil, Proxima2A | DLS, I02 | Soleil, Proxima2A |
| Wavelength (Å) | 1.00 | 0.98 | 1.00 | 0.98 | 0.98 | 0.98 |
| Space group | $H3_2$ | $H3_2$ | $H3_2$ | $H3_2$ | $H3_2$ | $H3_2$ |
| Unit cell parameters | | | | | | |
| a, b, c (Å) | 139.31 139.31 72.46 | 139.98 139.98 72.64 | 140.89 140.89 72.65 | 141.36 141.36 73.11 | 139.55 139.55 72.86 | 140.62 140.62 72.91 |
| $\alpha, \beta, \gamma$ (°) | 90.00 90.00 120.00 | 90.00 90.00 120.00 | 90.00 90.00 120.00 | 90.00 90.00 120.00 | 90.00 90.00 120.00 | 90.00 90.00 120.00 |
| Resolution range (Å) | 46.36 – 1.75 (1.78 – 1.75)* | 46.54 – 1.95 (2.00 – 1.95) | 46.72 – 1.70 (1.73 – 1.70)* | 46.93 – 1.90 (1.94 – 1.90) | 46.51 – 1.55 (1.58 – 1.55)* | 46.73 – 2.13 (2.19 – 2.13) |
| $R_{merge}$ (%) | 5.0 (66.4) | 6. 4 (66.4) | 4.8 (63.0) | 4.5 (75.9) | 4.1 (64.5) | 10.7 (61.3) |
| $R_{meas}$ (%) | 5.5 (73.5) | 6.8 (70.7) | 5.4 (70.0) | 4.7 (79.8) | 4.6 (72.6) | 11.3 (64.5) |
| $R_{p.i.m.}$ (%) | 2.4 ( 31.2) | 2.3 (24.2) | 2.4 (30.4) | 1.4 (24.7) | 2.0 (32.9) | 3.5 (20.1) |
| $CC_{1/2}$ (%) | 100.0 (90.1) | 100.0 (93.8) | 100.0 (91.0) | 100.0 (96.7) | 99.9 (87.3) | 99.8 (97.3) |
| $<I/\sigma(I)>$ | 25.8 (3.8) | 26.3 (4.7) | 25.0 (4.0) | 37.7 (4.5) | 22.2 (2.7) | 19.5 (5.8) |
| No. of reflections | 276606 (15638) | 332432 (23052) | 287315 (15892) | 445440 (29009) | 384350 (17407) | 315941 (25612) |
| No. of unique reflections | 27188 (1478) | 19962 (1407) | 29944 (1570) | 22123 (1410) | 38704 (1885) | 15569 (1261) |
| Multiplicity | 10.2 (10.6) | 16.7 (16.4) | 9.6 (10.1) | 20.1 (20.6) | 9.9 (9.2) | 20.3 (20.3) |
| Completeness (%) | 100.0 (99.8) | 99.9 (99.7) | 98.7 (97.7) | 100.0 (100.0) | 98.5 (97.6) | 100.0 (100.0) |
| $R_{work}/R_{free}$§ (%) | 19.16 / 22.77 | 17.65 / 22.61 | 19.87 / 24.82 | 19.89 / 24.99 | 19.32 / 23.21 | 20.09 / 25.27 |
| On/Off -state chrom. occ. (%) | 100 / 0 | 0 / 100 | 33 / 67 | 23 / 77 | 70 / 30 | 0 / 100 |
| RMSD from ideal | | | | | | |
| Bond lengths (Å) | 0.005 | 0.007 | 0.007 | 0.007 | 0.006 | 0.009 |
| Bond angles (°) | 0.842 | 0.922 | 0.841 | 0.9 | 0.854 | 0.91 |
| No. of protein atoms | 1961 | 1925 | 1930 | 1892 | 2008 | 1818 |
| No. of water molecules | 222 | 226 | 253 | 174 | 255 | 169 |
| Average isotropic B-factors (Å$^2$) | | | | | | |
| Main chain | 29.90 | 37.78 | 30.48 | 37.97 | 34.04 | 39.87 |
| Side chain | 32.47 | 40.40 | 32.78 | 40.65 | 36.33 | 42.15 |
| Water molecules | 38.01 | 47.29 | 40.39 | 44.06 | 44.44 | 42.85 |
| Ligands | 39.47 | 61.41 | NA | NA | NA | 61.624 |
| Ramachandranplot# (%) | | | | | | |
| Residues in favored regions | 98.70 | 96.49 | 97.79 | 97.35 | 98.25 | 96.38 |
| Outliers | 0.00 | 0.00 | 0.44 | 0.44 | 0.00 | 0.00 |
| Rotamer outlier# (%) | 1.43 | 0.49 | 0.00 | 1.00 | 1.43 | 0.00 |

* Values in parentheses are for the highest resolution shell.
§ $R_{free}$ is calculated using a random 5% of data excluded from the refinement.
#Ramachandran and rotamer analysis was carried out using Molprobity [19].

*(Continued)* Data collection, processing and refinement statistics.

| | rsGreen0.7-F145M Green-on | rsGreen0.7-F145M Green-off | rsGreen0.7-F145Q Green-on | rsGreen0.7-K106A-F165L Green-on | rsGreen0.7-K206A-F165W Green-on | rsGreen0.7-K206A-F165W Green-off |
|---|---|---|---|---|---|---|
| PDB ID | 7A7W | 7A7X | 7A7Y | 7A8F | 7A8D | 7A8E |
| Beamline | Elettra, XRD1 | Soleil, Proxima2A | Elettra, XRD1 | Soleil, Proxima2A | SLS, X06DA | Soleil, Proxima1 |
| Wavelength (Å) | 1.00 | 0.98 | 1.00 | 0.98 | 1.00 | 0.98 |
| Space group | $P2_12_12_1$ | $P2_12_12_1$ | $P2_1$ | $H3_2$ | $H3_2$ | $H3_2$ |
| Unit cell parameters | | | | | | |
| a, b, c (Å) | 31.81 61.30 105.21 | 31.92 62.08 106.45 | 47.20 51.27 48.58 | 141.74 141.74 73.04 | 140.57 140.57 72.97 | 140.74 140.74 72.90 |
| $\alpha, \beta, \gamma$ (°) | 90.00 90.00 90.00 | 90.00 90.00 90.00 | 90.00 101.08 90.00 | 90.00 90.00 120.00 | 90.00 90.00 120.00 | 90.00 90.00 120.00 |
| Resolution range (Å) | 39.92 – 1.30 (1.32 – 1.30)* | 40.41 – 1.85 (1.89 – 1.85) | 47.68 – 0.97 (0.99 – 0.97) | 46.99 – 2.27 (2.34 – 2.27) | 46.74 – 1.65 (1.68 – 1.65)* | 46.76 – 2.20 (2.27 – 2.20) |
| $R_{merge}$ (%) | 7.5 (35.5) | 8.2 (62.3) | 5.4 (47.0) | 6.3 (59.5) | 4.2 (71.1) | 5.7 (64.1) |
| $R_{meas}$ (%) | 9.9(47.1) | 8.9 (67.8) | 7.5 (63.3) | 6.6 (62.3) | 4.7 (78.7) | 6.0 (67.7) |
| $R_{p.i.m.}$ (%) | 6.5 (30.6) | 3.4 (26.7) | 5.2 (41.9) | 2.0 (18.7) | 1.9 (33.4) | 1. 9 (21.6) |
| $CC_{1/2}$ (%) | 99.5 (83.0) | 99.9 (93.4) | 99.5 (74.5) | 100.0 (98.7) | 99.9 (89.4) | 100.0 (96.4) |
| $<I/\sigma(I)>$ | 9.1 (3.0) | 19.1 (4.3) | 10.9 (1.8) | 29.0 (5.6) | 27.7 (3.2) | 32.1 (5.3) |
| No. of reflections | 183154 (9519) | 235440 (14647) | 405332 (8645) | 268648 (25721) | 371129 (17525) | 284967 (23391) |
| No. of unique reflections | 51172 (2712) | 18843 (1160) | 129948 (4118) | 13100 (1187) | 33227 (1624) | 14148 (1211) |
| Multiplicity | 3.6 (3.5) | 12.5 (12.6) | 3.1 (2.1) | 20.5 (21.7) | 11.2 (10.8) | 20.1 (19.3) |
| Completeness (%) | 99.2 (99.4) | 100.0 (100.0) | 96.8 (62.3) | 99.9 (99.7) | 100.0 (100.0) | 100.0 (99.9) |
| $R_{work}/R_{free}$ § (%) | 12.78 / 16.45 | 14.73 / 20.70 | 12.53 / 14.17 | 19.64 / 25.71 | 19.19 / 21.91 | 20.50 / 26.13 |
| On/Off -state chrom. occ. (%) | 71 / 29 | 0 / 100 | 100 / 0 | 100 / 0 | 65 / 35 | 53 / 47 |
| RMSD from ideal | | | | | | |
| Bond lengths (Å) | 0.008 | 0.006 | 0.007 | 0.005 | 0.006 | 0.002 |
| Bond angles (°) | 1.079 | 0.877 | 1.052 | 0.8 | 0.905 | 0.542 |
| No. of protein atoms | 4072 | 1901 | 4526 | 1843 | 1988 | 1886 |
| No. of water molecules | 398 | 269 | 506 | 105 | 272 | 123 |
| Average isotropic B-factors ($Å^2$) | | | | | | |
| Main chain | 9.82 | 18.65 | 10.18 | 50.39 | 31.96 | 49.45 |
| Side chain | 13.94 | 23.57 | 14.36 | 52.34 | 34.71 | 51.56 |
| Water molecules | 26.58 | 33.49 | 27.58 | 50.06 | 43.35 | 50.35 |
| Ligands | NA | NA | 35.82 | NA | NA | NA |
| Ramachandranplot# (%) | | | | | | |
| Residues in favored regions | 98.68 | 98.65 | 98.29 | 96.02 | 98.68 | 98.21 |
| Outliers | 0.00 | 0.00 | 0.00 | 0.00 | 0.00 | 0.00 |
| Rotamer outlier# (%) | 1.35 | 0 | 0.40 | 0.51 | 0.48 | 0.50 |

* Values in parentheses are for the highest resolution shell.
§ $R_{free}$ is calculated using a random 5% of data excluded from the refinement.
#Ramachandran and rotamer analysis was carried out using Molprobity [19].

*(Continued)* Data collection, processing and refinement statistics.

| | rsGreen0.7-K206A-H148V Green-on | rsGreen0.7-K206A-H148V Green-off | rsGreen0.7-K206A-H148S Green-on | rsGreen0.7-K206A-H148S Green-off | rsGreen0.7-K206S-H148G Green-on |
|---|---|---|---|---|---|
| PDB ID | 7A8J | 7A8K | 7A8H | 7A8I | 7A8G |
| Beamline | Elettra, XRD1 | Elettra, XRD1 | Soleil, Proxima2A | Soleil, Proxima2A | Soleil, Proxima2A |
| Wavelength (Å) | 1.00 | 1.00 | 0.98 | 0.98 | 0.98 |
| Space group | $H3_2$ | $I2$ | $P2_12_12_1$ | $P2_12_12_1$ | $P22_12_1$ |
| Unit cell parameters | | | | | |
| a, b, c (Å) | 140.62 140.62 73.00 | 48.01 68.93 76.38 | 97.59 98.60 100.04 | 96.29 97.88 98.34 | 49.44 88.62 131.36 |
| $\alpha, \beta, \gamma$ (°) | 90.00 90.00 120.00 | 90.00 94.84 90.00 | 90.00 90.00 90.00 | 90.00 90.00 90.00 | 90.00 90.00 90.00 |
| Resolution range (Å) | 46.76 – 1.85 (1.89 – 1.85)* | 42.13 – 2.25 (2.32 – 2.25) | 44.51 – 1.70 (1.73 – 1.70)* | 48.94 – 2.00 (2.05 – 2.00) | 49.44 – 2.00 (2.05 – 2.00)* |
| $R_{merge}$ (%) | 4.8 (62.9) | 6.7 (60.6) | 4.8 (73.3) | 7.8 (70.1) | 5.2 (58.8) |
| $R_{meas}$ (%) | 5.4 (70.0) | 8.9 (82.1) | 5.2 (79.5) | 8.4 (75.8) | 6.0 (68.2) |
| $R_{p.i.m.}$ (%) | 2.3 (30.4) | 5.9 (55.0) | 2.0 (30.4) | 3.2 (28.7) | 3.0 (34.0) |
| $CC_{1/2}$ (%) | 100.0 (93.2) | 99.7 (84.5) | 100.0 (90.9) | 99.9 (90.4) | 100.0 (96.5) |
| $<I/\sigma(I)>$ | 28.4 (3.8) | 10.0 (1.9) | 27.9 (3.6) | 18.1 (3.6) | 19.9 (3.5) |
| No. of reflections | 241673 (14811) | 43972 (3804) | 1384862 (70818) | 816370 (60535) | 297790 (22425) |
| No. of unique reflections | 23673 (1443) | 11804 (1104) | 106573 (5259) | 63498 (4430) | 39875 (2908) |
| Multiplicity | 10.2 (10.3) | 3.7 (3.4) | 13.0 (13.5) | 12.9 (13.7) | 7.5 (7.7) |
| Completeness (%) | 100.0 (100.0 ) | 99.4 (99.5) | 100.0 (99.9) | 100.0 (100.0) | 99.9 (99.9) |
| $R_{work}/R_{free}$§ (%) | 18.90 / 23.15 | 18.88 / 26.46 | 17.92 / 20.91 | 16.43 / 22.84 | 17.94 / 21.71 |
| On/Off -state chrom. occ. (%) | 100 / 0 | 50 / 50 | 100 / 0 100 / 0† 100 / 0 100 / 0 | 62 / 38 100 /0 52 / 48 100 / 0 | 100 / 0 100 / 0† |
| RMSD from ideal | | | | | |
| Bond lengths (Å) | 0.009 | 0.008 | 0.007 | 0.007 | 0.007 |
| Bond angles (°) | 0.956 | 1.002 | 0.868 | 0.921 | 0.915 |
| No. of protein atoms | 1894 | 1846 | 7583 | 7483 | 3667 |
| No. of water molecules | 232 | 66 | 893 | 616 | 255 |
| Average isotropic B-factors ($Å^2$) | | | | | |
| Main chain | 32.51 | 49.40 | 23.89 / 29.52 /† 25.73 / 26.22 | 29.73 / 33.77 / 33.42 / 35.34 | 32.90 / 32.83† |
| Side chain | 34.81 | 51.01 | 28.17 / 33.17 / 29.32 / 31.04 | 34.10 / 37.26 / 37.61 / 39.43 | 37.21 / 36.92 |
| Water molecules | 40.66 | 49.93 | 37.98 | 41.75 | 41.08 |
| Ligands | 51.43 | 56.079 | 36.45 | NA | 57.75 |
| Ramachandranplot# (%) | | | | | |
| Residues in favored regions | 96.02 | 95.91 | 98.54 | 98.53 | 98.4 |
| Outliers | 0.00 | 0.45 | 0.00 | 0.00 | 0.00 |
| Rotamer outlier# (%) | 0.00 | 0.00 | 0.49 | 1.25 | 0.00 |

* Values in parentheses are for the highest resolution shell.
§ $R_{free}$ is calculated using a random 5% of data excluded from the refinement.
#Ramachandran and rotamer analysis was carried out using Molprobity [19].
† Values reported per protein chain.

**Table S3:** Structural quantification and PCA variable grouping names. See Figure S9 for chromophore-only involved descriptors and chromophore anchor points for distances and angles.

| | | | |
|---|---|---|---|
| **(i) Chromophore-only involved descriptors** = "Internal"[#] | | | |
| Tilt | Twist | Methylene bond angle | |
| **(ii) Distances chromophore – residues** = "<chromophore part or residue name>" | | | |
| Hydroxybenz$_{OH}$ | Hydroxybenz$_{planecenter}$ | Imidazolinone$_{O}$ | Imidazolinone$_{N}$ |
| A150$_{CB}$ | E222$_{CD}$ | L69$_{CD2}$ | E222$_{OE1/OE2}$* |
| F145$_{CA}$ | F165$_{planecenter}$ | R96$_{NH1}$ | |
| F145$_{CB}$ | H148[§]$_{planecenter}$ | | |
| F165$_{CZ}$ | R96$_{NE2}$ | | |
| H148$_{CA}$ | T62$_{CG2}$ | | |
| H148[§]$_{planecenter}$ | L69$_{CD2}$ | | |
| I167$_{CD1}$ | T203$_{CB}$ | | |
| L201$_{CD1}$ | | | |
| N205$_{CA}$ | | | |
| N205$_{OD1/ND2}$* | | | |
| N205$_{OD1/ND2/HOH}$* | | | |
| $H_2O$* | | | |
| **(iii) Distances residue – residue** = "<residue name>" | | | |
| Involved atoms: | | | |
| L69$_{CD2}$ | Q94$_{NE2}$ | R96$_{NE2}$ | F145$_{CA}$ |
| H148$_{CA}$ | A150$_{CB}$ | F165$_{CB}$ | I167$_{CD1}$ |
| L201$_{CD1}$ | T203$_{CB}$ | N205$_{CA}$ | E222$_{CD}$ |
| **(iv) Angles chromophore – residues** = "<chromophore part or residue name>" | | | |
| F165$_{plane}$ – Hydroxybenz$_{plane}$ | | F145$_{plane}$ – Hydroxybenz$_{plane}$ | |
| **(v) Chromophore environment** = "Pocket" | | | |
| Chromophore pocket volume | | Total No. of heteroatoms around chromophore[†] | |
| Total No. of atoms around Hydroxybenz$_{OH}$[†] | | Total No. of atoms around Hydroxybenz$_{plane}$[†] | |
| Total No. of atoms around Imidazolinone$_{plane}$[†] | | Total No. water molecules around chromophore[†] | |
| Total No. of H-bonds | | Total No. of H-bonds Hydroxybenz$_{OH}$ | |
| Total No. of H-bonds Imidazolinone$_{O}$ | | Total No. of H-bonds Imidazolinone$_{N}$ | |

[#] Variable groups used for visual inspection of PCA are between quotation marks.

* Atom that is closest to chromophore anchor point.

[§] If multiple alternative conformations: plane most close to chromophore anchor point

[†] Total No. of atoms within a sphere of 3.5 Å.

**Table S4:** Summary of Principal component analyses. All PCs showed in the table were used during the analysis.

**(a)** PCA with spectroscopic properties only.

| PC | Eigenvalue of cov(X) | Variance (%) | Cumulative variance(%) |
|---|---|---|---|
| 1 | 6.69 | 35.21 | 35.21 |
| 2 | 4.08 | 21.45 | 56.67 |
| 3 | 2.25 | 11.87 | 68.53 |
| 4 | 1.79 | 9.45 | 77.98 |
| 5 | 1.26 | 6.65 | 84.63 |
| 6 | 0.91 | 4.78 | 89.40 |
| 7 | 0.58 | 3.05 | 92.45 |
| 8 | 0.47 | 2.47 | 94.92 |
| 9 | 0.39 | 2.07 | 96.99 |
| 10 | 0.23 | 1.21 | 98.20 |

**(b)** PCA with spectroscopic properties and on-state structural descriptors.

| PC | Eigenvalue of cov(X) | Variance (%) | Cumulative variance(%) |
|---|---|---|---|
| 1 | 21.00 | 18.46 | 18.46 |
| 2 | 16.80 | 14.74 | 33.20 |
| 3 | 12.30 | 10.77 | 43.98 |
| 4 | 9.99 | 8.77 | 52.74 |
| 5 | 9.19 | 8.06 | 60.81 |
| 6 | 7.54 | 6.61 | 67.42 |
| 7 | 5.40 | 4.74 | 72.16 |
| 8 | 4.83 | 4.24 | 76.40 |
| 9 | 4.33 | 3.79 | 80.19 |
| 10 | 3.89 | 3.41 | 83.61 |
| 11 | 3.04 | 2.67 | 86.27 |
| 12 | 2.80 | 2.45 | 88.72 |
| 13 | 2.22 | 1.95 | 90.67 |

**Table S5:** 15 highest loadings (positive or negative) for relevant PCs in the PCA concerning spectroscopic and on-state structural descriptors in decreasing order.

| PC1 | PC2 |
|---|---|
| Hydroxybenz$_{OH}$ – A150$_{CB}$ | E222$_{CD}$ – Q94$_{NE2}$ |
| Hydroxybenz$_{OH}$ – L201$_{CD1}$ | A150$_{CB}$ – L69$_{CD2}$ |
| Hydroxybenz$_{OH}$ – F165$_{CZ}$ | F145$_{CA}$ – I167$_{CD1}$ |
| Hydroxybenz$_{planecenter}$ – L69$_{CD2}$ | F145$_{CA}$ – Q94$_{NE2}$ |
| Hydroxybenz$_{OH}$ – N205$_{CA}$ | A150$_{CB}$ – R96$_{NH2}$ |
| Hydroxybenz$_{planecenter}$ – F165$_{planecenter}$ | A150$_{CB}$ – L201$_{CD1}$ |
| Hydroxybenz$_{planecenter}$ – R96$_{NH2}$ | tilt |
| H148$_{CA}$ – Q94$_{NE2}$ | Hydroxybenz$_{OH}$ – H148$_{CA}$ |
| I167$_{CD1}$ – Q94$_{NE2}$ | N205$_{CA}$ – Q94$_{NE2}$ |
| Hydroxybenz$_{OH}$ – F145$_{CB}$ | A150$_{CB}$ –Q94$_{NE2}$ |
| Hydroxybenz$_{OH}$ – N205$_{OD1/ND2}$ | E222$_{CD}$ – L69$_{CD2}$ |
| H148$_{CA}$ – R96$_{NH2}$ | F145$_{CA}$ – R96$_{NH2}$ |
| Hydroxybenz$_{OH}$ – F145$_{CA}$ | Q$_{on}$ |
| Imidazolinone$_{O}$ – L69$_{CD2}$ | F145$_{CA}$ – L201$_{CD1}$ |
| I167$_{CD1}$ – R96$_{NH2}$ | F165$_{CB}$ – L69$_{CD2}$ |

| PC4 | PC5 | PC9 |
|---|---|---|
| Imidazolinone$_{N}$ – E222$_{OE1}$ | E222$_{CD}$ – F165$_{CB}$ | k$_{therm}$ |
| Hydroxybenz$_{OH}$ – N205$_{OD1/ND2/HOH}$ | E222$_{CD}$ – A150$_{CB}$ | Total No. of H-bonds Imidazolinone$_{O}$ |
| N205$_{CA}$ – T203$_{CB}$ | I167$_{CD1}$ – F165$_{CB}$ | Total No. of atoms around Imidazolinone$_{plane}$* |
| F165$_{CB}$ – L201$_{CD1}$ | E222$_{CD}$ – L201$_{CD1}$ | Total No. of heteroatoms around chromophore* |
| F145$_{CA}$ – L69$_{CD2}$ | Total No. of H-bonds Hydroxybenz$_{OH}$ | Total No. of waters around chromophore* |
| A150$_{CB}$– F165$_{CB}$ | F145$_{CA}$ – N205$_{CA}$ | I167$_{CD1}$ – H148$_{CA}$ |
| N205$_{CA}$ – H148$_{CA}$ | I167$_{CD1}$ – A150$_{CB}$ | Q94$_{NE2}$ – R96$_{NH2}$ |
| A$_{res,\infty}$ | E222$_{CD}$ – R96$_{NH2}$ | $\lambda_{abs}^{off}$ |
| A$_{res}$ | $\lambda_{ex}$-$\lambda_{abs}$ | I167$_{CD1}$ – F165$_{CB}$ |
| F145$_{CA}$ –E222$_{CD}$ | $\varepsilon_{anionic}$ | E222$_{CD}$ – T203$_{CB}$ |
| T203$_{CB}$ – R96$_{NH2}$ | T203$_{CB}$ – H148$_{CA}$ | F165$_{CB}$ – Q94$_{NE2}$ |
| T203$_{CB}$ – Q94$_{NE2}$ | E222$_{CD}$ – L69$_{CD2}$ | N205$_{CA}$ – L201$_{CD1}$ |
| T203$_{CB}$ – L69$_{CD2}$ | N205$_{CA}$ – E222$_{CD}$ | Total No. of heteroatoms around Hydroxybenz$_{OH}$ |
| F165$_{CB}$ – H148$_{CA}$ | Total No. of H-bonds | Twist |
| Hydroxybenz$_{planecenter}$ – T203$_{CB}$ | Q$_{fluo}$ | I167$_{CD1}$ – R96$_{NH2}$ |

*Total No. of atoms within a sphere of 3.5 Å.

**Table S6:** Pearson correlation coefficients between spectroscopic and on-state structural descriptors.

| | methylene bond angle | tilt | twist | Pocket Volume | Total No. H-bonds | Total No. H-bonds $Hydroxybenz_{OH}$ | Total No. H-bonds $Imidazolinone_N$ | Total No. H-bonds $Imidazolinone_O$ | Total No. of heteroatoms $Hydroxybenz_{OH}$ | Total No. of atoms $Hydroxybenz_{plane}$ | Total No. of atoms $Imidazolinone_{plane}$ | Total No. water molecules | Total No. of heteroatoms | $Hydroxybenz_{OH}$ – $A150_{CB}$ | $Hydroxybenz_{OH}$ – $F145_{CA}$ | $Hydroxybenz_{OH}$ – $F145_{CB}$ | $Hydroxybenz_{OH}$ – $F165_{CZ}$ | $Hydroxybenz_{OH}$ – $H148_{CA}$ | $Hydroxybenz_{OH}$ – $I167_{CD1}$ |
|---|---|---|---|---|---|---|---|---|---|---|---|---|---|---|---|---|---|---|---|
| $\lambda_{abs}$ | 0.21 | -0.32 | 0.08 | -0.17 | -0.34 | -0.44 | 0.16 | -0.07 | -0.31 | 0.10 | -0.11 | 0.01 | -0.23 | -0.32 | 0.38 | 0.37 | -0.25 | -0.32 | 0.08 |
| $\lambda_{ex}$ | 0.13 | -0.13 | 0.11 | -0.18 | 0.02 | -0.04 | 0.09 | 0.06 | 0.01 | 0.03 | -0.06 | 0.03 | -0.08 | -0.07 | 0.04 | 0.03 | -0.04 | -0.03 | 0.05 |
| $\lambda_{em}$ | -0.04 | -0.45 | 0.35 | -0.26 | 0.10 | 0.02 | 0.06 | 0.23 | -0.10 | 0.09 | -0.07 | 0.00 | -0.02 | 0.08 | -0.08 | -0.09 | 0.20 | -0.20 | 0.39 |
| $\lambda^{off}_{abs}$ | -0.07 | -0.46 | 0.03 | -0.04 | -0.05 | -0.23 | 0.33 | -0.07 | -0.42 | 0.47 | -0.08 | 0.11 | -0.17 | -0.41 | 0.48 | 0.49 | -0.27 | -0.42 | -0.06 |
| $A_{res}$ | -0.02 | -0.42 | 0.15 | 0.05 | -0.07 | -0.37 | 0.54 | -0.11 | -0.48 | 0.23 | -0.05 | 0.13 | -0.24 | -0.39 | 0.38 | 0.37 | -0.26 | -0.45 | 0.00 |
| $A_{res,\infty}$ | -0.07 | -0.35 | 0.15 | 0.12 | -0.07 | -0.42 | 0.63 | -0.10 | -0.46 | 0.20 | -0.01 | 0.16 | -0.19 | -0.39 | 0.38 | 0.36 | -0.26 | -0.46 | -0.07 |
| $\varepsilon_{off}$ | 0.04 | -0.15 | -0.13 | -0.08 | -0.32 | -0.36 | 0.09 | -0.19 | -0.23 | -0.07 | 0.00 | 0.15 | -0.18 | -0.35 | 0.32 | 0.29 | -0.33 | -0.17 | 0.02 |
| $pKa_{app}$ | 0.12 | -0.24 | 0.01 | -0.10 | -0.23 | -0.29 | 0.16 | -0.25 | -0.27 | -0.09 | 0.00 | 0.14 | -0.15 | -0.19 | 0.24 | 0.20 | -0.13 | -0.24 | 0.11 |
| $A_{unrecov}$ | 0.02 | -0.01 | -0.05 | 0.00 | -0.22 | -0.16 | -0.06 | -0.19 | -0.13 | -0.28 | -0.08 | 0.12 | -0.10 | 0.07 | -0.08 | -0.13 | 0.06 | 0.03 | 0.15 |
| $k_{therm}$ | 0.09 | -0.37 | -0.10 | -0.31 | 0.03 | 0.11 | -0.11 | -0.09 | -0.31 | 0.21 | -0.20 | -0.13 | -0.38 | -0.15 | 0.25 | 0.24 | -0.08 | -0.05 | 0.14 |
| $\varepsilon_{anionic}$ | -0.09 | -0.01 | 0.02 | 0.16 | -0.26 | -0.48 | 0.40 | -0.15 | -0.27 | 0.07 | -0.07 | -0.07 | -0.29 | -0.35 | 0.50 | 0.46 | -0.34 | -0.35 | -0.25 |
| $Q_{fluo}$ | -0.21 | 0.12 | -0.11 | -0.07 | 0.07 | 0.03 | 0.09 | -0.05 | 0.10 | 0.17 | -0.15 | 0.12 | 0.17 | 0.20 | -0.12 | -0.09 | 0.07 | -0.04 | 0.02 |
| M. br | -0.18 | 0.12 | -0.07 | 0.04 | 0.07 | 0.02 | 0.09 | 0.01 | 0.09 | 0.23 | -0.10 | -0.02 | 0.07 | 0.03 | 0.00 | 0.05 | -0.05 | -0.06 | -0.12 |
| $\varepsilon_{overall}$ | -0.03 | 0.12 | 0.00 | 0.15 | 0.04 | -0.03 | 0.08 | 0.11 | 0.03 | 0.16 | 0.06 | -0.13 | -0.04 | -0.11 | 0.10 | 0.14 | -0.11 | -0.07 | -0.27 |
| $Q_{off}$ | -0.19 | 0.48 | -0.11 | 0.27 | 0.36 | 0.58 | -0.37 | 0.09 | 0.50 | -0.11 | -0.08 | 0.13 | 0.31 | 0.49 | -0.53 | -0.54 | 0.37 | 0.70 | -0.18 |
| $\lambda_{ex} - \lambda_{abs}$ | -0.20 | 0.38 | 0.01 | 0.05 | 0.62 | 0.72 | -0.17 | 0.21 | 0.54 | -0.14 | 0.11 | 0.03 | 0.30 | 0.46 | -0.62 | -0.60 | 0.39 | 0.52 | -0.07 |
| $\delta_{Stokes}$ | -0.20 | -0.13 | 0.09 | 0.08 | 0.04 | 0.07 | -0.08 | 0.07 | -0.08 | 0.02 | 0.04 | -0.05 | 0.09 | 0.15 | -0.11 | -0.10 | 0.20 | -0.10 | 0.20 |
| $\Delta pH$ | 0.07 | -0.20 | -0.16 | -0.25 | -0.16 | -0.20 | 0.10 | -0.12 | -0.29 | -0.03 | 0.25 | -0.21 | -0.30 | -0.57 | 0.55 | 0.55 | -0.48 | -0.37 | -0.14 |
| $Q_{on}$ | -0.33 | -0.37 | 0.45 | 0.08 | 0.39 | 0.39 | 0.05 | 0.01 | 0.08 | 0.25 | -0.33 | 0.01 | 0.15 | 0.36 | 0.04 | 0.02 | 0.48 | -0.22 | 0.37 |

M. br: Molecular brightness; $\Delta pH$: $\lambda_{abs,pH10} - \lambda_{abs,pH6.5}$.

*(Continued)* Pearson correlation coefficients between spectroscopic and on-state structural descriptors.

| | Hydroxybenz$_{OH}$ – L201$_{CD1}$ | Hydroxybenz$_{OH}$ – N205$_{CA}$ | Hydroxybenz$_{OH}$ – N205$_{OD1/ND2}$* | Hydroxybenz$_{OH}$ – N205$_{OD1/ND2/HOH}$* | Hydroxybenz$_{planecenter}$ – E222$_{CD}$ | Hydroxybenz$_{planecenter}$ – F165$_{plane}$ | Hydroxybenz$_{planecenter}$ – H148$_{plneClose}$ | Hydroxybenz$_{planecenter}$ – R96$_{NH2}$ | Hydroxybenz$_{planecenter}$ – T62$_{CG2}$ | Hydroxybenz$_{planecenter}$ – L69$_{CD2}$ | Hydroxybenz$_{planecenter}$ – T203$_{CB}$ | Imidazoline$_{O}$ – L69$_{CD2}$ | Imidazoline$_{O}$ – R96$_{NH1}$ | Imidazoline$_{N}$ – E222$_{OE1/OE2}$* | Hydroxybenz$_{plane}$ – $_{F165}$ | A150$_{CB}$ – F165$_{CB}$ | A150$_{CB}$ – H148$_{CA}$ | A150$_{CB}$ – L201$_{CD1}$ | A150$_{CB}$ – Q94$_{NE2}$ |
|---|---|---|---|---|---|---|---|---|---|---|---|---|---|---|---|---|---|---|---|
| $\lambda_{abs}$ | -0.32 | 0.19 | 0.27 | -0.24 | 0.23 | -0.38 | -0.38 | -0.23 | 0.66 | -0.36 | 0.00 | 0.28 | -0.07 | 0.28 | 0.42 | 0.17 | -0.06 | -0.30 | -0.07 |
| $\lambda_{ex}$ | 0.00 | -0.02 | -0.01 | -0.27 | -0.04 | -0.16 | -0.38 | -0.07 | 0.41 | -0.07 | -0.04 | 0.18 | -0.14 | 0.19 | 0.29 | 0.03 | -0.18 | -0.18 | -0.11 |
| $\lambda_{em}$ | 0.06 | -0.18 | -0.08 | -0.31 | -0.05 | 0.04 | -0.27 | 0.11 | 0.31 | 0.03 | -0.13 | -0.09 | -0.55 | 0.08 | 0.42 | 0.13 | -0.22 | -0.20 | -0.27 |
| $\lambda^{off}_{abs}$ | -0.52 | 0.33 | 0.50 | 0.05 | 0.30 | -0.37 | -0.04 | -0.41 | 0.27 | -0.47 | 0.15 | 0.12 | -0.18 | 0.03 | 0.48 | 0.30 | 0.14 | -0.31 | -0.22 |
| $A_{res}$ | -0.40 | 0.27 | 0.40 | -0.20 | 0.24 | -0.40 | -0.06 | -0.35 | 0.49 | -0.44 | 0.00 | 0.02 | -0.06 | -0.07 | 0.48 | 0.41 | -0.07 | -0.36 | -0.29 |
| $A_{res,\infty}$ | -0.39 | 0.31 | 0.40 | -0.18 | 0.30 | -0.40 | -0.04 | -0.38 | 0.41 | -0.40 | -0.02 | -0.04 | -0.02 | -0.08 | 0.43 | 0.42 | -0.01 | -0.36 | -0.30 |
| $\varepsilon_{off}$ | -0.33 | 0.32 | 0.30 | 0.01 | 0.24 | -0.39 | -0.34 | -0.34 | 0.48 | -0.33 | 0.23 | 0.37 | 0.05 | 0.27 | 0.41 | 0.15 | 0.04 | -0.22 | 0.08 |
| $pKa_{app}$ | -0.15 | 0.09 | 0.15 | -0.07 | 0.03 | -0.25 | -0.33 | -0.13 | 0.44 | -0.17 | 0.13 | 0.25 | 0.00 | 0.37 | 0.31 | 0.20 | 0.04 | -0.31 | -0.06 |
| $A_{unrecov}$ | 0.14 | -0.11 | -0.11 | -0.10 | -0.14 | 0.02 | -0.30 | 0.08 | 0.14 | 0.12 | 0.02 | 0.18 | 0.03 | 0.30 | 0.09 | -0.03 | 0.00 | -0.09 | 0.12 |
| $k_{therm}$ | -0.23 | -0.01 | 0.16 | -0.03 | -0.12 | -0.21 | -0.21 | -0.12 | 0.38 | -0.29 | 0.23 | 0.33 | -0.20 | 0.13 | 0.49 | 0.23 | 0.10 | -0.41 | -0.21 |
| $\varepsilon_{anionic}$ | -0.37 | 0.32 | 0.30 | -0.23 | 0.40 | -0.45 | -0.26 | -0.39 | 0.44 | -0.39 | -0.09 | 0.09 | 0.29 | 0.06 | 0.27 | 0.20 | 0.23 | -0.40 | -0.14 |
| $Q_{fluo}$ | 0.08 | -0.03 | 0.17 | -0.33 | 0.13 | 0.16 | -0.13 | -0.05 | 0.01 | 0.15 | -0.33 | 0.01 | 0.07 | -0.02 | -0.04 | 0.13 | 0.27 | -0.21 | 0.06 |
| M. br | -0.07 | 0.10 | 0.21 | -0.25 | 0.24 | 0.03 | -0.01 | -0.10 | -0.03 | 0.00 | -0.30 | -0.03 | 0.14 | -0.26 | 0.00 | 0.12 | 0.22 | -0.13 | -0.04 |
| $\varepsilon_{overall}$ | -0.13 | 0.16 | 0.12 | -0.05 | 0.22 | -0.09 | 0.16 | -0.07 | -0.12 | -0.08 | -0.14 | -0.12 | 0.14 | -0.34 | -0.04 | 0.05 | 0.12 | -0.01 | -0.14 |
| $Q_{off}$ | 0.55 | -0.44 | -0.58 | 0.22 | -0.60 | 0.52 | 0.15 | 0.31 | -0.79 | 0.54 | 0.03 | -0.11 | 0.10 | -0.02 | -0.60 | -0.54 | -0.06 | 0.46 | 0.33 |
| $\lambda_{ex} - \lambda_{abs}$ | 0.56 | -0.36 | -0.48 | 0.06 | -0.46 | 0.45 | 0.15 | 0.31 | -0.60 | 0.53 | -0.05 | -0.24 | -0.06 | -0.23 | -0.35 | -0.27 | -0.13 | 0.27 | -0.02 |
| $\delta_{Stokes}$ | 0.03 | -0.09 | -0.04 | 0.16 | 0.03 | 0.24 | 0.35 | 0.18 | -0.36 | 0.12 | -0.03 | -0.32 | -0.18 | -0.22 | -0.11 | 0.05 | 0.10 | 0.12 | -0.04 |
| $\Delta pH$ | -0.54 | 0.55 | 0.46 | 0.47 | 0.47 | -0.61 | -0.22 | -0.46 | 0.24 | -0.51 | 0.57 | 0.31 | -0.10 | 0.37 | 0.47 | 0.19 | 0.15 | -0.16 | -0.21 |
| $Q_{on}$ | 0.18 | -0.41 | -0.22 | -0.15 | -0.21 | 0.29 | -0.16 | 0.31 | 0.12 | 0.16 | -0.15 | -0.22 | -0.26 | 0.26 | 0.06 | -0.10 | 0.21 | -0.27 | -0.24 |

M. br: Molecular brightness; $\Delta pH$: $\lambda_{abs,pH10} - \lambda_{abs,pH6.5}$.

*Atom that is closest to chromophore anchor point.

*(Continued)* Pearson correlation coefficients between spectroscopic and on-state structural descriptors.

| | $A150_{CB} - R96_{NH2}$ | $A150_{CB} - L69_{CD2}$ | $F165_{CB} - H148_{CA}$ | $F165_{CB} - L201_{CD1}$ | $F165_{CB} - Q94_{NE2}$ | $F165_{CB} - R96_{NH2}$ | $F165_{CB} - L69_{CD2}$ | $H148_{CA} - Q94_{NE2}$ | $H148_{CA} - R96_{NH2}$ | $H148_{CA} - L69_{CD2}$ |
|---|---|---|---|---|---|---|---|---|---|---|
| $\lambda_{abs}$ | -0.20 | -0.33 | -0.23 | -0.16 | 0.13 | 0.02 | -0.06 | 0.23 | 0.04 | -0.04 |
| $\lambda_{ex}$ | -0.27 | -0.28 | -0.39 | -0.16 | 0.06 | -0.10 | -0.13 | 0.11 | -0.09 | -0.11 |
| $\lambda_{em}$ | -0.50 | -0.42 | -0.51 | -0.02 | -0.02 | -0.27 | -0.24 | -0.19 | -0.42 | -0.44 |
| $\lambda_{abs}^{off}$ | -0.31 | -0.39 | -0.02 | 0.06 | 0.19 | 0.13 | 0.07 | 0.17 | 0.02 | -0.06 |
| $A_{res}$ | -0.37 | -0.46 | -0.23 | 0.05 | 0.10 | 0.00 | 0.01 | 0.07 | -0.10 | -0.15 |
| $A_{res,\infty}$ | -0.37 | -0.42 | -0.18 | 0.07 | 0.10 | 0.00 | 0.05 | 0.06 | -0.10 | -0.09 |
| $\varepsilon_{off}$ | -0.06 | -0.16 | 0.02 | -0.11 | 0.20 | 0.11 | 0.09 | 0.41 | 0.22 | 0.27 |
| $pKa_{app}$ | -0.17 | -0.28 | -0.05 | -0.19 | 0.05 | -0.03 | -0.06 | 0.19 | 0.03 | -0.02 |
| $A_{unrecov}$ | 0.00 | -0.03 | -0.02 | -0.19 | 0.06 | -0.02 | -0.03 | 0.14 | 0.01 | 0.07 |
| $k_{therm}$ | -0.29 | -0.45 | -0.21 | -0.20 | 0.06 | -0.03 | -0.15 | 0.20 | 0.04 | -0.16 |
| $\varepsilon_{anionic}$ | -0.11 | -0.22 | -0.14 | -0.27 | 0.02 | -0.02 | -0.02 | 0.20 | 0.08 | 0.23 |
| $Q_{fluo}$ | -0.01 | 0.03 | -0.41 | -0.01 | 0.30 | 0.25 | 0.25 | -0.14 | -0.24 | -0.10 |
| M. br | 0.01 | 0.01 | -0.25 | 0.07 | 0.17 | 0.20 | 0.19 | -0.08 | -0.07 | 0.03 |
| $\varepsilon_{overall}$ | 0.00 | -0.01 | 0.04 | 0.07 | -0.07 | 0.02 | 0.02 | -0.03 | 0.09 | 0.13 |
| $Q_{off}$ | 0.34 | 0.52 | 0.19 | -0.06 | -0.04 | 0.03 | 0.02 | -0.02 | 0.08 | 0.22 |
| $\lambda_{ex} - \lambda_{abs}$ | -0.02 | 0.20 | -0.12 | 0.05 | -0.13 | -0.16 | -0.06 | -0.25 | -0.19 | -0.08 |
| $\delta_{Stokes}$ | 0.03 | 0.10 | 0.20 | 0.22 | -0.10 | -0.05 | 0.02 | -0.29 | -0.17 | -0.15 |
| $\Delta pH$ | -0.24 | -0.32 | 0.37 | 0.03 | -0.03 | -0.06 | -0.05 | 0.29 | 0.24 | 0.11 |
| $Q_{on}$ | -0.28 | -0.27 | -0.33 | -0.39 | -0.21 | -0.34 | -0.45 | -0.38 | -0.48 | -0.56 |

| | $Q94_{NE2} - R96_{NH2}$ | $Q94_{NE2} - L69_{CD2}$ | $F145_{CA} - A150_{CB}$ | $F145_{CA} - E222_{CD}$ | $F145_{CA} - F165_{CB}$ | $F145_{CA} - H148_{CA}$ | $F145_{CA} - I167_{CD1}$ | $F145_{CA} - L69_{CD2}$ | $F145_{CA} - L201_{CD1}$ |
|---|---|---|---|---|---|---|---|---|---|
| $\lambda_{abs}$ | 0.28 | 0.42 | 0.03 | 0.07 | 0.09 | 0.23 | 0.20 | 0.14 | 0.16 |
| $\lambda_{ex}$ | 0.36 | 0.30 | -0.04 | 0.12 | -0.07 | 0.27 | 0.06 | 0.02 | 0.10 |
| $\lambda_{em}$ | 0.52 | 0.05 | 0.04 | -0.22 | 0.01 | 0.23 | 0.17 | -0.05 | 0.06 |
| $\lambda_{abs}^{off}$ | 0.23 | 0.18 | 0.03 | -0.45 | 0.24 | 0.07 | 0.35 | 0.03 | -0.07 |
| $A_{res}$ | 0.29 | 0.20 | -0.07 | -0.25 | 0.02 | -0.04 | 0.18 | 0.00 | -0.03 |
| $A_{res,\infty}$ | 0.28 | 0.13 | -0.08 | -0.22 | 0.00 | -0.13 | 0.15 | 0.01 | -0.04 |
| $\varepsilon_{off}$ | 0.25 | 0.43 | -0.14 | 0.11 | -0.11 | 0.02 | -0.04 | 0.07 | 0.01 |
| $pKa_{app}$ | 0.21 | 0.41 | 0.03 | 0.08 | 0.05 | 0.03 | 0.15 | 0.21 | 0.21 |
| $A_{unrecov}$ | 0.18 | 0.30 | -0.03 | 0.17 | -0.06 | -0.09 | -0.08 | 0.13 | 0.15 |
| $k_{therm}$ | 0.20 | 0.45 | 0.14 | -0.17 | 0.17 | 0.45 | 0.26 | 0.10 | 0.12 |
| $\varepsilon_{anionic}$ | 0.18 | 0.18 | 0.14 | 0.35 | 0.11 | 0.05 | 0.11 | 0.29 | 0.21 |
| $Q_{fluo}$ | 0.21 | 0.07 | 0.22 | -0.23 | 0.04 | 0.12 | -0.02 | -0.12 | -0.06 |
| M. br | 0.00 | -0.05 | 0.09 | -0.13 | -0.03 | 0.06 | -0.04 | -0.17 | -0.16 |
| $\varepsilon_{overall}$ | -0.20 | -0.20 | -0.01 | 0.04 | 0.00 | -0.05 | 0.00 | -0.06 | -0.12 |
| $Q_{off}$ | -0.16 | -0.26 | 0.00 | 0.21 | 0.01 | -0.04 | -0.18 | 0.05 | 0.02 |
| $\lambda_{ex} - \lambda_{abs}$ | 0.00 | -0.33 | -0.11 | 0.04 | -0.25 | -0.05 | -0.28 | -0.22 | -0.14 |
| $\delta_{Stokes}$ | -0.14 | -0.39 | 0.08 | -0.33 | 0.11 | -0.22 | 0.04 | -0.06 | -0.10 |
| $\Delta pH$ | 0.05 | 0.24 | -0.17 | -0.01 | -0.02 | -0.11 | 0.08 | 0.08 | -0.04 |
| $Q_{on}$ | 0.25 | -0.13 | 0.63 | -0.19 | 0.61 | 0.49 | 0.58 | 0.40 | 0.51 |

M. br: Molecular brightness; $\Delta pH$: $\lambda_{abs,pH10} - \lambda_{abs,pH6.5}$.

*(Continued)* Pearson correlation coefficients between spectroscopic and on-state structural descriptors.

| | $F145_{CA}-N205_{CA}$ | $F145_{CA}-Q94_{NE2}$ | $F145_{CA}-R96_{NH2}$ | $F145_{CA}-T203_{CB}$ | $N205_{CA}-A150_{CB}$ | $N205_{CA}-E222_{CD}$ | $N205_{CA}-F165_{CB}$ | $N205_{CA}-H148_{CA}$ | $N205_{CA}-I167_{CD1}$ | $N205_{CA}-L69_{CD2}$ | $N205_{CA}-L201_{CD1}$ | $N205_{CA}-Q94_{NE2}$ | $N205_{CA}-R96_{NH2}$ | $N205_{CA}-T203_{CB}$ | $E222_{CD}-A150_{CB}$ | $E222_{CD}-F165_{CB}$ | $E222_{CD}-H148_{CA}$ | $E222_{CD}-I167_{CD1}$ | $E222_{CD}-L69_{CD2}$ |
|---|---|---|---|---|---|---|---|---|---|---|---|---|---|---|---|---|---|---|---|
| $\lambda_{abs}$ | 0.14 | 0.28 | 0.20 | 0.28 | -0.32 | -0.07 | -0.17 | -0.03 | 0.07 | -0.08 | -0.05 | 0.28 | 0.08 | 0.12 | -0.15 | -0.05 | 0.16 | 0.20 | 0.06 |
| $\lambda_{ex}$ | 0.03 | 0.11 | 0.02 | 0.04 | -0.23 | 0.18 | -0.22 | 0.09 | 0.02 | -0.02 | 0.11 | 0.22 | 0.05 | 0.01 | -0.22 | -0.19 | 0.17 | 0.18 | -0.12 |
| $\lambda_{em}$ | -0.04 | 0.06 | -0.04 | -0.06 | -0.04 | -0.09 | -0.03 | 0.10 | 0.17 | 0.21 | 0.17 | 0.40 | 0.24 | 0.01 | -0.12 | -0.06 | -0.20 | -0.15 | 0.23 |
| $\lambda_{abs}^{off}$ | -0.12 | 0.22 | 0.17 | 0.20 | -0.21 | -0.60 | 0.09 | 0.00 | 0.08 | -0.09 | -0.38 | 0.16 | 0.03 | 0.23 | -0.10 | 0.08 | -0.27 | -0.44 | 0.32 |
| $A_{res}$ | 0.08 | 0.23 | 0.13 | 0.26 | -0.39 | -0.50 | -0.08 | 0.01 | 0.08 | -0.25 | -0.35 | 0.14 | -0.03 | 0.32 | -0.18 | -0.06 | -0.10 | -0.19 | 0.16 |
| $A_{res,\infty}$ | 0.11 | 0.23 | 0.11 | 0.23 | -0.36 | -0.46 | -0.04 | 0.00 | 0.10 | -0.21 | -0.33 | 0.13 | -0.02 | 0.32 | -0.14 | -0.02 | -0.06 | -0.12 | 0.16 |
| $\varepsilon_{off}$ | 0.30 | 0.12 | 0.03 | 0.11 | -0.22 | 0.17 | 0.01 | 0.04 | 0.26 | 0.09 | 0.07 | 0.34 | 0.20 | -0.02 | -0.19 | -0.04 | 0.17 | 0.33 | -0.05 |
| $pKa_{app}$ | 0.30 | 0.29 | 0.22 | 0.24 | -0.32 | -0.02 | -0.16 | -0.32 | 0.09 | 0.04 | 0.05 | 0.40 | 0.19 | -0.06 | -0.22 | -0.15 | -0.12 | 0.11 | 0.07 |
| $A_{unrecov}$ | 0.34 | 0.10 | 0.04 | 0.11 | -0.14 | 0.16 | -0.09 | -0.25 | 0.08 | 0.10 | 0.19 | 0.28 | 0.15 | -0.09 | -0.10 | -0.09 | -0.03 | 0.22 | -0.02 |
| $k_{therm}$ | -0.02 | 0.24 | 0.20 | 0.24 | -0.27 | -0.34 | -0.22 | 0.01 | -0.12 | -0.17 | -0.21 | 0.18 | 0.00 | 0.05 | -0.28 | -0.20 | -0.15 | -0.41 | 0.09 |
| $\varepsilon_{anionic}$ | 0.35 | 0.37 | 0.26 | 0.36 | -0.24 | -0.15 | -0.18 | 0.10 | -0.10 | -0.28 | -0.19 | -0.05 | -0.21 | 0.17 | -0.01 | -0.04 | 0.41 | 0.28 | -0.05 |
| $Q_{fluo}$ | -0.06 | -0.20 | -0.24 | -0.01 | 0.38 | -0.22 | 0.16 | 0.27 | 0.12 | -0.03 | 0.16 | -0.16 | -0.19 | 0.10 | 0.36 | 0.36 | 0.17 | 0.18 | 0.21 |
| M. br | -0.16 | -0.20 | -0.20 | -0.04 | 0.26 | -0.20 | 0.12 | 0.44 | 0.05 | -0.20 | -0.07 | -0.38 | -0.31 | 0.22 | 0.26 | 0.28 | 0.31 | 0.13 | 0.03 |
| $\varepsilon_{overall}$ | -0.16 | -0.04 | 0.00 | -0.02 | 0.04 | -0.11 | 0.03 | 0.31 | -0.08 | -0.22 | -0.23 | -0.37 | -0.24 | 0.23 | 0.05 | 0.05 | 0.23 | -0.03 | -0.10 |
| $Q_{off}$ | -0.10 | -0.13 | -0.08 | -0.14 | 0.18 | 0.27 | -0.06 | -0.11 | -0.33 | 0.02 | 0.04 | -0.30 | -0.22 | -0.17 | 0.09 | -0.11 | -0.03 | -0.20 | -0.23 |
| $\lambda_{ex}-\lambda_{abs}$ | -0.20 | -0.34 | -0.32 | -0.44 | 0.24 | 0.36 | 0.00 | 0.17 | -0.08 | 0.12 | 0.24 | -0.19 | -0.08 | -0.20 | -0.04 | -0.16 | -0.05 | -0.11 | -0.26 |
| $\delta_{Stokes}$ | -0.07 | -0.12 | -0.06 | -0.09 | 0.30 | -0.32 | 0.29 | -0.06 | 0.08 | 0.18 | -0.04 | -0.03 | 0.10 | -0.01 | 0.22 | 0.23 | -0.38 | -0.36 | 0.32 |
| $\Delta pH$ | 0.07 | 0.16 | 0.14 | -0.12 | -0.26 | 0.17 | 0.06 | -0.27 | 0.20 | 0.28 | -0.02 | 0.48 | 0.43 | -0.35 | -0.38 | -0.15 | -0.26 | 0.03 | 0.02 |
| $Q_{on}$ | 0.07 | 0.39 | 0.38 | 0.27 | 0.25 | -0.49 | -0.10 | -0.25 | -0.20 | 0.21 | 0.24 | 0.27 | 0.10 | -0.25 | 0.23 | -0.02 | -0.53 | -0.60 | 0.55 |

M. br: Molecular brightness; ΔpH: $\lambda_{abs,pH10}-\lambda_{abs,pH6.5}$.

*(Continued)* Pearson correlation coefficients between spectroscopic and on-state structural descriptors.

| | $E222_{CD} - L201_{CD1}$ | $E222_{CD} - Q94_{NE2}$ | $E222_{CD} - R96_{NH2}$ | $E222_{CD} - T203_{CB}$ | $I167_{CD1} - A150_{CB}$ | $I167_{CD1} - F165_{CB}$ | $I167_{CD1} - H148_{CA}$ | $I167_{CD1} - L69_{CD2}$ | $I167_{CD1} - L201_{CD1}$ | $I167_{CD1} - Q94_{NE2}$ | $I167_{CD1} - R96_{NH2}$ | $I167_{CD1} - T203_{CB}$ | $T203_{CB} - A150_{CB}$ | $T203_{CB} - F165_{CB}$ | $T203_{CB}$-$H148_{CA}$ | $T203_{CB} - L69_{CD2}$ | $T203_{CB} - L201_{CD1}$ | $T203_{CB} - Q94_{NE2}$ | $T203_{CB} - R96_{NH2}$ |
|---|---|---|---|---|---|---|---|---|---|---|---|---|---|---|---|---|---|---|---|
| $\lambda_{abs}$ | 0.06 | 0.25 | 0.11 | 0.06 | -0.02 | -0.12 | -0.01 | 0.13 | 0.09 | 0.42 | 0.17 | 0.24 | -0.29 | -0.14 | 0.17 | -0.19 | -0.17 | 0.14 | 0.00 |
| $\lambda_{ex}$ | -0.10 | 0.07 | -0.02 | -0.04 | -0.02 | -0.12 | 0.06 | 0.04 | 0.14 | 0.21 | -0.04 | -0.01 | -0.22 | -0.28 | -0.06 | -0.02 | 0.07 | 0.15 | -0.04 |
| $\lambda_{em}$ | 0.14 | 0.35 | 0.26 | 0.11 | -0.01 | -0.12 | -0.09 | -0.35 | -0.02 | -0.19 | -0.47 | -0.09 | -0.11 | -0.18 | -0.26 | 0.08 | 0.12 | 0.18 | -0.04 |
| $\lambda_{abs}^{off}$ | 0.20 | 0.29 | 0.11 | 0.14 | -0.22 | -0.08 | -0.04 | -0.25 | -0.41 | 0.07 | -0.11 | 0.13 | -0.25 | 0.05 | 0.13 | -0.18 | -0.38 | 0.10 | 0.00 |
| $A_{res}$ | 0.10 | 0.23 | 0.06 | 0.08 | -0.09 | -0.12 | -0.05 | -0.11 | -0.09 | 0.15 | -0.08 | 0.20 | -0.47 | -0.15 | 0.13 | -0.32 | -0.43 | -0.01 | -0.15 |
| $A_{res,\infty}$ | 0.10 | 0.20 | 0.06 | 0.06 | -0.08 | -0.12 | -0.11 | -0.04 | -0.07 | 0.19 | -0.05 | 0.18 | -0.46 | -0.13 | 0.15 | -0.30 | -0.42 | -0.04 | -0.17 |
| $\varepsilon_{off}$ | -0.06 | 0.12 | -0.01 | -0.16 | -0.02 | -0.11 | 0.13 | 0.44 | 0.21 | 0.62 | 0.36 | 0.45 | -0.12 | 0.09 | 0.24 | 0.04 | 0.00 | 0.30 | 0.20 |
| $pKa_{app}$ | 0.07 | 0.28 | 0.07 | -0.11 | 0.03 | -0.10 | 0.01 | 0.19 | 0.12 | 0.36 | 0.16 | 0.29 | -0.21 | -0.05 | -0.01 | -0.02 | -0.01 | 0.25 | 0.11 |
| $A_{unrecov}$ | 0.02 | 0.14 | 0.04 | -0.18 | 0.13 | 0.05 | 0.07 | 0.28 | 0.28 | 0.27 | 0.13 | 0.21 | -0.08 | -0.06 | 0.00 | 0.00 | 0.10 | 0.13 | 0.02 |
| $k_{therm}$ | 0.08 | 0.26 | 0.01 | 0.25 | 0.04 | -0.02 | 0.27 | -0.27 | -0.24 | -0.07 | -0.18 | 0.09 | -0.21 | -0.15 | -0.08 | -0.03 | -0.11 | 0.31 | 0.16 |
| $\varepsilon_{anionic}$ | 0.00 | 0.05 | 0.00 | 0.02 | 0.19 | 0.01 | -0.15 | 0.27 | 0.09 | 0.45 | 0.23 | 0.12 | -0.28 | -0.19 | 0.34 | -0.23 | -0.27 | -0.06 | -0.13 |
| $Q_{fluo}$ | 0.33 | 0.03 | 0.07 | -0.01 | 0.35 | 0.05 | -0.33 | 0.15 | -0.01 | 0.14 | -0.01 | -0.01 | 0.18 | 0.03 | 0.34 | -0.29 | -0.11 | -0.31 | -0.41 |
| M. br | 0.13 | -0.16 | -0.04 | 0.09 | 0.19 | -0.01 | -0.22 | 0.08 | -0.06 | 0.07 | 0.05 | -0.07 | 0.05 | -0.02 | 0.38 | -0.31 | -0.22 | -0.35 | -0.35 |
| $\varepsilon_{overall}$ | -0.10 | -0.23 | -0.10 | 0.16 | -0.04 | -0.02 | -0.07 | -0.07 | -0.13 | -0.09 | 0.03 | -0.18 | -0.10 | -0.09 | 0.14 | -0.14 | -0.21 | -0.23 | -0.13 |
| $Q_{off}$ | -0.22 | -0.33 | -0.27 | -0.18 | -0.04 | 0.26 | 0.19 | -0.08 | -0.05 | -0.33 | -0.15 | -0.35 | 0.24 | -0.05 | -0.16 | 0.19 | 0.19 | -0.09 | -0.04 |
| $\lambda_{ex} - \lambda_{abs}$ | -0.25 | -0.33 | -0.22 | -0.16 | 0.01 | 0.04 | 0.11 | -0.18 | 0.04 | -0.45 | -0.35 | -0.43 | 0.22 | -0.12 | -0.38 | 0.31 | 0.39 | -0.05 | -0.04 |
| $\delta_{Stokes}$ | 0.24 | 0.14 | 0.21 | 0.14 | 0.03 | 0.09 | -0.15 | -0.29 | -0.21 | -0.42 | -0.26 | -0.05 | 0.23 | 0.26 | -0.09 | 0.09 | -0.02 | -0.09 | 0.02 |
| $\Delta pH$ | -0.17 | 0.23 | 0.12 | -0.24 | -0.24 | -0.14 | 0.20 | 0.13 | -0.01 | 0.27 | 0.19 | 0.28 | 0.01 | 0.27 | -0.20 | 0.41 | 0.25 | 0.52 | 0.55 |
| $Q_{on}$ | 0.55 | 0.49 | 0.36 | 0.08 | 0.17 | 0.04 | -0.49 | -0.62 | -0.40 | -0.36 | -0.48 | -0.31 | 0.27 | -0.04 | -0.23 | 0.11 | 0.21 | 0.11 | 0.00 |

M. br: Molecular brightness; ΔpH: $\lambda_{abs,pH10} - \lambda_{abs,pH6.5}$.

**Table S7:** Principal component analysis on spectroscopic and on-state structural properties indicating the FP's structural properties compared to average.

| | **Zone1**† | **Zone 2**$,† | **Fast switchers**#,† |
|---|---|---|---|
| Chromophore descriptors | | **Tilt** ↑ | |
| Chromophore and environment based descriptors | Total No. of heteroatoms around chromophore* ↓<br>**Total No. of H-bonds Hydroxybenz$_{OH}$** ↓<br>Total No. of H-bonds Imidazolinone$_{N}$ ↑<br>Total No. of H-bonds Imidazolinone$_{O}$ ~↑<br>**Hydroxybenz$_{OH}$ – H148$_{CA}$** ↓<br>Hydroxybenz$_{OH}$ – I167$_{CD1}$ ~↑<br>**Hydroxybenz$_{OH}$ – N205$_{OD1/ND2/HOH}$** ↓<br>Hydroxybenz$_{planecenter}$ – E222$_{CD}$ ↑<br>Hydroxybenz$_{planecenter}$ – T62$_{CG2}$ ↑<br>Imidazolinone$_{O}$ – L69$_{CD2}$ ~↓ | **Hydroxybenz$_{OH}$ – A150$_{CB}$** ↑<br>**Hydroxybenz$_{OH}$ – F145$_{CA}$** ↓<br>**Hydroxybenz$_{OH}$ – F145$_{CB}$** ↓<br>**Hydroxybenz$_{OH}$ – H148$_{CA}$** ↑<br>**Hydroxybenz$_{OH}$ – L201$_{CD1}$** ↑<br>**Hydroxybenz$_{OH}$ – N205$_{CA}$** ~↓<br>**Hydroxybenz$_{OH}$ – N205$_{OD1/ND2}$** ↓<br>**Hydroxybenz$_{planecenter}$ – F165$_{planecenter}$** ↑<br>**Hydroxybenz$_{planecenter}$ – R96$_{NH2}$** ~↑<br>**Hydroxybenz$_{planecenter}$ – L69$_{CD2}$** ↓ | **Total No. of H-bonds** ↑§<br>**Total No. of H-bonds Hydroxybenz$_{OH}$** ↑<br>Total No. of H-bonds Imidazolinone$_{N}$ ~↓<br>**Total No. of H-bonds Imidazolinone$_{O}$** ~↑<br>**Hydroxybenz$_{OH}$ – F145$_{CA}$** ~↓<br>Hydroxybenz$_{OH}$ – H148$_{CA}$ ↑<br>**Hydroxybenz$_{OH}$ – N205$_{CA}$** ~↓<br>**Hydroxybenz$_{OH}$ – N205$_{OD1/ND2}$** ~↓<br>Hydroxybenz$_{planecenter}$ – E222$_{CD}$ ↓<br>**Hydroxybenz$_{planecenter}$ – R96$_{NH2}$** ~↑<br>Hydroxybenz$_{planecenter}$ – T62$_{CD2}$ ↓<br>Imidazolinone$_{O}$ – R96$_{NH1}$ ↓ |
| Chromophore independent descriptors | F165$_{CB}$ – H148$_{CA}$ ↓<br>H148$_{CA}$ – Q94$_{NE2}$ ~↓<br>H148$_{CA}$ – R96$_{NH2}$ ↓<br>Q94$_{NE2}$ – R96$_{NH2}$ ↑<br>**F145$_{CA}$ –E222$_{CD}$** ↓<br>**N205$_{CA}$ – E222$_{CD}$** ↓<br>E222$_{CD}$ – T203$_{CB}$ ~↑<br>I167$_{CD1}$ – H148$_{CA}$ ↓ | **A150$_{CB}$ – L201$_{CD1}$** ↑<br>**A150$_{CB}$ – L69$_{CD2}$** ↑<br>**H148$_{CA}$ – Q94$_{NE2}$** ↓<br>**H148$_{CA}$ – R96$_{NH2}$** ↓<br>**F145$_{CA}$ – Q94$_{NE2}$** ~↓<br>**F145$_{CA}$ – R96$_{NH2}$** ~↓<br>**N205$_{CA}$ – Q94$_{NE2}$** ↓<br>**I176$_{CD1}$ – Q94$_{NE2}$** ↓<br>**I167$_{CD1}$ – R96$_{NH2}$** ↓ | A150$_{CB}$ – L201$_{CD1}$ ~↑<br>A150$_{CB}$ – R96$_{NH2}$ ↓<br>F145$_{CA}$ – H148$_{CA}$ ↑<br>F145$_{CA}$ – Q94$_{NE2}$ ~↓<br>F145$_{CA}$ – R96$_{NH2}$ ~↓<br>**E222$_{CD}$ – F165$_{CB}$** ~↑<br>E222$_{CD}$ – I167$_{CD1}$ ↓<br>**E222$_{CD}$ – T203$_{CB}$** ~↑<br>I167$_{CD1}$ – L69$_{CD2}$ ↓<br>I167$_{CD1}$ – L201$_{CD1}$ ~↓<br>**I167$_{CD1}$ – Q94$_{NE2}$** ↓<br>**I167$_{CD1}$ – R96$_{NH2}$** ↓<br>I167$_{CD1}$ – T203$_{CB}$ ↓<br>T203$_{CB}$ – F165$_{CB}$ ↓<br>**T203$_{CB}$ – H148$_{CA}$** ↓ |

↑: higher than average, ↓: lower than average, ~: equal to average

† Zone 1: PC2+, PC4-, PC5+; Zone 2: PC1-, PC2-; Fast switchers: PC1-, PC5-, PC9+

$ Zone 2 can only be differentiated from the other FPs using only two PCs (with exceptions in PC2). Only those variables in equal sign in the loadings and are within the 15 highest of PC1 or PC2 (Supplementary Table S5) are listed.

### See Supplementary Table S1 for FP classification.

§ Descriptors in bold are within the 15 highest loadings of the relevant PCs (Supplementary Table S5).

*Total No. of atoms within a sphere of 3.5 Å

**Table S8:** 15 highest loadings (positive or negative) in decreasing order for LV1 and LV2 in the PLS-DA with the spectroscopic and off-state structural descriptors.

| LV1 | LV2 |
|---|---|
| $E222_{CD}$ – L201CD1 | $Hydroxybenz_{OH}$ – $N205_{OD1/ND2}$* |
| $N205_{CA}$ – $L201_{CD1}$ | $A150_{CB}$ – $L201_{CD1}$ |
| $N205_{CA}$ – $F165_{CB}$ | $L201_{CD1}$ – $Q94_{NE2}$ |
| $H148_{CA}$ – $L69_{CD2}$ | $\lambda_{ex} - \lambda_{abs}$ |
| $A150_{CB}$ – $L69_{CD2}$ | $\varepsilon_{off}$ |
| $I167_{CD1}$ – $L69_{CD2}$ | $Hydroxybenz_{OH}$ – $N205_{CA}$ |
| $F165_{CB}$ – $L69_{CD2}$ | $I167_{CD1}$ – $A150_{CB}$ |
| $E222_{CD}$ – $F165_{CB}$ | $\varepsilon_{anionic}$ |
| $T203_{CB}$ – $F165_{CB}$ | $F165_{CB}$ – $H148_{CA}$ |
| $T203_{CB}$ – $L201_{CD1}$ | $Hydroxybenz_{OH}$ – $H148_{CA}$ |
| $I167_{CD1}$ – $F165_{CB}$ | $E222_{CD}$ – $I167_{CD1}$ |
| $E222_{CD}$ – $H148_{CA}$ | $Q_{off}$ |
| $Hydroxybenz_{OH}$ – $F145_{CA}$ | $\lambda_{abs}$ |
| $T203_{CB}$ – $A150_{CB}$ | $Q94_{NE2}$ – $R96_{NH2}$ |
| $I167_{CD1}$ – $L201_{CD1}$ | $E222_{CD}$ – $A150_{CB}$ |

*Atom that is closest to chromophore anchor point..

**Table S9:** Pearson correlation coefficients between spectroscopic and off-state structural descriptors.

| | methylene bond angle | tilt | twist | Pocket volume | Total No. H-bonds | Total No. H-bonds $Hydroxybenz_{OH}$ | Total No. H-bonds $Imidazolinone_{N}$ | Total No. H-bonds $Imidazolinone_{O}$ | Total No. of heteroatoms $Hydroxybenz_{OH}$ | Total No. of atoms $Hydroxybenz_{plane}$ | Total No. of atoms $Imidazolinone_{plane}$ | Total No. of heteroatoms $Imidazolinone_{O}$ | Total No. water molecules | Total No. of heteroatoms | $Hydroxybenz_{OH}$ – $F145_{CA}$ | $Hydroxybenz_{OH}$ – $F145_{CB}$ | $Hydroxybenz_{OH}$ – $H148_{CA}$ | $Hydroxybenz_{OH}$ – $H148_{planecenter}$ | $Hydroxybenz_{OH}$ – $I167_{CD1}$ | $Hydroxybenz_{OH}$ – $N205_{CA}$ |
|---|---|---|---|---|---|---|---|---|---|---|---|---|---|---|---|---|---|---|---|---|
| $\lambda_{abs}$ | 0.01 | -0.32 | 0.48 | -0.76 | 0.13 | 0.23 | -0.35 | 0.00 | -0.59 | -0.05 | 0.35 | 0.18 | -0.50 | -0.30 | -0.07 | -0.09 | -0.34 | 0.16 | -0.28 | -0.39 |
| $\lambda_{ex}$ | -0.25 | 0.05 | -0.03 | -0.48 | 0.14 | 0.09 | -0.07 | 0.17 | -0.38 | -0.26 | 0.46 | 0.14 | -0.46 | -0.01 | -0.33 | -0.33 | -0.23 | -0.09 | -0.42 | -0.06 |
| $\lambda_{em}$ | -0.31 | 0.25 | -0.19 | -0.29 | 0.56 | 0.47 | 0.00 | 0.45 | -0.21 | -0.21 | 0.11 | -0.06 | -0.35 | 0.03 | -0.28 | -0.34 | -0.30 | -0.04 | -0.31 | 0.12 |
| $\lambda_{abs}^{off}$ | -0.40 | -0.17 | 0.15 | 0.21 | -0.18 | -0.14 | 0.00 | -0.16 | 0.07 | 0.19 | 0.12 | -0.39 | 0.12 | -0.15 | 0.60 | 0.63 | 0.58 | 0.48 | 0.02 | 0.11 |
| $A_{res}$ | -0.17 | 0.37 | -0.03 | -0.32 | 0.38 | 0.41 | 0.10 | 0.17 | -0.16 | -0.08 | 0.07 | 0.02 | -0.18 | -0.16 | 0.02 | 0.00 | -0.32 | 0.08 | 0.25 | -0.21 |
| $A_{res,\infty}$ | -0.11 | 0.38 | -0.04 | -0.30 | 0.39 | 0.42 | 0.07 | 0.17 | -0.14 | -0.08 | 0.05 | 0.06 | -0.19 | -0.15 | 0.00 | -0.02 | -0.37 | 0.06 | 0.24 | -0.21 |
| $\varepsilon_{off}$ | 0.30 | 0.08 | 0.17 | -0.40 | 0.42 | 0.50 | -0.26 | 0.19 | -0.29 | -0.03 | -0.07 | 0.12 | -0.21 | -0.20 | -0.04 | -0.08 | -0.66 | 0.03 | 0.00 | -0.19 |
| $pKa_{app}$ | 0.04 | 0.21 | 0.19 | -0.39 | 0.61 | 0.66 | -0.08 | 0.30 | -0.19 | -0.12 | -0.16 | 0.16 | -0.10 | -0.08 | -0.05 | -0.15 | -0.70 | 0.04 | 0.04 | -0.18 |
| $A_{unrecov}$ | 0.19 | 0.24 | 0.08 | -0.23 | 0.61 | 0.62 | -0.12 | 0.37 | -0.11 | -0.18 | -0.31 | 0.12 | -0.05 | 0.02 | -0.22 | -0.34 | -0.74 | -0.10 | -0.09 | -0.05 |
| $k_{therm}$ | -0.38 | 0.31 | -0.05 | -0.26 | 0.24 | 0.26 | 0.21 | 0.07 | -0.25 | -0.19 | 0.13 | -0.19 | -0.12 | -0.23 | 0.12 | 0.11 | -0.02 | 0.23 | 0.19 | -0.02 |
| $\varepsilon_{anionic}$ | 0.19 | -0.73 | 0.63 | 0.09 | -0.23 | -0.07 | -0.46 | -0.28 | -0.38 | 0.14 | 0.37 | 0.30 | -0.32 | -0.24 | 0.17 | 0.22 | -0.16 | 0.05 | -0.21 | -0.58 |
| $Q_{fluo}$ | 0.17 | 0.23 | -0.11 | -0.33 | -0.31 | -0.28 | 0.03 | -0.22 | -0.60 | -0.49 | 0.49 | 0.37 | -0.66 | -0.24 | -0.52 | -0.47 | -0.27 | -0.12 | -0.24 | -0.08 |
| M. br | 0.10 | -0.18 | 0.03 | -0.16 | -0.58 | -0.54 | -0.01 | -0.40 | -0.45 | -0.20 | 0.51 | 0.15 | -0.37 | -0.21 | -0.16 | -0.07 | 0.23 | 0.03 | -0.19 | 0.01 |
| $\varepsilon_{overall}$ | -0.05 | -0.49 | 0.14 | 0.47 | -0.50 | -0.47 | -0.04 | -0.32 | 0.00 | 0.27 | 0.31 | -0.14 | 0.18 | -0.01 | 0.30 | 0.40 | 0.61 | 0.18 | -0.02 | 0.08 |
| $Q_{off}$ | 0.11 | 0.15 | -0.34 | 0.75 | -0.05 | -0.17 | 0.10 | 0.12 | 0.61 | 0.09 | -0.38 | -0.13 | 0.48 | 0.42 | 0.03 | 0.04 | 0.17 | -0.24 | -0.06 | 0.37 |
| $\lambda_{ex} - \lambda_{abs}$ | -0.21 | 0.39 | -0.54 | 0.47 | -0.04 | -0.18 | 0.32 | 0.13 | 0.36 | -0.14 | -0.03 | -0.09 | 0.19 | 0.32 | -0.17 | -0.15 | 0.19 | -0.24 | -0.01 | 0.37 |
| $\delta_{Stokes}$ | 0.05 | 0.19 | -0.15 | 0.43 | 0.39 | 0.38 | 0.11 | 0.23 | 0.36 | 0.17 | -0.60 | -0.30 | 0.34 | 0.04 | 0.21 | 0.13 | 0.03 | 0.10 | 0.31 | 0.23 |
| $\Delta pH$ | -0.43 | -0.24 | 0.35 | -0.37 | 0.17 | 0.12 | 0.02 | 0.16 | 0.05 | 0.33 | -0.01 | -0.15 | 0.07 | 0.03 | 0.30 | 0.30 | 0.22 | -0.01 | 0.19 | -0.38 |
| $Q_{on}$ | -0.44 | -0.08 | -0.01 | -0.50 | 0.26 | 0.18 | -0.21 | 0.32 | 0.19 | 0.17 | 0.33 | 0.07 | 0.02 | 0.28 | -0.08 | -0.14 | -0.17 | -0.24 | -0.16 | -0.24 |

M. br: Molecular brightness; ΔpH: $\lambda_{abs,pH10} - \lambda_{abs,pH6.5}$.

*(Continued)* Pearson correlation coefficients between spectroscopic and off-state structural descriptors.

| | Hydroxybenz$_{OH}$ – N205$_{OD1/ND2}$* | Hydroxybenz$_{OH}$ – HOH* | Hydroxybenz$_{planecenter}$ – H148$_{planecenter}$ | Hydroxybenz$_{planecenter}$ – T62$_{CG2}$ | Imidazolinone$_{O}$ – L69$_{CD2}$ | Imidazolinone$_{O}$ – R96$_{NH1}$ | A150$_{CB}$ – F165$_{CB}$ | A150$_{CB}$ – L201$_{CD1}$ | A150$_{CB}$ – Q94$_{NE2}$ | A150$_{CB}$ – R96$_{NH2}$ | A150$_{CB}$ – L69$_{CD2}$ | F165$_{CB}$ – H148$_{CA}$ | F165$_{CB}$ – L201$_{CD1}$ | F165$_{CB}$ – Q94$_{NE2}$ | F165$_{CB}$ – R96$_{NH2}$ | F165$_{CB}$ – L69$_{CD2}$ | H148$_{CA}$ – Q94$_{NE2}$ | H148$_{CA}$ – R96$_{NH2}$ | H148$_{CA}$ – L69$_{CD2}$ |
|---|---|---|---|---|---|---|---|---|---|---|---|---|---|---|---|---|---|---|---|
| $\lambda_{abs}$ | -0.61 | 0.10 | 0.21 | -0.38 | 0.03 | -0.03 | -0.01 | -0.42 | 0.13 | -0.26 | -0.25 | -0.26 | -0.28 | 0.08 | -0.25 | -0.21 | -0.13 | -0.35 | -0.36 |
| $\lambda_{ex}$ | 0.00 | -0.07 | -0.10 | -0.23 | -0.14 | -0.33 | -0.40 | 0.16 | -0.14 | -0.25 | 0.05 | -0.01 | -0.29 | -0.44 | -0.53 | -0.40 | -0.24 | -0.19 | -0.22 |
| $\lambda_{em}$ | 0.17 | 0.28 | -0.07 | -0.15 | -0.04 | -0.47 | -0.17 | 0.06 | -0.16 | -0.47 | -0.02 | -0.04 | -0.14 | -0.31 | -0.60 | -0.27 | -0.18 | -0.33 | -0.25 |
| $\lambda^{off}_{abs}$ | -0.02 | -0.24 | 0.48 | 0.14 | 0.09 | 0.25 | 0.05 | 0.42 | -0.18 | -0.07 | -0.39 | 0.34 | 0.35 | 0.15 | 0.23 | 0.06 | 0.43 | 0.37 | -0.03 |
| $A_{res}$ | -0.34 | 0.58 | 0.08 | 0.28 | 0.21 | -0.39 | 0.51 | -0.25 | -0.26 | -0.68 | -0.60 | -0.45 | 0.40 | 0.29 | -0.13 | 0.06 | -0.19 | -0.53 | -0.50 |
| $A_{res,\infty}$ | -0.34 | 0.65 | 0.06 | 0.27 | 0.26 | -0.43 | 0.53 | -0.28 | -0.20 | -0.67 | -0.56 | -0.47 | 0.41 | 0.34 | -0.10 | 0.10 | -0.18 | -0.55 | -0.48 |
| $\varepsilon_{off}$ | -0.42 | 0.56 | 0.06 | 0.04 | 0.22 | -0.22 | 0.35 | -0.56 | 0.16 | -0.41 | -0.32 | -0.53 | 0.01 | 0.29 | -0.21 | -0.02 | -0.33 | -0.70 | -0.48 |
| $pKa_{app}$ | -0.18 | 0.58 | 0.01 | -0.03 | 0.08 | -0.43 | 0.42 | -0.61 | -0.08 | -0.58 | -0.42 | -0.72 | 0.07 | 0.25 | -0.23 | -0.04 | -0.56 | -0.79 | -0.65 |
| $A_{unrecov}$ | 0.01 | 0.48 | -0.14 | -0.04 | 0.06 | -0.40 | 0.29 | -0.57 | 0.02 | -0.37 | -0.16 | -0.61 | -0.07 | 0.10 | -0.27 | -0.06 | -0.59 | -0.72 | -0.52 |
| $k_{therm}$ | -0.13 | 0.07 | 0.19 | 0.36 | -0.05 | -0.16 | 0.24 | 0.09 | -0.53 | -0.66 | -0.69 | -0.24 | 0.36 | -0.07 | -0.27 | -0.21 | -0.30 | -0.37 | -0.60 |
| $\varepsilon_{anionic}$ | -0.71 | -0.18 | 0.12 | -0.50 | -0.19 | 0.40 | -0.30 | -0.36 | 0.49 | 0.23 | 0.06 | -0.17 | -0.52 | 0.02 | -0.14 | -0.29 | 0.06 | -0.12 | -0.02 |
| $Q_{fluo}$ | -0.27 | 0.02 | -0.06 | 0.18 | 0.29 | -0.23 | 0.07 | 0.13 | -0.04 | -0.06 | 0.02 | 0.01 | 0.15 | 0.03 | 0.03 | 0.09 | -0.04 | -0.05 | -0.13 |
| M. br | -0.23 | -0.48 | 0.09 | 0.08 | 0.02 | 0.30 | -0.27 | 0.34 | 0.01 | 0.27 | 0.11 | 0.37 | -0.06 | -0.19 | 0.06 | -0.09 | 0.17 | 0.33 | 0.14 |
| $\varepsilon_{overall}$ | -0.04 | -0.66 | 0.19 | -0.09 | -0.30 | 0.61 | -0.49 | 0.35 | 0.06 | 0.43 | 0.17 | 0.49 | -0.28 | -0.31 | 0.04 | -0.23 | 0.29 | 0.52 | 0.37 |
| $Q_{off}$ | 0.71 | -0.14 | -0.28 | 0.08 | -0.08 | -0.01 | -0.25 | 0.38 | 0.09 | 0.45 | 0.51 | 0.27 | -0.03 | -0.20 | 0.15 | 0.09 | 0.06 | 0.34 | 0.40 |
| $\lambda_{ex} - \lambda_{abs}$ | 0.66 | -0.16 | -0.30 | 0.24 | -0.14 | -0.21 | -0.28 | 0.57 | -0.25 | 0.10 | 0.30 | 0.27 | 0.08 | -0.41 | -0.12 | -0.08 | -0.04 | 0.24 | 0.22 |
| $\delta_{Stokes}$ | 0.18 | 0.42 | 0.09 | 0.20 | 0.17 | -0.01 | 0.43 | -0.18 | 0.04 | -0.14 | -0.10 | -0.02 | 0.30 | 0.33 | 0.17 | 0.33 | 0.18 | -0.06 | 0.08 |
| $\Delta pH$ | -0.38 | 0.03 | -0.04 | -0.19 | -0.07 | 0.07 | 0.06 | -0.14 | -0.26 | -0.34 | -0.55 | -0.10 | 0.03 | -0.02 | -0.15 | -0.25 | 0.01 | -0.10 | -0.27 |
| $Q_{on}$ | 0.01 | -0.02 | -0.30 | -0.40 | -0.27 | -0.29 | -0.44 | -0.13 | 0.00 | -0.19 | 0.04 | -0.24 | -0.52 | -0.45 | -0.61 | -0.54 | -0.28 | -0.28 | -0.10 |

M. br: Molecular brightness; ΔpH: $\lambda_{abs,pH10} - \lambda_{abs,pH6.5}$.

*Atom that is closest to chromophore anchor point.

*(Continued)* Pearson correlation coefficients between spectroscopic and off-state structural descriptors.

| | $L201_{CD1}-Q94_{NE2}$ | $L201_{CD1}-L69_{CD2}$ | $Q94_{NE2}-R96_{NH2}$ | $Q94_{NE2}-L69_{CD2}$ | $F145_{CA}-A150_{CB}$ | $F145_{CA}-E222_{CD}$ | $F145_{CA}-F165_{CB}$ | $F145_{CA}-H148_{CA}$ | $F145_{CA}-I167_{CD1}$ | $F145_{CA}-L69_{CD2}$ | $F145_{CA}-L201_{CD1}$ | $F145_{CA}-Q94_{NE2}$ | $F145_{CA}-R96_{NH2}$ | $F145_{CA}-T203_{CB}$ | $N205_{CA}-A150_{CB}$ | $N205_{CA}-F165_{CB}$ | $N205_{CA}-I167_{CD1}$ | $N205_{CA}-L69_{CD2}$ | $N205_{CA}-L201_{CD1}$ |
|---|---|---|---|---|---|---|---|---|---|---|---|---|---|---|---|---|---|---|---|
| $\lambda_{abs}$ | 0.03 | -0.06 | 0.51 | 0.08 | 0.08 | 0.42 | -0.12 | -0.21 | -0.07 | 0.07 | 0.08 | 0.10 | -0.02 | 0.14 | -0.06 | -0.30 | -0.25 | -0.42 | -0.08 |
| $\lambda_{ex}$ | -0.33 | 0.17 | 0.16 | -0.36 | -0.31 | -0.10 | -0.47 | -0.48 | -0.32 | -0.39 | -0.28 | -0.28 | -0.30 | -0.37 | -0.02 | -0.32 | -0.22 | -0.35 | 0.06 |
| $\lambda_{em}$ | -0.25 | 0.18 | 0.47 | -0.32 | -0.32 | -0.25 | -0.46 | -0.45 | -0.17 | -0.39 | -0.36 | -0.19 | -0.31 | -0.41 | 0.01 | -0.19 | 0.05 | -0.15 | 0.05 |
| $\lambda^{off}_{abs}$ | -0.05 | -0.62 | -0.11 | 0.29 | 0.06 | 0.29 | 0.61 | 0.76 | 0.66 | 0.47 | 0.34 | 0.57 | 0.57 | 0.69 | -0.68 | -0.06 | 0.02 | -0.21 | -0.55 |
| $A_{res}$ | 0.32 | -0.11 | 0.66 | 0.15 | 0.03 | 0.04 | -0.17 | -0.02 | 0.08 | -0.08 | -0.05 | 0.07 | -0.13 | 0.12 | -0.14 | -0.19 | 0.05 | -0.23 | -0.21 |
| $A_{res,\infty}$ | 0.37 | -0.08 | 0.69 | 0.19 | 0.06 | 0.03 | -0.15 | 0.00 | 0.07 | -0.07 | -0.04 | 0.07 | -0.13 | 0.12 | -0.07 | -0.12 | 0.09 | -0.18 | -0.17 |
| $\varepsilon_{off}$ | 0.41 | 0.02 | 0.79 | 0.20 | 0.34 | 0.21 | -0.05 | -0.06 | 0.04 | 0.09 | 0.13 | 0.08 | -0.13 | 0.09 | 0.37 | 0.04 | 0.16 | 0.09 | 0.12 |
| $pKa_{app}$ | 0.22 | 0.08 | 0.70 | -0.04 | 0.20 | 0.07 | -0.22 | -0.33 | -0.02 | -0.05 | -0.06 | 0.08 | -0.10 | -0.06 | 0.18 | -0.18 | -0.02 | -0.04 | -0.05 |
| $A_{unrecov}$ | 0.11 | 0.22 | 0.55 | -0.13 | 0.14 | -0.10 | -0.33 | -0.51 | -0.14 | -0.20 | -0.18 | -0.11 | -0.24 | -0.30 | 0.41 | -0.04 | 0.11 | 0.12 | 0.16 |
| $k_{therm}$ | -0.05 | -0.29 | 0.35 | -0.05 | -0.11 | -0.01 | -0.20 | -0.01 | 0.12 | -0.15 | -0.06 | 0.02 | -0.10 | 0.15 | -0.37 | -0.39 | -0.03 | -0.38 | -0.32 |
| $\varepsilon_{anionic}$ | 0.27 | 0.08 | 0.27 | 0.11 | 0.48 | 0.78 | 0.23 | 0.15 | -0.02 | 0.48 | 0.56 | 0.32 | 0.26 | 0.40 | 0.02 | -0.42 | -0.56 | -0.33 | -0.04 |
| $Q_{fluo}$ | 0.08 | 0.12 | 0.00 | 0.21 | -0.32 | -0.19 | -0.42 | -0.29 | -0.38 | -0.50 | -0.26 | -0.49 | -0.49 | -0.26 | 0.27 | 0.26 | 0.18 | -0.24 | 0.39 |
| M. br | -0.10 | -0.08 | -0.35 | 0.14 | -0.17 | 0.06 | -0.04 | 0.09 | -0.16 | -0.13 | 0.06 | -0.25 | -0.16 | 0.06 | 0.03 | 0.14 | 0.01 | -0.20 | 0.23 |
| $\varepsilon_{overall}$ | -0.25 | -0.20 | -0.50 | -0.07 | 0.09 | 0.29 | 0.38 | 0.43 | 0.14 | 0.32 | 0.34 | 0.15 | 0.29 | 0.34 | -0.23 | -0.08 | -0.21 | -0.05 | -0.09 |
| $Q_{off}$ | -0.14 | 0.16 | -0.54 | -0.20 | 0.02 | -0.35 | 0.19 | 0.11 | 0.06 | 0.00 | -0.06 | -0.05 | 0.11 | -0.25 | 0.23 | 0.31 | 0.19 | 0.48 | 0.13 |
| $\lambda_{ex}-\lambda_{abs}$ | -0.28 | 0.19 | -0.43 | -0.36 | -0.32 | -0.53 | -0.22 | -0.13 | -0.16 | -0.36 | -0.29 | -0.32 | -0.20 | -0.43 | 0.06 | 0.08 | 0.11 | 0.19 | 0.14 |
| $\delta_{Stokes}$ | 0.24 | -0.07 | 0.27 | 0.21 | 0.13 | -0.11 | 0.23 | 0.25 | 0.31 | 0.18 | 0.04 | 0.22 | 0.13 | 0.12 | 0.04 | 0.29 | 0.39 | 0.39 | -0.05 |
| $\Delta pH$ | 0.00 | -0.32 | 0.23 | -0.03 | 0.06 | 0.35 | 0.07 | 0.02 | 0.22 | 0.25 | 0.13 | 0.32 | 0.23 | 0.30 | -0.62 | -0.64 | -0.38 | -0.45 | -0.58 |
| $Q_{on}$ | -0.24 | 0.24 | 0.25 | -0.57 | -0.06 | 0.10 | -0.35 | -0.45 | -0.26 | -0.04 | -0.06 | -0.02 | -0.05 | -0.30 | -0.17 | -0.70 | -0.54 | -0.15 | -0.12 |

M. br: Molecular brightness; ΔpH: $\lambda_{abs,pH10} - \lambda_{abs,pH6.5}$.

*(Continued)* Pearson correlation coefficients between spectroscopic and off-state structural descriptors.

| | $N205_{CA} - Q94_{NE2}$ | $N205_{CA} - R96_{NH2}$ | $E222_{CD} - A150_{CB}$ | $E222_{CD} - F165_{CB}$ | $E222_{CD} - H148_{CA}$ | $E222_{CD} - I167_{CD1}$ | $E222_{CD} - L201_{CD1}$ | $I167_{CD1} - A150_{CB}$ | $I167_{CD1} - F165_{CB}$ | $I167_{CD1} - H148_{CA}$ | $I167_{CD1} - L69_{CD2}$ | $I167_{CD1} - L201_{CD1}$ | $I167_{CD1} - R96_{NH2}$ | $I167_{CD1} - T203_{CB}$ | $T203_{CB} - A150_{CB}$ | $T203_{CB} - F165_{CB}$ | $T203_{CB} - H148_{CA}$ | $T203_{CB} - L69_{CD2}$ | $T203_{CB} - L201_{CD1}$ |
|---|---|---|---|---|---|---|---|---|---|---|---|---|---|---|---|---|---|---|---|
| $\lambda_{abs}$ | -0.42 | -0.52 | 0.14 | 0.03 | 0.20 | 0.10 | 0.05 | 0.06 | -0.09 | -0.34 | -0.08 | -0.15 | -0.11 | -0.14 | 0.00 | -0.17 | 0.05 | -0.37 | -0.12 |
| $\lambda_{ex}$ | -0.26 | -0.17 | -0.01 | -0.37 | -0.08 | 0.01 | -0.05 | -0.10 | -0.21 | -0.16 | -0.37 | -0.11 | -0.22 | -0.36 | -0.01 | -0.39 | -0.32 | 0.02 | 0.22 |
| $\lambda_{em}$ | 0.06 | 0.04 | -0.17 | -0.41 | -0.28 | -0.20 | -0.14 | -0.17 | -0.41 | -0.14 | -0.29 | -0.16 | -0.49 | -0.27 | -0.13 | -0.36 | -0.43 | 0.07 | 0.10 |
| $\lambda_{abs}^{off}$ | -0.07 | -0.15 | -0.58 | -0.07 | -0.12 | -0.61 | -0.53 | -0.55 | -0.18 | 0.37 | -0.27 | -0.28 | -0.31 | 0.23 | -0.65 | 0.11 | 0.16 | -0.45 | -0.75 |
| $A_{res}$ | -0.02 | -0.34 | -0.24 | 0.02 | -0.19 | -0.07 | -0.22 | 0.01 | -0.35 | -0.26 | 0.01 | 0.15 | -0.34 | 0.10 | -0.28 | -0.11 | -0.05 | -0.44 | -0.42 |
| $A_{res,\infty}$ | 0.03 | -0.30 | -0.19 | 0.08 | -0.16 | -0.06 | -0.17 | 0.05 | -0.31 | -0.27 | 0.04 | 0.16 | -0.35 | 0.12 | -0.23 | -0.06 | -0.02 | -0.42 | -0.39 |
| $\varepsilon_{off}$ | 0.05 | -0.19 | 0.29 | 0.22 | 0.04 | 0.17 | 0.20 | 0.24 | -0.19 | -0.58 | 0.20 | 0.08 | -0.18 | 0.02 | 0.25 | 0.02 | 0.06 | -0.25 | -0.04 |
| $pKa_{app}$ | 0.07 | -0.16 | -0.01 | -0.04 | -0.32 | 0.10 | -0.06 | 0.21 | -0.26 | -0.54 | -0.11 | -0.06 | -0.37 | -0.17 | 0.00 | -0.23 | -0.35 | -0.12 | -0.13 |
| $A_{unrecov}$ | 0.18 | 0.08 | 0.17 | -0.04 | -0.29 | 0.18 | 0.11 | 0.27 | -0.25 | -0.53 | -0.05 | -0.05 | -0.26 | -0.20 | 0.22 | -0.19 | -0.40 | 0.10 | 0.17 |
| $k_{therm}$ | -0.17 | -0.37 | -0.45 | -0.27 | -0.37 | -0.19 | -0.42 | -0.22 | -0.51 | -0.19 | -0.28 | 0.06 | -0.37 | 0.01 | -0.47 | -0.28 | -0.24 | -0.49 | -0.45 |
| $\varepsilon_{anionic}$ | -0.58 | -0.69 | 0.43 | 0.09 | 0.42 | 0.40 | 0.23 | 0.27 | 0.25 | -0.43 | 0.15 | -0.02 | 0.42 | -0.30 | 0.28 | -0.20 | 0.19 | -0.10 | 0.13 |
| $Q_{fluo}$ | -0.06 | 0.00 | 0.32 | 0.33 | 0.41 | 0.17 | 0.36 | 0.09 | 0.05 | 0.05 | 0.04 | 0.30 | -0.01 | 0.24 | 0.26 | 0.27 | 0.38 | -0.41 | 0.22 |
| M. br | -0.27 | -0.10 | 0.25 | 0.21 | 0.48 | 0.11 | 0.26 | -0.06 | 0.16 | 0.16 | 0.05 | 0.19 | 0.27 | 0.18 | 0.18 | 0.23 | 0.47 | -0.32 | 0.16 |
| $\varepsilon_{overall}$ | -0.34 | -0.15 | 0.03 | -0.07 | 0.27 | 0.00 | -0.01 | -0.17 | 0.22 | 0.19 | 0.00 | -0.07 | 0.38 | -0.03 | -0.03 | 0.02 | 0.27 | -0.01 | -0.01 |
| $Q_{off}$ | 0.41 | 0.62 | 0.04 | -0.10 | -0.20 | -0.09 | 0.02 | -0.04 | 0.15 | 0.23 | -0.04 | -0.08 | 0.10 | -0.08 | 0.20 | 0.10 | -0.20 | 0.62 | 0.33 |
| $\lambda_{ex} - \lambda_{abs}$ | 0.25 | 0.43 | -0.15 | -0.31 | -0.28 | -0.10 | -0.09 | -0.13 | -0.06 | 0.25 | -0.19 | 0.08 | -0.04 | -0.13 | -0.01 | -0.11 | -0.30 | 0.41 | 0.30 |
| $\delta_{Stokes}$ | 0.48 | 0.31 | -0.17 | 0.13 | -0.19 | -0.24 | -0.07 | -0.04 | -0.14 | 0.09 | 0.25 | -0.01 | -0.20 | 0.27 | -0.12 | 0.22 | 0.03 | 0.05 | -0.23 |
| $\Delta pH$ | -0.49 | -0.69 | -0.45 | -0.29 | -0.24 | -0.12 | -0.49 | -0.20 | -0.32 | -0.21 | -0.08 | -0.13 | 0.08 | -0.15 | -0.49 | -0.35 | -0.20 | -0.26 | -0.49 |
| $Q_{on}$ | -0.35 | -0.34 | -0.12 | -0.63 | -0.29 | 0.17 | -0.16 | 0.05 | -0.17 | -0.36 | -0.22 | -0.15 | 0.03 | -0.69 | -0.13 | -0.75 | -0.62 | 0.42 | 0.25 |

M. br: Molecular brightness; $\Delta pH$: $\lambda_{abs,pH10} - \lambda_{abs,pH6.5}$.

**Table S10:** Partial least-squares discriminant analysis on spectroscopic and off-state structural properties indicating the FP's structural properties compared to average.

| | **Fast switchers↑[†], Slow switchers↓** | **Fast switchers↓, Slow switchers↑** |
|---|---|---|
| Chromophore descriptors | Tilt | Twist |
| Chromophore and environment based descriptors | Total No. of heteroatoms around chromophore*<br>Chromophore pocket volume<br>**Hydroxybenz$_{OH}$ – N205$_{CA}$**[§]<br>**Hydroxybenz$_{OH}$ – N205$_{OD1/ND2}$**$^{\$}$<br>Hydroxybenz$_{plane}$ – T62$_{CG2}$ | Total No. of atoms around Imidazolinone$_{plane}$<br>Hydroxybenz$_{plane}$ – H148$_{plane}$ |
| Chromophore independent descriptors | **A150$_{CB}$ – L69$_{CD2}$**$^{\$}$<br>F165$_{CB}$ – H148$_{CA}$<br>F165$_{CB}$ – L201$_{CD1}$<br>H148$_{CA}$ – Q94$_{NE2}$<br>H148$_{CA}$ – R96$_{NH2}$<br>H148$_{CA}$ – L69$_{CD2}$<br>**N205$_{CA}$ – F165$_{CB}$**<br>N205$_{CA}$ – I167$_{CD1}$<br>N205$_{CA}$ – L69$_{CD2}$<br>N205$_{CA}$ – Q94$_{NE2}$<br>N205$_{CA}$ – R96$_{NH2}$<br>I167$_{CD1}$ – H148$_{CA}$<br>I167$_{CD1}$ – T203$_{CB}$$^{\$}$<br>T203$_{CB}$ – L69$_{CD2}$ | **Q94$_{NE2}$ – R96$_{NH2}$**<br>F145$_{CA}$ – A150$_{CB}$<br>F145$_{CA}$ – E222$_{CD}$<br>F145$_{CA}$ – L69$_{CD2}$<br>**F145$_{CA}$ – L201$_{CD1}$**<br>F145$_{CA}$ – Q94$_{NE2}$<br>F145$_{CA}$ – T203$_{CB}$ |

↑: Values for parameters in column are higher than average, ↓: Values for parameters in column are lower than average.

[†] Fast switchers: LV1+, LV2-, Slow switchers: LV1-, LV2+.

* Total No. of atoms within a sphere of 3.5 Å.

[§] Descriptors in bold are within the 15 highest loadings of LV1 and LV2 (Supplementary Table S8).

$^{\$}$ higher (lower) than average or equal to average.

**Table S11:** Summary of partial least-square analyses.

**(a)** PLS-DA of the spectroscopic parameters with PFs classified according to zone 1, zone 2 and zone 3

| Latent variable (LV) | X-Block LV | X-Block Cumulative LV | Y-Block LV | Y-Block Cumulative LV | CV Error Class 1 | CV Error Class 2 | CV Error Class 3 |
|---|---|---|---|---|---|---|---|
| **1***| **31.48** | **31.48** | **40.95** | **40.95** | **0.154** | **0.333** | **0.519** |
| **2** | **26.95** | **58.44** | **15.75** | **56.7** | **0.125** | **0.292** | **0.294** |
| **3** | **9.5** | **67.94** | **10.57** | **67.27** | **0.125** | **0.292** | **0.425** |
| **4** | **5.47** | **73.41** | **9.51** | **76.78** | **0.125** | **0.292** | **0.425** |
| **5** | **4.19** | **77.6** | **6.79** | **83.57** | **0.125** | **0.194** | **0.294** |
| **6** | **8.41** | **86.01** | **1.48** | **85.05** | **0** | **0.194** | **0.294** |
| **7** | **5.29** | **91.3** | **1.17** | **86.22** | **0** | **0.153** | **0.263** |
| 8 | 2.68 | 93.98 | 1.17 | 87.39 | 0 | 0.153 | 0.263 |
| 9 | 1.83 | 95.81 | 0.48 | 87.87 | 0 | 0.153 | 0.263 |
| 10 | 2.4 | 98.21 | 0.15 | 88.02 | 0 | 0.208 | 0.362 |

* Bold: LV's used during analysis.

**(b)** PLS-DA of the spectroscopic parameters with PFs classified according to zone 1 and zone 2

| Latent variable (LV) | X-Block LV | X-Block Cumulative LV | Y-Block LV | Y-Block Cumulative LV | Average CV Class Error |
|---|---|---|---|---|---|
| **1*** | **33.13** | **33.13** | **86.29** | **86.29** | **0** |
| 2 | 21.38 | 54.5 | 8.49 | 94.78 | 0 |
| 3 | 12.61 | 67.12 | 3.41 | 98.19 | 0.063 |
| 4 | 10.12 | 77.24 | 0.81 | 99 | 0.063 |
| 5 | 10.7 | 87.94 | 0.16 | 99.16 | 0.125 |
| 6 | 3.99 | 91.93 | 0.18 | 99.33 | 0.125 |
| 7 | 2.01 | 93.94 | 0.38 | 99.71 | 0.188 |
| 8 | 2.88 | 96.82 | 0.12 | 99.83 | 0.188 |
| 9 | 0.77 | 97.59 | 0.11 | 99.95 | 0.25 |
| 10 | 0.86 | 98.45 | 0.02 | 99.96 | 0.271 |

* Bold: LV's used during analysis.

**(c)** PLS-DA of the spectroscopic and off-state parameters with PFs classified according to good, average and bad switchers.

| Latent variable (LV) | X-Block LV | X-Block Cumulative LV | Y-Block LV | Y-Block Cumulative LV | CV Error Class 1 | CV Error Class 2 | CV Error Class 3 |
|---|---|---|---|---|---|---|---|
| **1*** | **20.55** | **20.55** | **41.86** | **41.86** | **0.417** | **0.134** | **0.25** |
| **2** | **15.24** | **35.79** | **39.23** | **81.09** | **0.208** | **0.134** | **0.2** |
| **3** | **12.71** | **48.5** | **6.01** | **87.1** | **0.083** | **0.188** | **0.25** |
| **4** | **8.11** | **56.61** | **5.04** | **92.14** | **0.25** | **0.125** | **0.1** |
| **5** | **6.91** | **63.51** | **3.7** | **95.84** | **0.083** | **0.125** | **0.1** |
| 6 | 8.92 | 72.43 | 2.08 | 97.92 | 0.125 | 0.196 | 0.15 |
| 7 | 7.94 | 80.38 | 0.97 | 98.89 | 0.25 | 0.321 | 0.15 |
| 8 | 5.64 | 86.02 | 0.52 | 99.41 | 0.292 | 0.25 | 0.25 |
| 9 | 2.82 | 88.83 | 0.32 | 99.73 | 0.208 | 0.134 | 0.15 |
| 10 | 2.29 | 91.12 | 0.14 | 99.87 | 0.542 | 0.589 | 0.45 |

* Bold: LV's used during analysis.

**Table S12:** List of primers.

| Mutation | Sequence |
|---|---|
| rsGreen0.7/rsGreen1/rsGreenF-**K206A** | 5’-GTC TTT GCT CAG CGC ATT CTG GGT GCT CAG GTA GTG G |
| rsGreen0.7-(K206A)-**F145X** | 5’-CAC AAG CTG GAG TAC AAC NNK AAC AGC CAC AAC GCC TAT A |
| rsGreen0.7-(K206A)-**F145L** | 5’-GTT GTG GCT GTT TAA GTT GTA CTC CAG CTT GTG CC |
| rsGreen0.7-(K206A)-**E222X** | 5’-CAC ATG GTC CTG CTG NNK TTC GTG ACC GCC GC |
| rsGreen0.7-(K206A)-**F165X** | 5’-ACG GCA TCA AGT CTA ACN NKA AGA TCC GCC ACA ACG TC |
| rsGreen0.7-(K206A)-**F165R/N/C/N/H/I/L/K/M/F/S/W/Y** | 5’-ACG GCA TCA AGT CTA ACH DSA AGA TCC GCC ACA ACG TC |
| rsGreen0.7-K206A-**N205X** | 5’-ACT ACC TGA GCA CCC AGN NKG CGC TGA GCA AAG ACC C |
| rsGreen0.7-K206A-**N205A/C/I/L/F/P/S/T** | 5’-ACT ACC TGA GCA CCC AGH BTG CGC TGA GCA AAG ACC C |
| rsGreen0.7-(K206A)-**H148X** | 5’-GGA GTA CAA CTT CAA CAG CNN KAA CGC CTA TAT CAC GGC CG |

**Table S13:** Crystallization conditions and cryoprotectants.

| Protein | State | Buffer* | Protein concentration | Crystallization cocktail | Cryoprotectant |
|---|---|---|---|---|---|
| **rsGreenF** | On | HN 50/30 | 10 mg/ml | 100 mM MIB pH 4.0<br>25 % PEG 1500 | 20 % PEG 400 |
| | Off | HN 50/30 | 10 mg/ml | 100 mM MIB pH 4.0<br>25 % PEG 1500 | 20 % PEG 400 |
| **rsGreen1** | On | TN 10/30 | 10 mg/ml | 2.0 M $(NH_4)_2SO_4$<br>200 mM NaCl<br>100 mM Na-cacodylate pH 6.5 | 10 % Glycerol |
| **rsEGFP** | On | H 50 | 10 mg/ml | 200 mM $(NH_4)_2SO_4$<br>100 mM Bis-trip pH 5.5<br>25 % PEG 3350 | 20 % PEG 400 |
| | Off | H 50 | 10 mg/ml | 2.0 M $(NH_4)_2SO_4$<br>200 mM K/Na-tartrate<br>100 mM citrate pH 5.0<br>100 mM b-Nicotinamide adenine dinucleotide | 20 % PEG 400 |
| **rsGreen0.7b** | On | H 50 | 10 mg/ml | 150 mM KBr<br>30 % PEG 2000 mme | 20 % PEG 400 |
| | Off | HN 50/30 | 10 mg/ml | 100 mM MMT pH 5.0<br>25 % PEG 1500 | 10 % Glycerol |

* H 50 : 50 mM HEPES pH7.4; HN 50/30: 50mM HEPES, 30 mM NaCl pH 7.4; TN 10/30: 10 mM Tris, 30 mM NaCl pH 7.4.

*(Continued)* Crystallization conditions and cryoprotectants.

| Protein | State | Buffer* | Protein concentration | Crystallization cocktail | Cryoprotectant |
|---|---|---|---|---|---|
| **rsGreen0.7-F145Q** | On | TN 10/30 | 10 mg/ml | 200 mM $KNO_3$<br>20 % PEG 3350 | 20 % PEG 400 |
| **rsGreen0.7-F145M** | On | TN 10/30 | 5 mg/ml | 200 mM $NH_4HCO_2$<br>20 % PEG 4000 | 20 % PEG 400 |
| | Off | TN 10/30 | 10 mg/ml | 100 mM MMT pH 8.0<br>25 % PEG 1500 | 10 % glycerol |
| **rsGreenF-K206A** | On | HN 50/30 | 10 mg/ml | 200 mM NaCl<br>100 mM MES pH 6.0<br>20 % PEG 6000 | 10 % Glycerol |
| | Off | HN 50/30 | 10 mg/ml | 200 mM NaCl<br>100 mM MES pH 6.0<br>20 % PEG 6000 | 10 % Glycerol |
| **rsGreen1-K206A** | Off | HN 50/30 | 10 mg/ml | 100 mM Bis-tris pH 5.5<br>200 mM $(NH_4)_2SO_4$<br>16 % PEG 8000 | 10 % Glycerol |
| **rsGreen0.7-K206A** | On | TN 10/30 | 10 mg/ml | 200 mM Ammonium acetate<br>100 mM Bis-tris pH 5.5<br>25 % PEG 3350 | 10 % glycerol |
| | Off | HN 50/30 | 10 mg/ml | 22 % PEG 4000<br>140 mM $Mg(NO_3)_2$ | 10 % Glycerol |
| **rsGreen0.7-K206A-N205S** | On | TN 10/30 | 10 mg/ml | 200 mM Maic acid pH 7.0<br>20 % PEG 3350 | 20 % PEG 400 |

* H 50 : 50 mM HEPES pH7.4; HN 50/30: 50mM HEPES, 30 mM NaCl pH 7.4; TN 10/30: 10 mM Tris, 30 mM NaCl pH 7.4.

*(Continued)* Crystallization conditions and cryoprotectants.

| Protein | State | Buffer* | Protein concentra- tion | Crystallization cocktail | Cryoprotectant |
|---|---|---|---|---|---|
| **rsGreen0.7-K206A-N205G** | On | TN 10/30 | 10 mg/ml | 100 mM MES pH 6.0<br>20 % PEG 2000 mme<br>10 mM $BaCl_2$ | 10 % Glycerol |
| **rsGreen0.7-K206A-N205C** | On | HN 50/30 | 10 mg/ml | 100 mM MES pH 6.5<br>25 % PEG 3000 | 10 % glycerol |
| **rsGreen0.7-K206A-N205L** | On | HN 50/30 | 10 mg/ml | 100 mM HEPES pH 7.5<br>25 % PEG 8000 | 20 % PEG 400 |
| **rsGreen0.7-K206A-H148V** | On# | TN 10/30 | 10 mg/ml | 100 mM MES pH 6.5<br>25 % PEG 550 mme<br>10 mM $ZnSO_4$ | 10 % Glycerol |
| | Off | TN 10/30 | 10 mg/ml | 50 mM HEPES pH 7.5<br>12 % PEG 4000<br>1% tryptone | 10 % Glycerol |
| **rsGreen0.7-K206A-H148G** | On | HN 50/30 | 10 mg/ml | 200 mM TMAO<br>100mM Tris pH 8.5<br>20 % Peg 2000 mme | 20 % PEG 400 |
| **rsGreen0.7-K206A-H148S** | On | HN 50/30 | 10 mg/ml | 100 mM Bis-tris pH 6.5<br>20 % PEG 5000 mme | 20 % PEG 400 |
| | Off | HN 50/30 | 10 mg/ml | 100 mM Bis-tris pH 6.5<br>20 % PEG 5000 mme | 20 % PEG 400 |

* H 50 : 50 mM HEPES pH7.4; HN 50/30: 50mM HEPES, 30 mM NaCl pH 7.4; TN 10/30: 10 mM Tris, 30 mM NaCl pH 7.4.
### crystals grown at 4°C.

*(Continued)* Crystallization conditions and cryoprotectants.

| Protein | State | Buffer* | Protein concentration | Crystallization cocktail | Cryoprotectant |
|---|---|---|---|---|---|
| **rsGreen0.7-K206A-F145H** | On | HN 50/30 | 10 mg/ml | 125 mM MES pH 6.0<br>9 % PEG 4000 | 20 % PEG 400 |
| | Off | HN 50/30 | 10 mg/ml | 100 mM MES pH 6.0<br>14 % PEG 400<br>300 mM NGSB-195 | 10 % glycerol |
| **rsGreen0.7-K206A-F145Q** | On | TN 10/30 | 10 mg/ml | 22 % PEG 4000<br>140 mM $Mg(NO_3)_2$ | 20 % PEG 400 |
| | Off | TN 10/30 | 10 mg/ml | 200 mM NaCl<br>100 mM MES pH 6.0<br>20 % PEG 6000 | 10 % glycerol |
| **rsGreen0.7-K206A-F145A** | On | TN 10/30 | 10 mg/ml | 200 mM NaCl<br>100 mM MES pH 6.0<br>20 % PEG 6000<br>4% $C_2H_3F_3O$ | 10 % glycerol |
| | Off# | TN 10/30 | 10 mg/ml | 200 mM NaCl<br>100 mM MES pH 6.0<br>20 % PEG 6000<br>200 mM NDSB-211 | 10 % glycerol |
| **rsGreen0.7-K206A-F145L** | On | TN 10/30 | 10 mg/ml | 100 mM MES pH 6.5<br>12 % PEG 20 000<br>10 mM NaBr | 20 % PEG 400 |
| | Off | TN 10/30 | 10 mg/ml | 100 mM MES pH 6.5<br>12 % PEG 20 000<br>4 % Pentaerythritol ethoxylate | 20 % PEG 400 |

* H 50 : 50 mM HEPES pH7.4; HN 50/30: 50mM HEPES, 30 mM NaCl pH 7.4; TN 10/30: 10 mM Tris, 30 mM NaCl pH 7.4.
### crystals grown at 4°C.

*(Continued)* Crystallization conditions and cryoprotectants.

| Protein | State | Buffer* | Protein concentration | Crystallization cocktail | Cryoprotectant |
|---|---|---|---|---|---|
| **rsGreen0.7-K206A-F145S** | On | HN 50/30 | 10 mg/ml | 100 mM MES pH 6.0<br>14 % PEG 4000 | 20 % PEG 400 |
| | Off | TN 10/30 | 10 mg/ml | 100 mM MES pH 6.5<br>25 % PEG 4000<br>400 mM D-galactose | 20 % PEG 400 |
| **rsGreen0.7-K206A-F145M** | On | TN 10/30 | 5 mg/ml | 100 mM Bis-tris pH 6.5<br>20 % PEG 1500 | 20 % PEG 400 |
| | Off | TN 10/30 | 5 mg/ml | 100 mM MMT<br>25 % PEG 1500 | 20 % PEG 400 |
| **rsGreen0.7-K206A-F165L** | On | HN 50/30 | 10 mg/ml | 100 mM HEPES pH 7.5<br>25 % PEG 4000 | 10 % Glycerol |
| **rsGreen0.7-K206A-F165W** | On | HN 50/30 | 10 mg/ml | 200 mM $NH_4F$<br>20 % PEG 3350 | 10 % glycerol |
| | Off | HN 50/30 | 10 mg/ml | 100 mM HEPES pH 7.5<br>25 % PEG 4000 | 20 % PEG 400 |
| **rsGreen0.7-K206A-E222G** | On | TN 10/30 | 10 mg/ml | 200 mM $(Li)_2SO_4$<br>100 mM Tris pH 8.5<br>40 % PEG 400 | 10 % glycerol |
| **rsGreen0.7-K206A-E222V** | On | HN 50/30 | 10 mg/ml | 26 % PEG 4000<br>500 mM $Mg(NO_3)_2$<br>50 mM HEPES pH 7.5 | 20 % PEG 400 |

* H 50 : 50 mM HEPES pH7.4; HN 50/30: 50mM HEPES, 30 mM NaCl pH 7.4; TN 10/30: 10 mM Tris, 30 mM NaCl pH 7.4.
